# Mitochondrial ATP production promotes T cell differentiation and function by regulating chromatin accessibility

**DOI:** 10.64898/2026.03.27.714789

**Authors:** Charles Ng, Tak Shun Fung, Dayi Li, Korbinian N Kropp, Luis F Somarribas Patterson, Alexandria Markovitz, Daniel N Weinberg, Olivia Jones, Ji-Young Kim, Guoan Zhang, Richard Koche, Mara Monetti, Huayuan Tang, Yun He, Zhengshuang Xu, Xin Cai, Ziqi Yu, Geetha Bhagavatula, Sean P Colgan, Ya-Hui Lin, Zhuoning Li, Elizabeth M Steinert, Christopher A Klebanoff, Santosha A Vardhana, Navdeep S Chandel, Lin Wu, Craig B Thompson

## Abstract

Immune elimination of chronic infection or cancer requires cytotoxic CD8^+^ T cells that adopt and maintain an effector phenotype. Cytotoxic T cell function is a bioenergetically demanding process and T cells subjected to chronic antigen exposure have compromised effector function despite high rates of glycolysis. Here we report the ability of the short-chain α-hydroxy acid, D-α-hydroxybutyrate, to act as a signaling molecule that increases mitochondrial ATP production and drives the conversion of proliferating T cells into cytotoxic effector cells. DAHB signaling switches ATP production from glycolysis to oxidative phosphorylation supported by fatty acid oxidation, even in glucose-replete media. This conversion suppresses both AMPK phosphorylation and the integrated stress response (ISR) in activated T cells while significantly elevating the level of the phosphagen, phosphocreatine (PCr). Both the PCr bioenergetic reserve and oxidative phosphorylation were required for T cell effector differentiation. DAHB-induction of CD8-effector gene transcription was coupled to bioenergetics by enhanced ATP-dependent remodeling of chromatin accessibility at effector gene loci. DAHB enhanced CD8^+^ T cell antitumor activity both *in vitro* and *in vivo,* and DAHB treatment of transferred T cells led to persistent *in vivo* antitumor effects. Together, these findings link cellular bioenergetics to the regulation of chromatin accessibility and gene expression required to support effector function.

## Introduction

Activated T cells undergoing proliferative expansion in secondary lymphoid organs produce ATP primarily through aerobic glycolysis. Glycolytic ATP production can be sufficient to support T cell proliferation and TCR-dependent kinase signaling, but concomitant upregulation of mitochondrial oxidative phosphorylation (OXPHOS) is required for optimal T cell effector function^1,2^. T cells with mutations in TCA cycle components or mtDNA are compromised when responding to microbial pathogens and tumors^3–5^. Exhaustion of effector T cells at peripheral sites of infection or within the tumor microenvironment is linked to mitochondrial dysfunction and compromised OXPHOS^6,7^. Despite an elevated glycolytic rate, exhausted T cells have depleted levels of ATP and reduced expression of effector genes^8^.

The extracellular microenvironment can also impose metabolic challenges to cytotoxic T effector cells. Extracellular depletion of glucose and amino acids activates energy-sensing and cytoprotective pathways such as AMP-activated protein kinase (AMPK) and the integrated stress response (ISR), which act as adaptive brakes on T cell function. AMPK, triggered by low ATP/AMP ratios, downregulates mTORC1 signaling and protein synthesis, curbing cytokine production under glucose-limited conditions^9^. The ISR similarly dampens biosynthetic output by repressing cap-dependent translation, while increasing transcription and translation of a subset of stress-response genes mediated by the transcription factor ATF4^10^. Although these adaptive mechanisms preserve cell viability during metabolic stress, they restrict the full potential of T cell effector function and contribute to T cell exhaustion.

T cells can use oxidizable substrates to sustain their survival and function when their bioenergetic needs are unmet by aerobic glycolysis. Butyrate and acetate have both been shown to support T cell function through their ability to be metabolized through oxidative phosphorylation when glucose is limiting^11,12^. While T cell effector function can be maintained by increasing the availability of oxidizable substrates that support mitochondrial electron transport chain (ETC) activity, this is not sufficient to convert T cells into better effectors^13^. These treatments can also induce redox stress that limits T cell effector function and contributes to the development of T cell dysfunction. In addition to their role as bioenergetic substrates, metabolites can also act as signaling molecules to allosterically modulate cellular processes. For example, L-serine activates pyruvate kinase to enhance glycolytic ATP production, mitochondrial accumulation of N-acetyl-glutamate activates carbamoyl phosphate synthase to increase ureagenesis, and L/D-2-hydroxyglutarate inhibits demethylases^14–16^.

Recently, we reported that D-lactate could act as a signaling molecule to stimulate mitochondrial oxidative phosphorylation without being oxidized or causing mitochondrial reductive stress^17^. D-lactate is known to accumulate systemically to millimolar levels in patients with small bowel dysbiosis, consistent with its primary origin as a microbial metabolite^18,19^. D-lactate can also be produced by degradation of methylglyoxal, which accumulates in metabolic and neurodegenerative disorders^20^. The accumulation of D-lactate has been associated with neurologic side effects, which could limit its study as a metabolic regulator *in vivo*, prompting us to test structurally related molecules^21^. The closest structurally related α-hydroxy acid to D-lactate is D-α-hydroxybutyrate (DAHB). Many bacteria produce short-chain D-α-hydroxy acids as fermentation end products^22^. Recently, *Fusobacterium nucleatum*, a Gram-negative pathobiont with enhanced capacity to produce DAHB during fermentation compared with other Gram-negative bacteria, was reported to be associated with the success of immunotherapy in the treatment of colorectal cancer^23–27^. This prompted us to investigate the effects of DAHB on T cell differentiation and effector function.

Like D-lactate, DAHB acts as an activator of ETC activity without being a meaningful oxidizable substrate. We found that DAHB-stimulated effector T cells used beta-oxidation to support mitochondrial ATP production even when glucose was abundant. Glycolytic carbon was directed into supporting pentose phosphate pathway activity, amino acid biosynthesis and protein glycosylation. DAHB treatment led to increased cellular bioenergetic reserves and preserved effector gene expression during chronic antigen stimulation. DAHB-treated T cells increased cellular phosphocreatine (PCr) levels and this increase was accompanied by increased T cell effector gene expression along with increased expression of genes involved in mitochondria-to-nuclear PCr shuttling. DAHB treatment increased chromatin accessibility at T cell effector loci through the ability of elevated PCr levels to buffer ATP while limiting stress-induced AMPK activation. DAHB-induced chromatin accessibility also required the presence of functional ATP-dependent chromatin remodeling complexes. These studies provide evidence that OXPHOS-derived ATP production is coupled to the regulation of nuclear gene expression, in part through maintaining chromatin accessibility at lineage-specific effector loci. Functionally, DAHB was well tolerated and enhanced the efficacy of *in vivo* immunotherapy using either murine or human T cells.

## Results

### DAHB is a potent inducer of T cell differentiation and function

To explore whether a D-α-hydroxy acid structurally related to D-lactate and known to be produced during bacterial metabolism could enhance T cell effector function, we tested DAHB for the ability to induce CD8^+^ T cell cytokine production in cells activated for 48 hours with anti-CD3 and anti-CD28 (Figure 1A and Figure S1A). DAHB treatment demonstrated an even more potent enhancement of both interferon gamma (IFN-γ) and Perforin than D-lactate when compared to controls as measured 24 hours after addition (Figure 1B). This effect was most striking for the cytolytic effector molecule Perforin. D-lactate stimulated less than half of the cells to produce Perforin. In contrast, nearly all cells treated with DAHB showed increased expression including per-cell levels as measured by MFI (Figures S1B and S1C). Genome-wide analysis with RNA-seq and proteomics revealed that even when DAHB was added 24 hours after initial activation, DAHB-mediated upregulation of numerous additional effector molecules including TNF family members, chemokines and cytokines, compared to cells activated without DAHB treatment (Figures 1C and 1D). To investigate the specificity of the response, we also tested L-α-hydroxybutyrate (LAHB), L-β-hydroxybutyrate (LBHB), and D-β-hydroxybutyrate (DBHB) at equimolar doses and compared to the maximal tolerated dose (0.5 mM) of butyrate (Figures 1E, S1D and S1E). DAHB and butyrate were the most potent cytokine inducers (Figure 1F). The effects of DAHB were dose-dependent between 2–20 mM and washout of DAHB followed by culture in control medium for 24 hours showed persistent enhancement of IFN-γ and Perforin expression (Figures S1F and S1G). To confirm the functional impact of DAHB, we pretreated antigen-specific OT-I CD8^+^ T cells overnight followed by incubation with the OVA-expressing lymphoma cell line EL4-OVA. Compared to control treatment, DAHB showed an approximate threefold increase in T cell cytotoxicity, and this enhancement was observed across different effector-to-target cell ratios (E:T) (Figures 1G and S1H).

**Figure 1.**
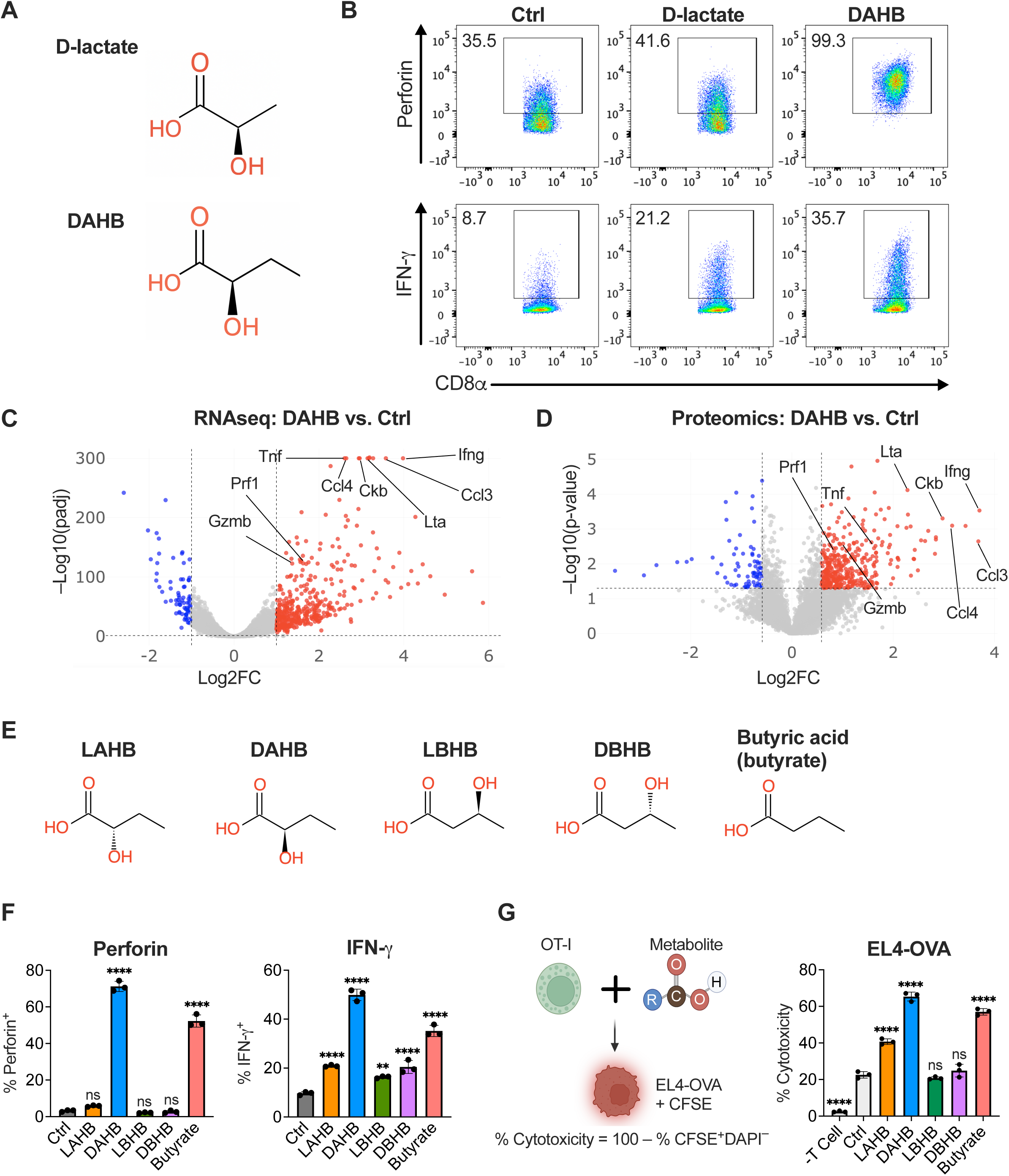
**DAHB promotes CD8^+^ T cell cytokine production and tumor cytotoxicity.** (A) Chemical structures of D-lactate and DAHB. (B) Flow cytometry of Perforin and IFN-γ production in CD8^+^ T cells activated with anti-CD3/CD28 for 48 h, followed by 24 h of treatment with NaCl (Ctrl, 20 mM), D-lactate (20 mM), or DAHB (20 mM). Data are representative of three independent experiments. Numbers indicate percent of cells in gates. (C) RNA-seq volcano plot showing differential gene expression in CD8^+^ T cells activated for 24 h with anti-CD3/CD28 followed by 24 h of treatment with DAHB or Ctrl; n = 3. (D) Proteomics volcano plot showing differential protein abundance in CD8^+^ T cells from the same experiment as (C); n = 3. (E) Chemical structures of L/D-α-hydroxybutyrate (LAHB and DAHB), L/D-β-hydroxybutyrate (LBHB and DBHB), and butyrate. (F) Perforin and IFN-γ production by flow cytometry in activated T cells after 24 h of treatment with Ctrl (20 mM), LAHB (20 mM), DAHB (20 mM), LBHB (20 mM), DBHB (20 mM), or butyrate (0.5 mM); n = 3. (G) OT-I CD8^+^ T cells treated with indicated metabolites as in (F) followed by washout and overnight co-culture of CFSE-labeled EL4-OVA tumor cells at an 8:1 effector-to-target (E:T) ratio with −T cell control included. Percent cytotoxicity was measured as 100 − %CFSE^+^DAPI^−^ cells; n = 3. Data are mean ± SD. Differential expression in (C) was determined by DESeq2 with Benjamini-Hochberg correction (adjusted p < 0.05 and fold change > 2). Differential protein abundance in (D) was determined by Welch’s t test (p < 0.05 and fold change > 1.5). Statistical analysis in (F, G) was performed by one-way ANOVA with Dunnett’s multiple comparisons test relative to Ctrl. ns, not significant; *p < 0.05; **p < 0.01; ***p < 0.001; ****p < 0.0001. See also Figure S1.

### DAHB induces OXPHOS and FAO independently of its own metabolism

To study if DAHB could enhance mitochondrial oxidative phosphorylation like D-lactate^17^, we next examined the effects of DAHB on activated T cells using a Seahorse analyzer. When butyrate, a known mitochondrial substrate for oxidation, was added to activated CD8^+^ T cells, both their oxygen consumption and spare respiratory capacity increased^11^. In contrast, DAHB addition to activated CD8^+^ T cells led to an increase in ATP-coupled oxygen consumption, but the spare respiratory capacity was unchanged, suggesting DAHB acts as a signaling molecule rather than a mitochondrial substrate in stimulating ETC-dependent mitochondrial ATP production (Figure 2A). DAHB also significantly reduced glycolysis as measured by both the extracellular acidification rate (ECAR) and the consumption of glucose from the medium (Figures 2B and 2C). The accumulation of lactate in the medium was also significantly reduced.

**Figure 2.**
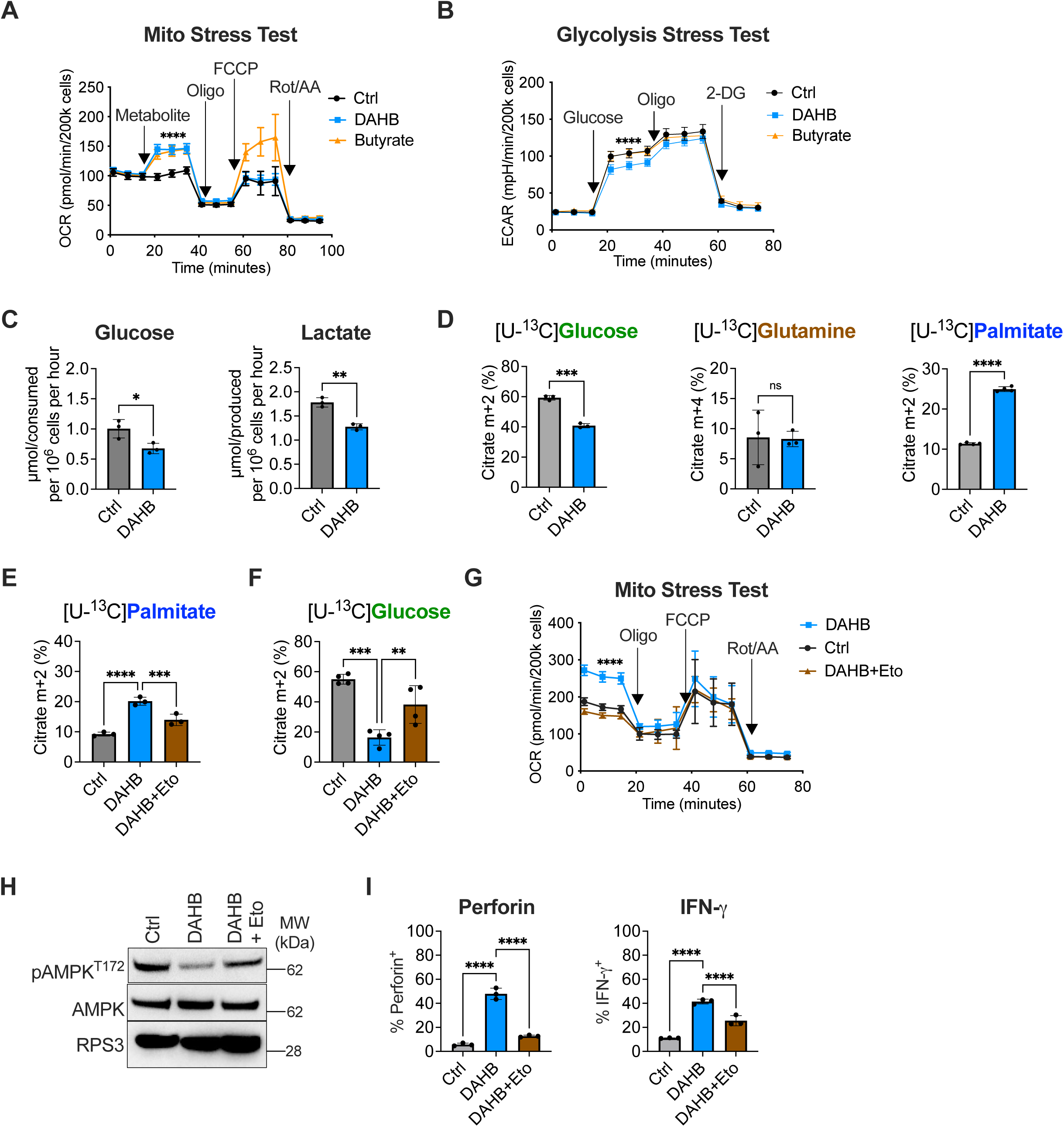
**DAHB signaling induces fatty acid oxidation (FAO) to enhance bioenergetics.** (A) Oxygen consumption rate (OCR) measured by Seahorse analysis in activated CD8^+^ T cells following acute injection of NaCl (Ctrl, 20 mM), DAHB (20 mM), or butyrate (0.5 mM) (n = 8 wells per condition). Oligomycin (Oligo), FCCP, and rotenone + antimycin A (Rot/AA) were injected sequentially from ports B–D. (B) Extracellular acidification rate (ECAR) measured by Seahorse analysis in activated CD8^+^ T cells treated with Ctrl, DAHB or butyrate at concentrations in (A) (n = 8 wells per condition). Glucose, Oligo, and 2-deoxyglucose (2-DG) were injected sequentially from ports A–C. (C) Glucose consumption and lactate accumulation in the culture medium of CD8^+^ T cells treated with Ctrl or DAHB for 24 h and normalized to cell number as measured by a YSI analyzer; n = 3. (D) Stable isotope tracing showing citrate mass isotopologue enrichment following labeling with [U-^13^C]glucose (30 min) or [U-^13^C]glutamine and [U-^13^C]palmitate (5 h) in Ctrl vs. DAHB-treated CD8^+^ T cells. Glucose and glutamine data; n = 3. Palmitate data pooled from two experiments performed in technical duplicate; n = 4. (E) Effect of CPT1A inhibition by etomoxir (Eto, 80 µM) on palmitate oxidation after treatment with Ctrl, DAHB, and DAHB + Eto in CD8^+^ T cells; n = 3. (F) Effect of Eto on glucose oxidation after treatment with Ctrl, DAHB, and DAHB + Eto in CD8^+^ T cells as in (E). Data pooled from two experiments performed in technical duplicate; n = 4. (G) OCR measured by Seahorse analysis of activated CD8^+^ T cells treated as in (E) with Ctrl, DAHB, or DAHB + Eto. Treatments were maintained during the assay; n = 9 wells per condition. Oligo, FCCP, and Rot/AA were injected sequentially from ports A–C. (H) Immunoblot of phosphorylated AMP-activated kinase (pAMPK^T172^) and AMPK (total) in Ctrl, DAHB, or DAHB + Eto-treated CD8^+^ T cells. Ribosomal protein S3 (RPS3) serves as a loading control. Representative of two independent experiments. (I) Perforin and IFN-γ production measured by flow cytometry in activated CD8^+^ T cells treated with Ctrl, DAHB, or DAHB + Eto; n = 3. Data are mean ± SD. Statistical analysis in (A, B, E, F, G, I) was performed by one-way ANOVA with Tukey’s multiple comparisons test and in (C, D) by unpaired two-tailed Student’s t test. For (A, B, G), statistical analysis was performed on all three timepoints within the indicated interval. ns, not significant; *p < 0.05; **p < 0.01; ***p < 0.001; ****p < 0.0001. See also Figures S2–S5.

We next tested whether DAHB could sustain elevated OXPHOS over longer time scales. Continuous oximetry revealed that the DAHB-mediated increase in OCR persisted over 24 hours in culture (Figure S2A). Notably, the increased mitochondrial respiration observed over 24 hours occurred without a significant increase in lipid peroxidation (Figure S2B). Even activated T cells cultured for 48 hours in DAHB-containing medium still maintained an increased level of oxidative phosphorylation in comparison to activated T cells cultured in the absence of DAHB suggesting that, in response to chronic ETC activation, they maintained access to substrates that could support OXPHOS (Figures S2C–S2E).

The above results suggested that DAHB stimulation of ETC-coupled ATP production is sustained by cellular metabolic changes that altered the substrate(s) being utilized to support TCA cycle-dependent oxidative phosphorylation. An obvious candidate substrate was DAHB itself. We first examined how DAHB is taken up by T cells. Monocarboxylates, like pyruvate and lactate that can support OXPHOS, enter cells through proton-coupled monocarboxylate transporters (MCTs)^28,29^.

Therefore, we asked whether DAHB also utilized MCTs for entry. Using a cytosolic pH-sensitive dye^30^, we found DAHB rapidly induced intracellular acidification comparable to that observed following addition of pyruvate, L-lactate and D-lactate. This acidification was not observed in control cells or cells treated with nutrients that are taken up in an MCT-independent manner such as alanine and glucose (Figure S2F). This influx of DAHB reached steady state within 30 minutes as did the influx of L-lactate at an equimolar dose (Figure S2G). To confirm DAHB uptake by the MCTs, we pretreated CD8^+^ T cells with the MCT1/4 inhibitor syrosingopine^31^ (Syro) and this blocked lactate- and DAHB-induced acidification (Figure S2H).

To investigate whether the DAHB taken up provides the carbon source for sustained OXPHOS, we used stable isotope tracing with carbon-13 ([U-^13^C]) to test whether DAHB carbon enters the TCA cycle. While [U-^13^C]butyrate readily labeled TCA intermediates, none were labeled by [U-^13^C]DAHB (Figure S3A). Furthermore, the oxidation of DAHB to α-ketobutyrate (AKB), which enters the TCA cycle as a succinyl-CoA^32^, was not detected. The only validated mammalian gene capable of metabolizing D-2-hydroxy acids including DAHB is mitochondrial D-lactate dehydrogenase^33^ (*Ldhd*). Therefore, to test if DAHB catabolism was required for its effect on T cell effector function, we generated a T cell conditional *Ldhd* knockout mouse and confirmed gene deletion at the DNA and RNA levels (*Ldhd*^F/F^;CD4Cre) (Figures S3B–S3D). The DAHB-treated KO CD8^+^ T cells phenocopied the WT DAHB-mediated cytokine and OXPHOS induction, consistent with a role for DAHB as a signaling metabolite rather than through its conversion to a metabolic intermediate (Figures S3E and S3F).

Despite not being a metabolic substrate, DAHB treatment led to significant changes in the substrates supporting the TCA cycle. Although glucose oxidation decreased and glutamine oxidation was unchanged, there was a doubling of fatty acid oxidation (FAO) of the long-chain fatty acid palmitate (Figure 2D). This shift towards fatty acid oxidation was not due to reduced glucose availability. Despite less glucose being oxidized or secreted as lactate, DAHB-treated cells had enhanced glucose flux into anabolic pathways such as Uridine 5′-diphospho-N-acetylglucosamine (UDP-GlcNAc) and serine/glycine biosynthesis (Figures S3G and S3H). DAHB treatment also reduced the total abundance of acetyl-CoA and propionyl-CoA, derived primarily from mitochondrial metabolism of pyruvate and amino acids (Figure S4A).

The significant acetyl-CoA reduction observed in DAHB-treated T cells suggested that the elevation of palmitate oxidation might be indicative of a generalized activation of fatty acid oxidation^34^. To test this, we assayed multiple classes of fatty acids and found oxidation of short-, medium- and long-chain fatty acids were all significantly increased (Figure S4B). To confirm that DAHB-mediated stimulation of OXPHOS is supported by FAO, we treated activated T cells with Etomoxir (Eto) to reduce the CPT1A-dependent entry of long-chain fatty acids (LCFAs) into mitochondria. Treatment with Eto reduced palmitate oxidation (Figure 2E). Eto treatment increased glucose carbon entering the TCA cycle, but neither the DAHB-induced increase in oxidative phosphorylation nor ETC-coupled ATP production was restored (Figures 2F and 2G). A novel feature of the DAHB-induced increase in OXPHOS was also the ability to reduce pAMPK levels and Eto treatment suppressed this effect (Figure 2H). Eto treatment also inhibited the DAHB-induced increase in T cell effector gene expression (Figure 2I).

To independently confirm the role of FAO in DAHB-mediated cytokine induction and mitigate the confounding off-target effects that Eto is reported to have on reducing ETC function^35,36^, we used CRISPR to KO *Cpt1a* (target of Eto) and *Hadhb*, a downstream LCFA thiolase in beta-oxidation (Figures S5A and S5B). CD8^+^ T cells impaired for LCFA oxidation showed partial inhibition of the DAHB-mediated expression of IFN-γ and Perforin (Figures S5C and S5D). We also used an additional pharmacological approach with teglicar (Teg), an analog of L-palmitoylcarnitine and competitive inhibitor of CPT1A^37,38^ (Figure S5E). Teg treatment inhibited DAHB-mediated oxidation of palmitate and activated AMPK (Figures S5F and S5G). Teg treatment reduced DAHB-induced OCR, but this suppression was partly rescued by the CPT1A-independent short-chain fatty acid, butyrate (Figure S5H). Butyrate was also able to significantly rescue the cytokine reduction mediated by Teg treatment of DAHB-stimulated T cells (Figure S5I).

To investigate the possibility that the DAHB-mediated reduction in pAMPK, downstream of FAO, was sufficient for the increased effector gene expression observed, we also targeted the alpha catalytic subunit *Prkaa1* (AMPKα1) with CRISPR (Figure S5J). The AMPKα1 KO failed to increase cytokine production in activated T cells. Furthermore, the AMPKα1 KO reduced the ability of DAHB to increase T cell effector gene expression, consistent with the requirement of basal levels of AMPK signaling to support T cell activation^39,40^ (Figures S5K and S5L). Finally, we tested whether DAHB-mediated metabolic changes could promote T cell effector function under glucose-limiting conditions by culturing DAHB-treated activated T cells in low-glucose media. DAHB treatment significantly increased effector cytokines without altering cell numbers under glucose-limiting conditions (Figure S5M).

### DAHB enhances OXPHOS-dependent gene expression

The effects of DAHB on mitochondrial ETC activity are distinct from butyrate and its other derivatives. However, short-chain fatty acids related to DAHB, including butyrate, are reported histone deacetylase (HDAC) inhibitors. We found that DAHB also inhibited total cellular HDAC activity, including HDAC1/2, although less so than butyrate or the pan-HDAC inhibitor trichostatin A (TSA) (Figures S6A and S6B). Profiling histone acetylation and methylation independently confirmed that DAHB treatment led to increases in acetylation while histone methylation was decreased (Figures S6C and S6D). However, the changes of histone modifications observed with DAHB treatment were not dependent on the DAHB-induced increase in mitochondrial ATP production. Inhibiting DAHB-induced OXPHOS with the ATP synthase inhibitor oligomycin did not block global histone acetylation or locus-specific histone acetylation as determined by ChIP-qPCR, indicating histone acetylation is not sufficient to drive effector gene expression (Figures S6E and S6F).

To explore further how DAHB-induced OXPHOS could be influencing gene expression, RNA-seq was performed on activated CD8^+^ T cells treated with Ctrl, DAHB, or DAHB+oligomycin (DAHB+O). Principal component analysis (PCA) revealed distinct changes in gene expression by DAHB, which were widely affected by oligomycin treatment (Figure S7A). K-means clustering produced 5 clusters of gene expression (Figure S7B). Cluster 1 revealed DAHB-induced genes whose induction was blocked by oligomycin treatment and included numerous effector genes such as *Prf1* and *Ifng* (Figure S7C). In contrast, Cluster 2 showed genes which were not induced by DAHB but were induced when OXPHOS was blocked by oligomycin, and these included transcription factors of the ISR, *Atf4* and *Ddit3* (CHOP). Cluster 3 included genes induced by DAHB but were also enhanced when cells were treated with oligomycin. Cluster 4 included genes expressed by activated T cells and downregulated by DAHB and DAHB+O, including *Pcna* and *Mcm2*, genes associated with DNA replication. Finally, Cluster 5 was composed of genes expressed in activated T cells that showed modest change in response to DAHB and were decreased by oligomycin, and these included the lineage-specific genes *Cd3d* and *Cd3e*.

### Induction of the ISR partially modulates effector gene expression after OXPHOS inhibition

Given the reduction of terminal effector molecules with oligomycin, we next asked whether this was a general effect of OXPHOS inhibition. Using additional electron transport chain (ETC) inhibitors, we targeted OXPHOS and measured cytokine production (Figures 3A–3C). Consistent with the RNA-seq data, protein levels of Perforin and IFN-γ induced by DAHB were significantly reduced by complex I (piericidin), complex III (antimycin A), or complex V (oligomycin) inhibitors. However, piericidin and antimycin had little effect on the TNF-α produced by activated T cells, while oligomycin led to an increase. The effects on cell viability with any of the ETC inhibitors were mild (Figure S8A). Stress responses to ETC inhibition are dependent on cell type and metabolic state and therefore we tested the three distinct ETC inhibitors for their ability to induce the ISR^41^. DAHB significantly reduced the hallmark ISR protein, ATF4. In contrast, addition of ETC inhibitors suppressed the DAHB-mediated reduction in ATF4 expression (Figure 3D). Oligomycin, a direct ATP-synthase inhibitor, was most potent in reversing DAHB-mediated ATF4 suppression. To explore the function of the ISR in T cell effector protein induction, we added the inhibitor ISRIB, which restores global translation by relieving ISR-associated inhibition of eIF2B^42^. In cells treated with DAHB and oligomycin, ISRIB suppressed induction of the ISR effectors ATF4 and CHOP while restoring global translation as measured by puromycin incorporation (Figures 3E and S8B). Inhibiting the oligomycin-mediated ISR with ISRIB did not significantly rescue loss of Perforin or IFN-γ, and the oligomycin-enhancement of TNF-α was significantly reduced (Figure 3F).

**Figure 3.**
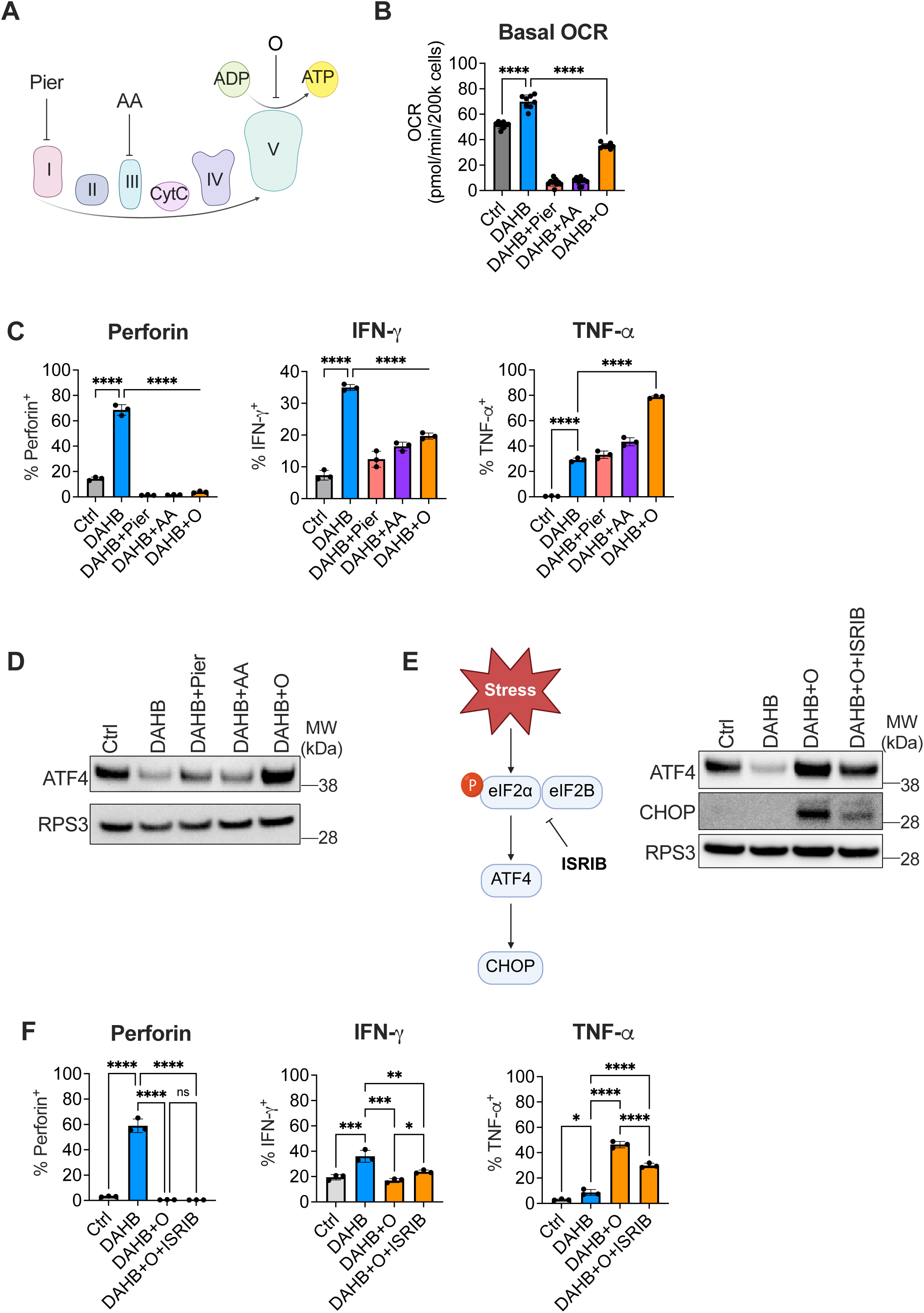
**OXPHOS inhibition modulates the effector program partly via the integrated stress response.** (A) Schematic of the electron transport chain and sites of action for inhibitors. Pier, piericidin; AA, antimycin A; O, oligomycin. (B) Basal oxygen consumption rate (OCR) at the first timepoint in activated CD8^+^ T cells treated with Ctrl, DAHB, DAHB + pier (100 nM), DAHB + AA (10 nM), or DAHB + O (10 nM). Ctrl, n = 8; DAHB, n = 8; DAHB + Pier, n = 10; DAHB + AA, n = 10; DAHB + O, n = 8. (C) Perforin, IFN-γ, and TNF-α production measured by flow cytometry in activated CD8^+^ T cells treated with Ctrl, DAHB, DAHB + Pier, DAHB + AA, or DAHB + O at concentrations as in (B); n = 3. (D) Immunoblot of ATF4 in cells treated as in (C). RPS3 serves as a loading control. Representative of two independent experiments. (E) Schematic of ISRIB mechanism of action (left) and immunoblot of ATF4 and CHOP after Ctrl, DAHB, DAHB + O, or DAHB + O + ISRIB treatments (right). RPS3 serves as a loading control. Representative of two independent experiments. (F) Perforin, IFN-γ, and TNF-α production in activated CD8^+^ T cells treated with Ctrl, DAHB, DAHB + O, or DAHB + O + ISRIB (n = 3). Data are mean ± SD. Statistical analysis in (B, C, F) was performed by one-way ANOVA with Tukey’s multiple comparisons test. ns, not significant; *p < 0.05; **p < 0.01; ***p < 0.001; ****p < 0.0001. See also Figures S6–S8.

### Mitochondrial OXPHOS enhances phosphocreatine levels

The above results suggested that mitochondrial ATP production could be contributing to the effects of DAHB on T cell gene expression. To independently assess the effects of DAHB ± oligomycin treatments on activated T cell metabolism, we next undertook unbiased metabolomics analysis of the treated cells. In activated T cells treated with DAHB, phosphocreatine (PCr) was the most significantly induced metabolite in comparison to control cells (Figure 4A). When DAHB + oligomycin-treated cells (DAHB+O) were compared to DAHB-only treatment, PCr was significantly reduced (Figure 4B). In contrast, creatine levels varied little in either situation.

**Figure 4.**
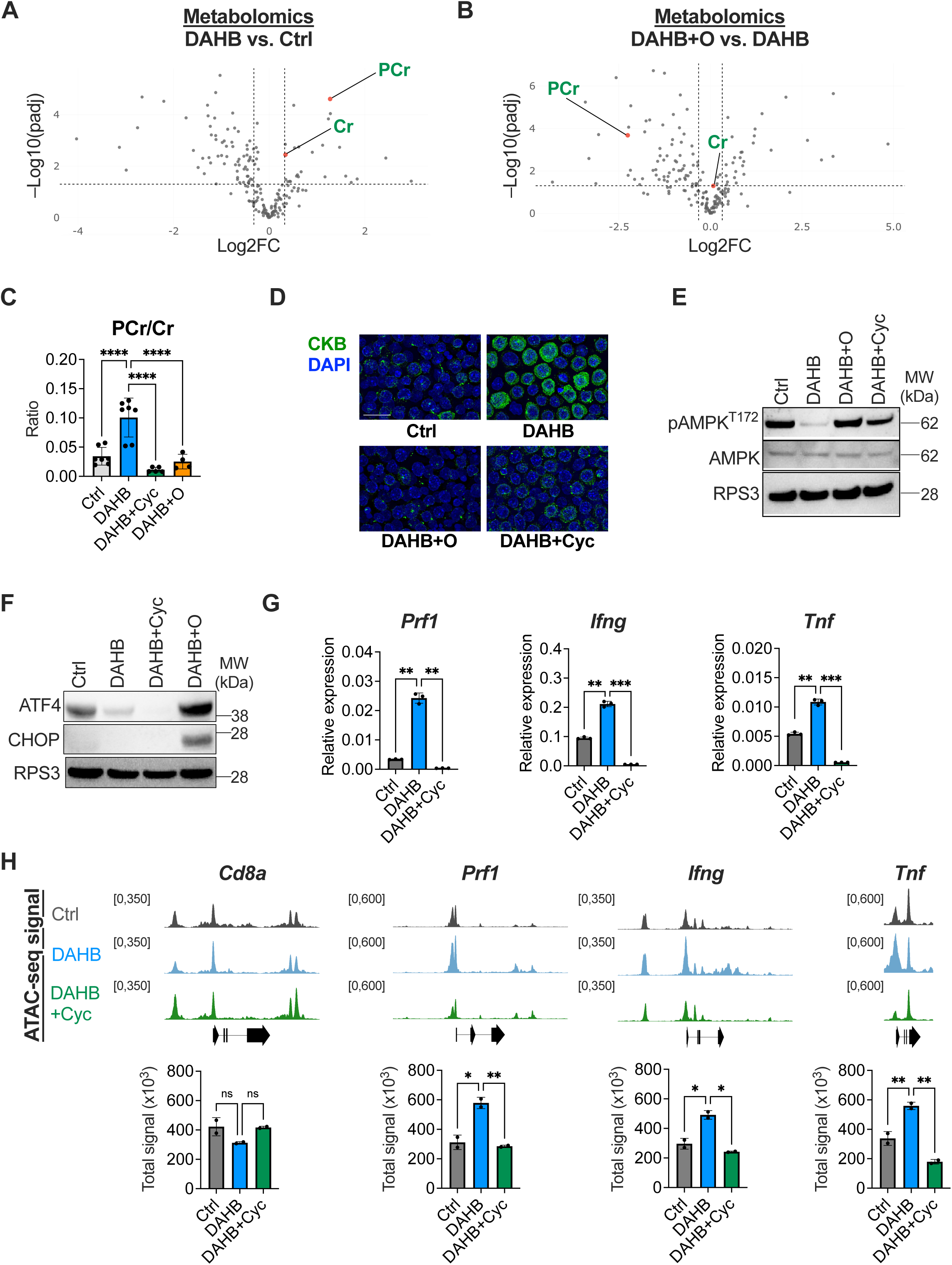
**OXPHOS-dependent phosphocreatine supports chromatin accessibility.** (A and B) Volcano plots of polar metabolomics in DAHB vs. Ctrl and DAHB + oligomycin (DAHB + O) vs. DAHB; n = 3. Highlighted in green are creatine (Cr) and phosphocreatine (PCr). (C) PCr/Cr ratio measured by LC-MS after treatment with Ctrl, DAHB, DAHB + cyclocreatine (Cyc, 10 mM), or DAHB + O in activated CD8^+^ T cells (data pooled from three experiments; n = 4–7 replicates per condition). (D) Representative immunofluorescence of CKB in activated CD8^+^ T cells treated with Ctrl, DAHB, DAHB + O, or DAHB + Cyc. CKB is pseudocolored in green and nuclei are stained with DAPI (blue). Scale bar, 20 μm. (E) Immunoblot of pAMPK^T172^ and AMPK in activated CD8^+^ T cells treated with Ctrl, DAHB, DAHB + O, or DAHB + Cyc. RPS3 serves as a loading control. Data are representative of two independent experiments. (F) Immunoblot of ATF4 and CHOP in activated CD8^+^ T cells treated with Ctrl, DAHB, DAHB + Cyc, or DAHB + O. RPS3 serves as a loading control. Data are representative of two independent experiments. (G) *Prf1*, *Ifng*, and *Tnf* expression measured by RT-qPCR after treatment with Ctrl, DAHB, or DAHB + Cyc in CD8^+^ T cells; n = 3. (H) ATAC-seq tracks at *Cd8a*, *Prf1*, *Ifng*, and *Tnf* loci in Ctrl, DAHB or DAHB + Cyc treated CD8^+^ T cells. Quantification of total signal per replicate in the region shown is plotted below. Data are representative of two independent experiments. Data are mean ± SD. Statistical analysis in (A, B) was determined by two-sided Welch’s t test with Benjamini-Hochberg correction (adjusted p < 0.05 and fold change > 1.25). Statistical analysis in (C, G, H) was performed by one-way ANOVA with Tukey’s multiple comparisons test. ns, not significant; *p < 0.05; **p < 0.01; ***p < 0.001; ****p < 0.0001. See also Figure S9.

Creatine metabolism plays a well-established role in energy storage in terminally differentiated muscle cells, neurons and in converting ETC activity into thermogenesis in brown adipocytes^43,44^. Recent studies also reported increased creatine uptake in activated T cells but how increased creatine contributes to T cell activation is unclear^45,46^. Furthermore, the canonical mechanism described in skeletal muscle demonstrated that it is the ratio of PCr relative to creatine (PCr/Cr) that provides a bioenergetic reserve that muscle-type creatine kinase (CKM) utilizes to convert ADP to ATP at the sarcolemma^47,48^. Although CKM is not expressed in T cells, T cells express the homologous enzyme brain-type CK (CKB).

The DAHB-induced increase in PCr was effectively blocked by either the OXPHOS inhibitor oligomycin or cyclocreatine (Cyc), a creatine analog that competitively inhibits PCr production^49^ (Figure 4C). Interestingly, the CKB enzyme that converts mitochondrial-derived phosphocreatine into ATP by phosphate transfer to ADP was the most potently induced metabolic enzyme when activated T cells were treated with DAHB (Figures 1C and 1D). The level of CKB induction as assayed by RNA-seq and proteomics was comparable to that found for T cell effector genes. Cytologic evaluation of DAHB treatment confirmed CKB was upregulated (Figure 4D). This suggested that mitochondrial OXPHOS might contribute to gene expression changes in activated T cells through an energy-shuttling mechanism. Consistent with an elevated bioenergetic status, DAHB treatment reduced the AMPK phosphorylation observed in activated T cells which was eliminated by treatment with Cyc or oligomycin (Figure 4E). Despite an increased ratio of PCr/Cr, the ratio of ATP/ADP remained constant, suggesting that creatine metabolism may promote distribution of phosphate equivalents throughout the cell to support location-specific ATP buffering rather than affecting the overall abundance of ATP to ADP (Figure S9A). Consistent with increased OXPHOS, DAHB-treated T cells also had reduced NADH/NAD^+^ and lactate/pyruvate ratios. This occurred without significantly affecting glutathione reserves. Interestingly, the increase in PCr/Cr was limited to the addition of the D-isomer of AHB, as LAHB-treated cells had no change in PCr/Cr (Figure S9B). To further confirm the role of creatine metabolism, we used a recently developed covalent creatine kinase inhibitor (CKi)^50^. CKi treatment collapsed the PCr/Cr ratio, activated AMPK and diminished cytokine production (Figures S9C–S9E).

Unlike oligomycin treatment, Cyc did not activate the ISR (Figure 4F). However, Cyc significantly reduced Perforin and IFN-γ at both the protein and transcript levels (Figures 4G and S9E). While TNF-α transcripts were diminished by treatment with Cyc, neither Cyc nor CKi decreased its protein levels, consistent with post-transcriptional regulation^51,52^. Either Cyc or oligomycin treatment also significantly reduced DAHB-enhanced cytotoxicity in tumor co-culture assays (Figure S9F).

Given the PCr-dependence on effector gene expression and cytotoxic function, we examined whether chromatin remodeling was also affected in an ATP-dependent fashion. We performed ATAC-seq in activated CD8^+^ T cells treated with Ctrl, DAHB or DAHB + Cyc. DAHB enhanced accessibility at the effector genes *Prf1* (Perforin), *Ifng* (IFN-γ) and *Tnf* (TNF-α), which was significantly reduced by the addition of Cyc (Figure 4H). In contrast, the control T cell lineage gene, *Cd8a*, showed no significant changes. As creatine kinase is required for phosphocreatine to generate ATP, we asked whether any of the CKB induced by DAHB treatment was observed in the nucleus. Imaging by STORM revealed that a portion of the CKB protein induced in response to DAHB localized to the nuclear periphery. Using antibodies to proteins known to localize near the nuclear envelope, we found that the fluorescence signals from Lamin B1 and CKB closely approximated each other (Figure S9G). To confirm CKB was spatially adjacent to Lamin B1, we performed proximity ligation assays. DAHB induced over six-fold enrichment in nuclear puncta between CKB and Lamin B1, confirming that part of the induced CKB is in close spatial proximity to the Lamin B1 network that lines the inner nuclear membrane^53^ (Figure S9H).

### DAHB-mediated bioenergetics supports ATP-dependent chromatin remodeling

The bioenergetic hurdle of changing chromatin accessibility is mediated by ATP-dependent nucleosome remodeling complexes. To directly assess chromatin remodeling, we asked whether inhibiting the BAF chromatin remodeling complex, known to play a role in CD8^+^ T cells, was sufficient to phenocopy the effects of PCr inhibition^54,55^ (Figure 5A). First, we targeted the BAF complex with the proteolysis-targeting chimera (PROTAC) AU-24118^56^ (AU). Degradation of the BAF complex led to reductions in DAHB-mediated IFN-γ and Perforin expression (Figures 5B and 5C). Next, we used a specific Brg1/Brm inhibitor, BRM014^57^, to inhibit the ATPase activity of the BAF complex and obtained similar results to degradation with the PROTAC (Figure 5D). Similarly to Cyc treatment, TNF-α expression was maintained at the protein level while transcript levels were repressed to that of Ctrl (Figure 5E). At the level of chromatin, BAF inhibition led to loss of accessibility at effector loci, phenocopying the effects we observed with Cyc (Figure 5F).

**Figure 5.**
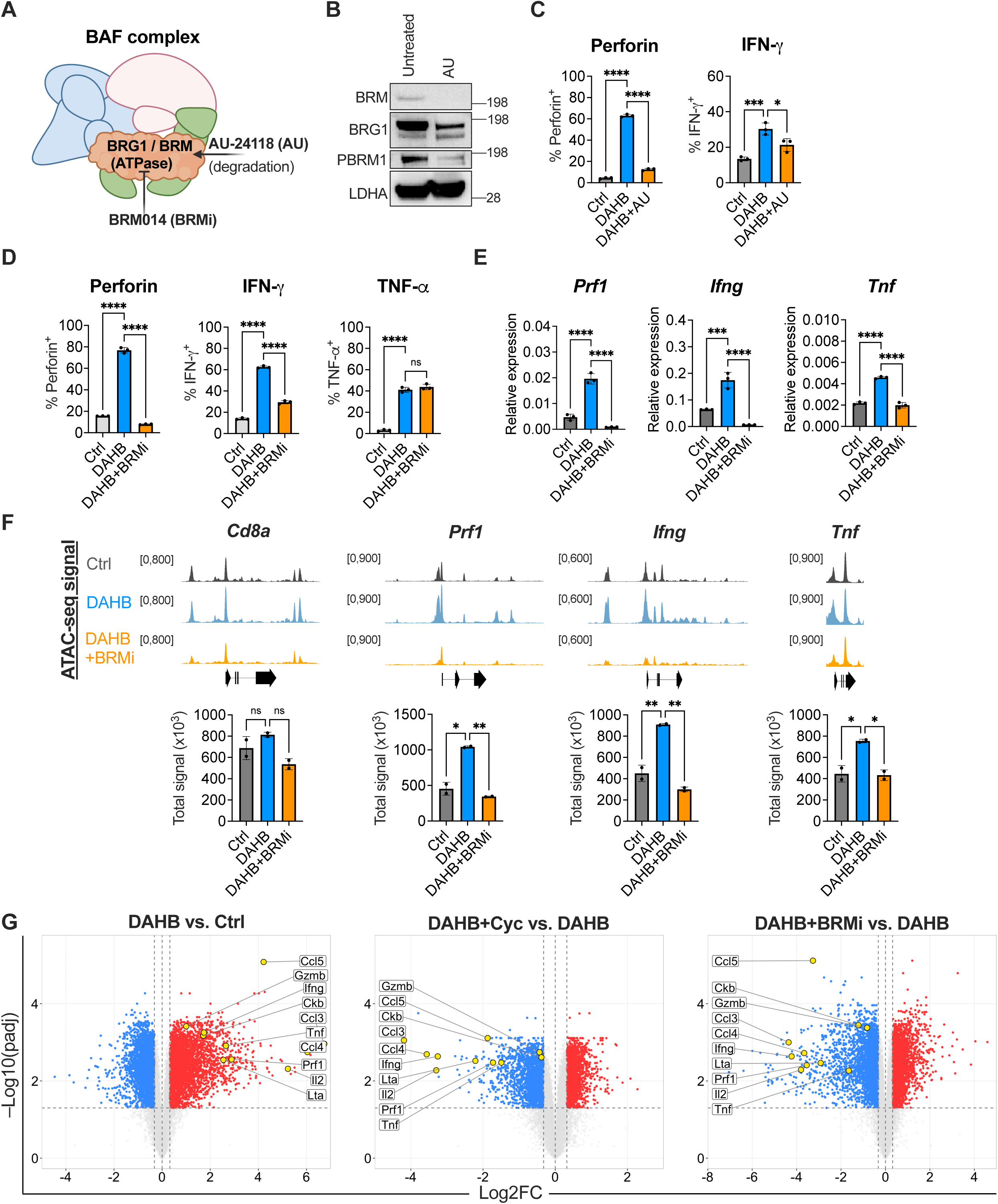
**The ATP-dependent chromatin remodeling complex BAF is required for DAHB-mediated effector gene expression.** (A) Diagram of BAF complex with the ATPase inhibitor BRM014 (BRMi, 1 μM) and the PROTAC AU-24118 (AU, 1 μM). (B) Immunoblot of BAF complex subunits BRM, BRG1 and PBRM1 after overnight treatment with AU in activated CD8^+^ T cells. LDHA serves as a loading control. Representative of two independent experiments. (C) Perforin and IFN-γ production after treatment with Ctrl, DAHB, or DAHB + AU; n = 3. (D) Perforin, IFN-γ and TNF-α protein levels measured by flow cytometry after treatment with Ctrl, DAHB, or DAHB + BRMi; n = 3. (E) *Prf1*, *Ifng*, and *Tnf* expression measured by RT-qPCR after DAHB ± BRMi treatment in activated CD8^+^ T cells as in (D); n = 3. (F) ATAC-seq tracks at *Cd8a*, *Prf1*, *Ifng* and *Tnf* loci after BRMi treatment. Quantification of total signal in the region shown per replicate is plotted below. Data are representative of two independent experiments. (G) RNA-seq volcano plots of CD8^+^ T cells comparing DAHB vs. Ctrl, DAHB + cyclocreatine (Cyc) vs. DAHB, and DAHB + BRMi vs. DAHB. Representative effector genes are highlighted in yellow; n = 3. Data are mean ± SD. Statistical analysis in (C, D, E, F) was performed by one-way ANOVA with Tukey’s multiple comparisons test. Differential expression in (G) was determined by DESeq2 with Benjamini-Hochberg correction (adjusted p < 0.05 and fold change > 1.25). ns, not significant; *p < 0.05; **p < 0.01; ***p < 0.001; ****p < 0.0001.

Finally, given the similar loss of accessibility by both BAF inhibition and Cyc, we measured global gene expression changes by RNA-seq using the MSKCC Integrated Genomics Core (Figure 5G). Consistent with our initial observations (Figure 1C), DAHB broadly upregulated the CD8^+^ T cell effector program including cytokines, cytolytic molecules, and CKB, which were all reversed by treatment with either Cyc or BRMi.

### DAHB treatment *in vivo* or *ex vivo* enhances antitumor immunity

Metabolic supplementation of adoptive T cell therapies including CAR (chimeric antigen receptor) T cells is an emerging strategy for cancer immunotherapy^58–60^. Thus, we asked whether DAHB-treated T cells would enhance effector function *in vivo*. OVA-specific OT-I CD8^+^ T cells were activated *ex vivo* with anti-CD3/CD28 and cultured overnight in medium with or without DAHB. The cells were then washed in PBS and adoptively transferred into RAG2 KO mice bearing EL4-OVA tumors (Figure 6A). *Ex vivo* treatment with DAHB prior to transfer improved tumor control, demonstrating durable enhancement of T cell effector function (Figures 6B–6D). To assess how culturing OT-I T cells in DAHB prior to transfer altered their effector phenotype *in vivo*, washed DAHB- and Ctrl-treated cells that were differentially labeled with fluorescent dyes and mixed in equal proportions, were injected adjacent to EL4-OVA tumors. After 24 hours, the tumors were harvested and the TILs isolated. The relative frequency of the DAHB- and Ctrl-treated cells was similar. However, DAHB-treated cells retained significantly greater expression of IFN-γ, Perforin, TNF-α, and CKB (Figures S10A–S10C).

**Figure 6.**
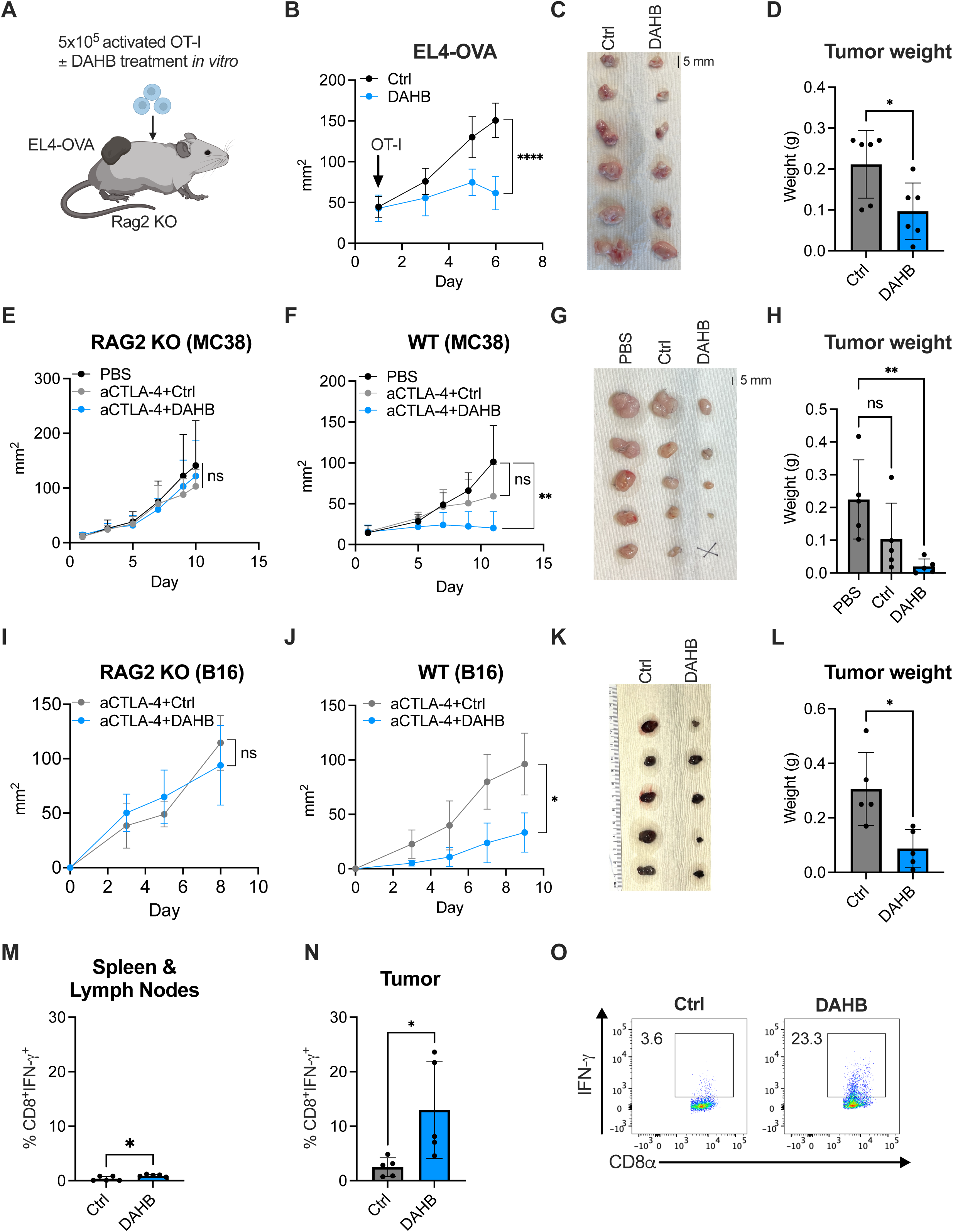
**DAHB enhances antitumor immunity *in vivo*.** (A) Schematic of adoptive transfer of activated OT-I CD8^+^ T cells treated with Ctrl or DAHB overnight prior to transfer into RAG2 KO mice bearing EL4-OVA tumors. (B) Tumor growth curves in mice from (A); n = 6 mice per group. (C and D) Photograph and weights of excised tumors from (B). (E) Tumor growth curves in RAG2 KO mice bearing MC38 tumors treated with PBS only, anti-CTLA-4 + Ctrl, or anti-CTLA-4 + DAHB; n = 5 mice per group. (F) Tumor growth curves of WT mice treated as in (E); n = 5 mice per group. (G and H) Photograph and weights of excised tumors from WT mice in (F). (I) Tumor growth curves in RAG2 KO mice bearing B16 tumors treated with anti-CTLA-4 + Ctrl or anti-CTLA-4 + DAHB; n = 5 mice per group. (J) Tumor growth curves in WT mice bearing B16 tumors treated as in (I); n = 5 mice per group. (K and L) Photograph and weights of excised tumors from WT mice in (J). (M) IFN-γ production after PMA/ionomycin restimulation in CD8^+^ T cells isolated from spleen/lymph nodes from WT mice in (J), treated with Ctrl + anti-CTLA-4 or DAHB + anti-CTLA-4; n = 5 spleen and lymph node collections. (N) IFN-γ production from tumor infiltrating lymphocytes (TILs) of WT mice in (J), treated with Ctrl + anti-CTLA-4 or DAHB + anti-CTLA-4; n = 5 tumors. (O) Representative flow cytometry of IFN-γ production in TILs from (N); n = 5 tumors. Data are mean ± SD. In each tumor model, data presented are from a representative experiment of two or more biologic replicates where n is the number of animals, tumors or tissue samples per replicate. Statistical analysis was performed by unpaired two-tailed Student’s t test (B, D, I, J, L, M, N) or one-way ANOVA (E, F, H) with Dunnett’s multiple comparisons test. For tumor growth curves (B, E, F, I, J), significance is shown for the endpoint tumor areas. ns, not significant; *p < 0.05; **p < 0.01; ***p < 0.001; ****p < 0.0001. See also Figure S10.

We next tested combining DAHB with immune checkpoint blockade using anti-CTLA-4. To determine if effective doses could be achieved *in vivo*, we measured the half-life of DAHB in mouse plasma (Figure S10D). Using a dose comparable to that previously used for L-lactate^61^, injection of either DAHB or LAHB led to *in vivo* blood levels maintained in the millimolar range for several hours (t_­_= 28 minutes for DAHB vs. 24 minutes for LAHB). We observed significant levels of AKB in the circulation within 15 minutes of injection of LAHB while only modest elevations of AKB were observed following DAHB injection (Figure S10E). In RAG2 KO mice lacking an adaptive immune system, established tumors of MC38 colorectal tumor cells showed no difference in tumor growth between anti-CTLA-4 + Ctrl, anti-CTLA-4 + DAHB or PBS alone (Figures 6E and S10F). When wild-type (WT) mice bearing MC38 tumors were treated with anti-CTLA-4 + Ctrl there was a modest and nonsignificant reduction in tumor burden. In contrast, animals treated with both anti-CTLA-4 + DAHB had a significant reduction in tumor burden, including one complete tumor rejection (Figures 6F–6H).

We also tested the combination of anti-CTLA-4 + DAHB in both RAG2 KO and WT animals harboring established B16 melanoma tumors (Figure S10G). We found DAHB also enhanced suppression of tumor growth in this model, and this antitumor activity depended on an intact adaptive immune system (Figures 6I–6L). Intratumoral CD8^+^ T cells from WT animals treated with DAHB + anti-CTLA-4 showed significant elevation of IFN-γ compared to animals treated with Ctrl + anti-CTLA-4 (13.0 ± 8.0% vs. 2.5 ± 1.5%) (Figures 6M–6O). In contrast, T cells isolated from secondary lymphoid organs from animals receiving DAHB + anti-CTLA-4 had a more modest increase in IFN-γ over animals treated with anti-CTLA-4 alone when restimulated *in vitro* (1.0 ± 0.2% vs. 0.4 ± 0.4%).

### DAHB enhances human T cell effector function

The above results prompted us to examine whether DAHB could have a similar effect on the *in vitro* and *in vivo* effector phenotype and function of human T cells. To initially assess the effect of DAHB in human T cells, we utilized an established model of T cell activation and subsequent exhaustion^8,62^. Human CD8^+^ T cells treated with DAHB showed markedly increased production of IFN-γ and TNF-α, and this was significantly maintained during chronic stimulation (Figures 7A–7C). The expression of inhibitory receptors PD-1 and CD39 was upregulated in chronically stimulated cells as previously reported^8^ and was increased by DAHB (Figures S11A and S11B).

**Figure 7.**
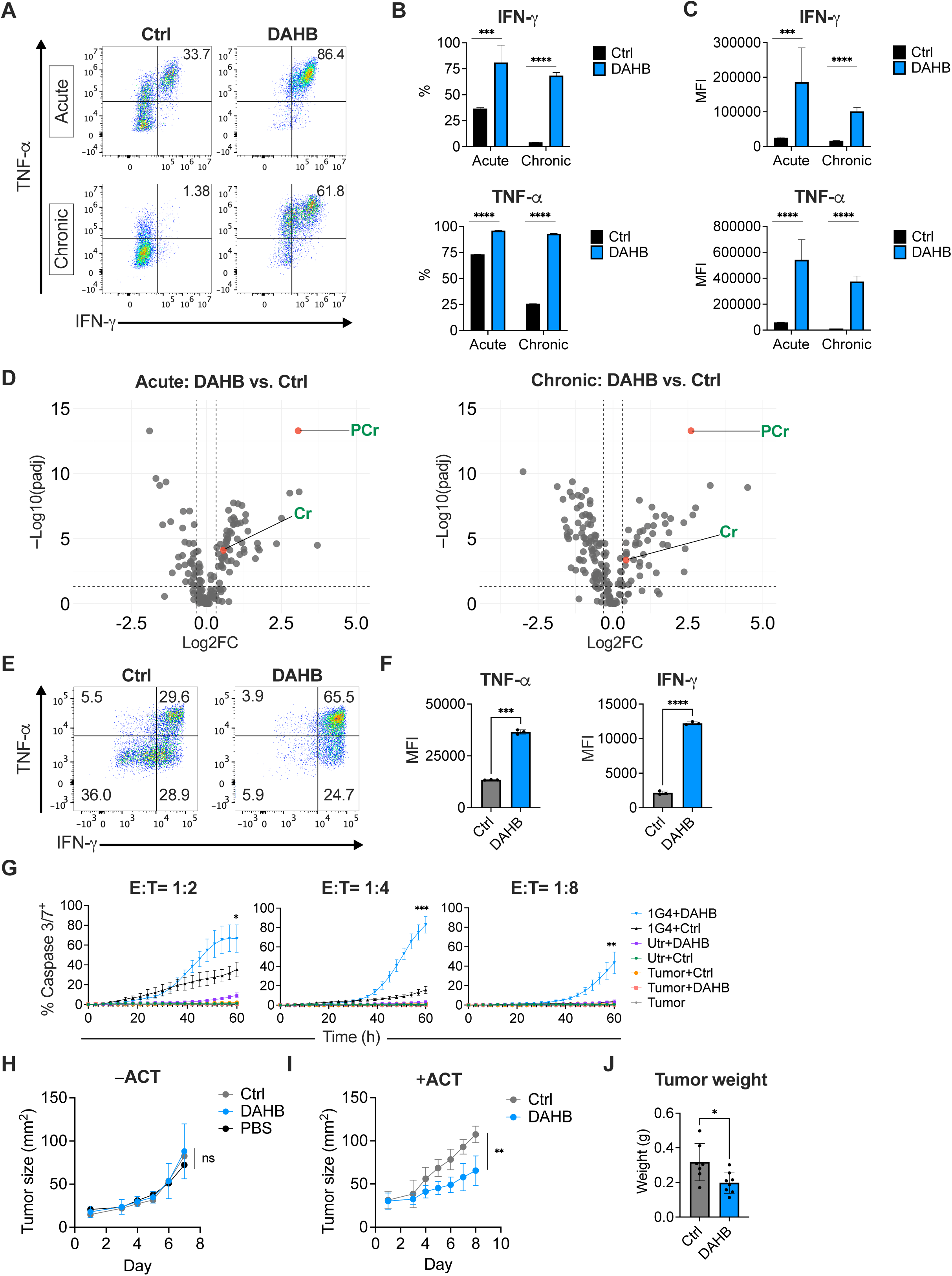
**DAHB enhances human T cell effector function.** (A) IFN-γ and TNF-α production from total T cells gated on CD8^+^, treated with NaCl (Ctrl) or DAHB from day 2 to day 8 post-activation under acute or chronic stimulation conditions as previously published^76^. Cells were analyzed on day 8. Representative plots are shown from one donor. (B and C) Quantification of cytokine production from triplicate wells from the donor shown in (A). Percent positive (B) and MFI (C) of IFN-γ (top) and TNF-α (bottom). Similar results were obtained from two additional independent donors. (D) Metabolomics volcano plot from acute (left) and chronic (right) stimulated total human T cells cultured as in (A). Phosphocreatine (PCr) and creatine (Cr) are highlighted in green. Similar results were obtained from a second independent donor. (E) Representative flow plots of IFN-γ and TNF-α production in 1G4-transduced human CD8^+^ T cells pretreated with Ctrl or DAHB for 48 h. Similar results were obtained from a second independent donor. (F) MFI of TNF-α and IFN-γ production from (E); n = 3. (G) Cytotoxicity assay of NY-ESO-1^+^mCherry^+^ A375 cells during co-culture with 1G4-transduced human CD8^+^ T cells treated as in (E) at E:T ratios of 1:2, 1:4, and 1:8 (triplicate wells). Controls include untransduced (Utr) CD8⁺ T cells ± DAHB, and A375 without T cells alone or treated with Ctrl or DAHB. Percent tumor apoptosis (% Caspase3/7^+^) was calculated as the percentage of A375 cells marked by mCherry also positive for caspase3/7 fluorescence. (H) Tumor growth curves of A375 xenografts in NSG mice without adoptive cell therapy (−ACT) treated daily by i.p. injection with NaCl (Ctrl), DAHB, or PBS only (Ctrl, n = 6 mice; DAHB, n = 5 mice; PBS, n = 3 mice). (I) Tumor growth curves of A375 xenografts in NSG mice treated by ACT of 1G4-transduced human CD8^+^ T cells preconditioned with Ctrl or DAHB (20 mM) for 48 h, plus daily i.p. injections of Ctrl or DAHB (Ctrl, n = 7 mice; DAHB, n = 8 mice). The data presented are from 1 of 2 biologic replicates, each using T cells from a different donor. (J) Endpoint tumor weights from mice in (I). Data are mean ± SD. Statistical analysis was performed by two-way ANOVA with Holm-Sidak’s multiple comparisons test (B, C), unpaired two-tailed Student’s t-test (F, I, J), or one-way ANOVA with Tukey’s multiple comparisons test (G, H). For (D), differential metabolite abundance was assessed using a two-sided Welch’s t-test with Benjamini-Hochberg correction; significant metabolites were defined as adjusted p < 0.05 and fold change > 1.25. For (G), significance is shown at the endpoint of each E:T ratio comparing DAHB- vs. Ctrl-treated 1G4-transduced CD8^+^ T cells. For (H, I), significance is shown at the study endpoint. ns, not significant; *p < 0.05; **p < 0.01; ***p < 0.001; ****p < 0.0001. See also Figure S11.

Metabolomics revealed that DAHB-treated human T cells under both acute and chronic stimulation upregulated PCr (Figure 7D). Additional human T cell metabolic similarities to those observed in murine T cells upon DAHB treatment included increased abundance of upstream glycolytic intermediates and decreased lactate and NADH levels (Figure S11C). Metabolites in the pentose phosphate pathway (PPP) were also increased, correlating with the reported role of the PPP in promoting T cell effector function^63^. DAHB-treated human T cells maintained their ATP/ADP ratio under chronic stimulation, consistent with PCr buffering, while human T cells chronically stimulated in the absence of DAHB had a significantly reduced ATP/ADP ratio as previously demonstrated (Figure S11D)^8^. Cyclocreatine addition significantly reduced the ability of DAHB to enhance effector cytokine production in both acute and chronically stimulated cells (Figures S11E and S11F).

To determine if DAHB-treated human CD8^+^ T cells could enhance their antitumor activity, we transduced T cells with the well-characterized TCR 1G4 which recognizes NY-ESO-1_­_ presented by HLA-A*02:01^64^. Upon restimulation, transduced human CD8^+^ T cells expressing 1G4 showed enhanced cytokine production after DAHB treatment (Figures 7E and 7F). Co-culture with the HLA-A*02:01^+^, NY-ESO-1^+^ human melanoma cell line A375 as target cells revealed enhanced cytotoxicity by DAHB-treated T cells (Figure 7G). In contrast, treatment of target cells with DAHB alone or with untransduced (Utr) CD8^+^ T cells ± DAHB had minimal tumor cell killing.

Finally, we asked if DAHB treatment of human CD8^+^ T cells led to greater antitumor activity *in vivo*. Using a xenograft tumor model in immunodeficient NSG mice, we found that established A375 melanoma tumors in mice treated with equivalent doses of DAHB, NaCl (Ctrl), or PBS i.p. had comparable growth rates (Figure 7H). However, when mice bearing established A375 tumors received adoptive cell therapy (ACT) with human CD8^+^ T cells transduced with TCR 1G4, pretreatment of the infused cells with DAHB followed by i.p. DAHB administration significantly decreased tumor growth and weight (Figures 7I, 7J and S11G).

## Discussion

Metabolites have emerged as regulators of cell differentiation and function. Here we found that DAHB reprograms T cell metabolism to increase chromatin accessibility and effector gene expression. By stimulating mitochondrial oxidative phosphorylation, DAHB allows T cells to build and sustain a bioenergetic reserve that supports T cell effector function.

Like other microbial metabolites that modulate immune function, DAHB is primarily produced by bacteria as a fermentation end product^23,65^. DAHB enters activated T cells through MCT transporters (Figures S2F–S2H). However, in contrast to other monocarboxylates that enter T cells, DAHB is not a meaningful metabolic substrate. Stable isotope tracing of [U-^13^C]DAHB revealed no labeling of TCA intermediates or oxidation to AKB, and cytokine induction was observed in cells lacking the DAHB-catabolizing enzyme LDHD^33^ (Figure S3). Instead, DAHB stimulated mitochondrial ATP production while suppressing aerobic glycolysis and lactate production. As mitochondrial oxidative phosphorylation increased in response to DAHB, cells became less dependent on glycolysis for ATP production (Figures 2A–2C), and more glucose-derived carbon was diverted into other biosynthetic pathways. DAHB-treated cells increased glucose flux into UDP-GlcNAc and the amino acids serine and glycine (Figure S3H). Thus, DAHB treatment promotes anabolic metabolism by shifting glucose utilization from export as L-lactate into the production of precursors for glycosylation and nonessential amino acid synthesis.

DAHB-enhanced ETC activity is supported by a shift of T cells toward fatty acid oxidation (Figures 2D–2F). While CD8^+^ T cells transitioning out of clonal expansion in secondary lymphoid organs maintain their bioenergetic potential through aerobic glycolysis, CD8^+^ T cell effector function may benefit from enhanced mitochondrial ATP generation to preserve glucose and glycolytic intermediates in support of macromolecular synthesis. The cytoskeletal rearrangements that support T cell migration in peripheral tissues are reported to depend on OXPHOS-derived ATP^66^. Shifting ATP production toward mitochondrial OXPHOS may also be beneficial during nutrient limitation at sites of wound repair or in the tumor microenvironment. Tumor-infiltrating effector T cells have been found to take up lipids to maintain effector function but excessive uptake of oxidized lipids in the tumor microenvironment has been associated with T cell dysfunction^67–70^. By inducing mitochondrial ETC activity and the building of a bioenergetic reserve, DAHB treatment buffers the potential toxicity of increased lipid oxidation. The use of FAO to support mitochondrial ATP production allows cells to preserve glycolytic carbon for biosynthetic purposes. Importantly, promoting mitochondrial function can protect against T cell exhaustion caused by chronic antigen stimulation^6–8^. In a human T cell model of exhaustion, DAHB-treated cells maintained robust cytokine production (Figures 7A–7C). DAHB treatment also induced significant upregulation of PCr and preserved the ATP/ADP ratio during chronic antigen stimulation (Figures 7D and S11D).

DAHB treatment dampened multiple stress-sensing pathways, including AMPK phosphorylation and expression of ISR effector proteins ATF4 and CHOP. These effects were reversed by pharmacologic inhibition of OXPHOS, indicating that mitochondrial ATP production is required to restrain multiple stress-sensing pathways. Notably, the strongest inducer of the ISR in CD8^+^ T cells was oligomycin (Figure 3D). This induction of the ISR mirrors the oligomycin response observed in skeletal muscle myotubes, raising the possibility of a shared metabolic state in effector cells across lineages^41^. While DAHB decreased expression of these stress sensors which may contribute to increased effector function, such homeostatic pathways can still provide supportive signals. For example, the KO of ATF4 has been shown to compromise T cell metabolic reprogramming and AMPK can promote IFN-γ production after restimulation^71,72^. Similarly, we observed AMPK KO reduced DAHB-mediated cytokine induction compared to WT cells (Figures S5J–S5L).

With an increase in ETC activity, mitochondrial production of ATP increases but this is limited by ADP availability in the mitochondrial matrix. The conversion of mitochondrial ATP to PCr in the mitochondrial intermembrane space allows the rapid return of ADP to the mitochondrial matrix for use by the ATP synthase. In turn, the PCr formed diffuses from the intermembrane space up to 7-fold faster than ATP, allowing the cell to build a bioenergetic reserve in the cytosol^73^. This form of building a bioenergetic reserve has functional roles in differentiated tissues, primarily muscle and brain^43^. The signaling that directs an increase in CK activity has remained elusive but our data suggest DAHB is one such signaling molecule. DAHB-treated T cells accumulate PCr in an OXPHOS-dependent manner (Figures 4A–4C). Pharmacologic blockade of PCr led to AMPK activation without inducing the ISR and repressed DAHB-mediated effector gene expression (Figures 4E–4G).

While DAHB, like other butyrate analogs, can inhibit HDAC-mediated deacetylation of histones, this alone was insufficient to stimulate effector gene expression in the absence of mitochondrial ATP production. In contrast, blocking PCr synthesis reduced chromatin accessibility at effector gene loci and recapitulated key features of the loss of effector gene expression and accessibility observed through inhibition of the SWI/SNF chromatin remodeling complex, BAF (Figures 4H, 5F and 5G). BAF-mediated chromatin remodeling is a central regulator of effector gene programs and dysregulation of BAF-complex activity has been implicated in the development of T cell exhaustion^62,74,75^. Thus, while mitochondria are required to produce ATP in support of effector function, the mechanisms enabling mitochondrial bioenergetics to meet localized energy demands in the nucleus are unclear. The fact that DAHB treatment increases CKB expression and localization to the nuclear lamina provides a potential mechanism to sustain ATP in a compartment-specific manner for nuclear processes, including chromatin remodeling (Figures S9G and S9H).

Functionally, the metabolic and epigenetic reprogramming conferred by DAHB translated into enhanced antitumor activity *in vivo* by both murine and human T cells. DAHB treatment of mouse T cells improved tumor control in adoptive transfer and immune checkpoint blockade models of transplantable tumors (Figure 6). Importantly, we also found adoptive transfer of human T cells combined with DAHB treatment led to superior tumor control in a xenograft model (Figures 7I and 7J). The adoptive transfer of T cells pretreated with DAHB indicates it has both a direct and epigenetic effect on T cells that support antitumor immunity.

In summary, the D-stereoisomer of α-hydroxybutyrate, DAHB, induces mitochondrial OXPHOS leading to increased PCr buffering, enhanced chromatin remodeling, and reduced inhibitory stress signaling in activated T cells. These findings establish a mechanistic link between mitochondrial ATP production and chromatin accessibility and support the use of metabolic adjuvants for antitumor immunity. Given the stereochemical diversity found in microbial metabolomes, it is possible that non-mammalian enantiomers like DAHB constitute an underappreciated class of immunoregulatory signals.

## Limitations of the Study

While this study identifies DAHB as a potent immunometabolic regulator of cytotoxic T cell function *in vitro* and *in vivo*, the *in vivo* effects of DAHB on other cells in the tumor microenvironment including the tumor itself were not assessed. It is also unclear if the doses of DAHB used here in murine models will be tolerable in humans. There was no observable toxicity of the doses used, nevertheless clinical translation will require long-term safety studies of pharmacokinetics, biodistribution, and dosing optimization. The fact that DAHB stimulates ETC activity in T cells in the absence of its known metabolizing enzyme, mitochondrial LDHD, suggests it is the direct signaling molecule, but the molecular sensor remains unknown. DAHB stimulation of OXPHOS suppresses glycolysis and promotes the use of glycolytic carbon to support other cellular processes. Whether other lineages can utilize a shift to mitochondrial oxidative phosphorylation to support cellular bioenergetics and chromatin remodeling during differentiation remains to be investigated. While these studies identify a novel role for stereochemistry in how immune cells sense and respond to their environment, the role of stereochemistry in metabolic regulation is just beginning to emerge.

## RESOURCE AVAILABILITY

### Lead contact

Further information and requests for resources and reagents should be directed to the lead contact, Craig Thompson.

## Materials availability

Material requests can be directed to the lead contact.

## Data and code availability

- RNA-seq data have been deposited as GSE304270, GSE304271, and GSE310052.

ATAC-seq data have been deposited as GSE304268 and GSE304269. Proteomics data have been deposited via ProteomeXchange (PRIDE) as PXD066950. Additional processed data for metabolomics, RNA-seq, proteomics, histone modifications (LC-MS), and raw data for pharmacokinetics, proximity ligation images and uncropped western blots have been deposited at Mendeley (DOI: 10.17632/b2yktg2b92.1).

- This paper does not report original code.
- Any additional information required to reanalyze the data reported in this paper is available from the lead contact upon request.

## ACKNOWLEDGMENTS

This work is supported by the NCI Cancer Center Support Grant (CCSG, P30 CA008748) to MSKCC. The Thompson lab is supported by a grant from the NIH/NCI (R35 CA283988) and the Thompson Research Lab Funds. C.N. is supported by the Weill Cornell Department of Pathology through NIH 5T32CA260293-03. N.S.C. is supported by the NIH (R01CA290678 and R01AI148190). C.A.K. was supported in part by NIH R37 CA259177, R01 CA286507, R01 P50 CA217694, and P30 CA008748; W. H. Goodwin and A. Goodwin and the Commonwealth Foundation for Cancer Research; The Center for Experimental Therapeutics at Memorial Sloan Kettering Cancer Center; The Parker Institute for Cancer Immunotherapy; Cycle for Survival; and the Metropoulos Family Foundation. We thank Yevgeniy Romin and Murray Tipping for their assistance and expertise working with the STORM super-resolution imaging.

## AUTHOR CONTRIBUTIONS

Experiments were performed by C.N., T.S.F., D.L., K.K., L.F.S.P., A.M., D.N.W., O.J., J.K., H.T., Y.H., Z.X., Z.Y., G.B., Y.L., and E.M.S. Experiments were conceived by C.N., S.P.C., X.C., S.A.V., C.A.K., N.S.C., L.W., and C.B.T. Data analysis was performed by C.N., R.K., G.Z., Z.L., and M.M. The manuscript was written and edited by C.N. and C.B.T. All authors reviewed the manuscript and approved the final version of the manuscript for submission.

## Supplementary figure legends

**Figure S1. DAHB enhances durable effector function without significant toxicity, related to Figure 1**.

(A) Representative gating strategy to determine cytokine production in activated CD8^+^ T cells treated with Ctrl, D-lactate or DAHB.

(B and C) Perforin and IFN-γ production by percent positive (B) or MFI (C) of CD8^+^ T cells treated as in Figure 1B (n = 3).

(A) (D) Viability using live/dead dye after indicated treatments (20 mM for LAHB, DAHB, LBHB, DBHB and 0.5 mM for butyrate); n = 3.

(B) (E) Viability using live/dead dye during butyrate titration (0–4 mM); n = 3.

(C) (F) Perforin production by flow cytometry after DAHB titration (2–20 mM) compared with Ctrl. Data are representative of three independent experiments.

(D) (G) Activated CD8^+^ T cells were treated for 24 h with Ctrl or DAHB, washed and cultured for an additional 24 h prior to restimulation and measurement of Perforin and IFN-γ by flow cytometry (n = 3).

(E) (H) OT-I CD8^+^ T cells pretreated with Ctrl or DAHB and assessed for cytotoxicity against EL4-OVA via titration of effector-to-target (E:T) cell ratios (0:1, 1:1, 5:1, 20:1); n = 3.

Data are mean ± SD. Statistical analysis was performed by one-way ANOVA with Tukey’s multiple comparisons test (B–E) or unpaired two-tailed Student’s t test (G,H; calculated from each E:T ratio for H). ns, not significant; *p < 0.05; **p < 0.01; ***p < 0.001; ****p < 0.0001.

**Figure S2. DAHB entry is mediated by monocarboxylate transporters (MCTs) and induces prolonged effects on OXPHOS, related to Figure 2**.

(A) Oxygen consumption measured by continuous oximetry over 24 h in activated CD8^+^ T cells with treatment of Ctrl or DAHB initiated at time 0 (n = 8 wells per condition).

(B) Lipid peroxidation after 24 h treatment with Ctrl or DAHB in activated CD8^+^ T cells as measured by the FITC/PE MFI ratio of BODIPY C11 staining; n = 3. RSL3, an inducer of lipid peroxidation, was added to untreated wells prior to measurement.

(C) Seahorse plot of oxygen consumption rate (OCR) in activated CD8^+^ T cells treated with Ctrl or DAHB for 48 h and maintained in assay medium with Ctrl or DAHB during the Seahorse assay; n = 6 wells per condition.

(D and E) Basal and maximal OCR quantification from (C). Basal OCR represents the mean of the three pre-injection measurements, and maximal OCR represents the mean of the three post-FCCP measurements; n = 18 measurements (6 wells per measurement at three timepoints).

(F) Intracellular acidification measured as the pHrodo Red AM MFI ratio after immediate treatment with the indicated metabolites, normalized to NaCl (Ctrl); NH_­_Cl is included as an alkalinizing control; n = 3.

(G) pHrodo Red AM MFI ratio from (F) after 30 min incubation with the indicated treatments normalized to Ctrl; n = 3.

(H) pHrodo Red AM MFI ratio normalized to Ctrl after pretreatment with DMSO (Veh) or syrosingopine (Syro, 1 μM), followed by acute administration of the indicated metabolites; n = 3.

Data are mean ± SD. Statistical analysis was performed by unpaired two-tailed Student’s t test (A, D, E) or one-way ANOVA with Tukey’s multiple comparisons test (B, F, G, H). ns, not significant; *p < 0.05; **p < 0.01; ***p < 0.001; ****p < 0.0001.

**Figure S3. No detectable metabolism of DAHB by LDHD despite increased mitochondrial oxidative phosphorylation, related to Figure 2**.

(A) [U-^13^C]DAHB (5 mM) and [U-^13^C]butyrate (0.5 mM) tracing for 5 h in CD8^+^ T cells showing isotopologue abundances in arbitrary units for α-hydroxybutyrate (AHB), α-ketobutyrate (AKB), citrate, α-ketoglutarate (AKG) and succinate. Data are representative of two independent experiments.

(B) Schematic of Cre recombinase-mediated targeting of *Ldhd* in *Ldhd-*flox mice.

(C) Genomic DNA (gDNA) PCR in CD8^+^ T cells from WT or KO (CD4Cre;*Ldhd*^F/F^) using primers shown in panel (B).

(D) RT-PCR from WT and KO CD8^+^ T cells with *Ldhd* primers spanning indicated exons or *Ifng* primers as a positive control.

(E) CD8^+^ T cells isolated from control (*Ldhd*^F/+^) or KO mice were restimulated and Perforin and IFN-γ production was measured by flow cytometry (n = 3).

(F) Seahorse Mito Stress Test with indicated treatments and genotypes showing basal OCR; n = 8 wells per condition.

(G) Schematic of [U-^13^C]glucose incorporation into uridine diphosphate N-acetylglucosamine (UDP-GlcNAc), serine and glycine. HBP, hexosamine biosynthetic pathway; SSP, serine synthesis pathway.

(H) [U-^13^C]Glucose fractional enrichment in UDP-GlcNAc, serine and glycine after 30 min labeling in CD8^+^ T cells treated with Ctrl or DAHB; n = 3.

Data are mean ± SD. Statistical analysis was performed by one-way ANOVA with Tukey’s multiple comparisons test (E, F) or unpaired two-tailed Student’s t test (H). ns, not significant; *p < 0.05; **p < 0.01; ***p < 0.001; ****p < 0.0001.

**Figure S4. Oxidation of multiple classes of fatty acids is increased by DAHB treatment, related to Figure 2**.

(A) CoA species profiling showing Log2(DAHB/Ctrl) ratios in activated CD8^+^ T cells; n = 3.

(B) Measurement of oxidation of isotope tracers using uniformly ^13^C-labeled ([U-^13^C]) acetate, butyrate, octanoate, nonanoate, palmitate, and oleate after 5 h labeling in CD8^+^ T cells treated with Ctrl or DAHB. Percentage labeling of individual citrate isotopologues, α-ketoglutarate (AKG) m+2, succinate m+2 and malate m+2 is shown; n = 3. SCFA, short-chain fatty acid; MCFA, medium-chain fatty acid; LCFA, long-chain fatty acid.

Data are mean ± SD. Statistical analysis was performed by one-sample t test with hypothetical value 0 (A), or multiple unpaired two-tailed Student’s t test with Holm-Sidak’s multiple comparisons test (citrate), or unpaired two-tailed Student’s t test (AKG, succinate, malate) (B). ns, not significant; *p < 0.05; **p < 0.01; ***p < 0.001; ****p < 0.0001.

**Figure S5. DAHB-mediated FAO is required to enhance cytokine production, related to Figure 2**.

(A and B) Immunoblot validation of CRISPR-mediated targeting of *Cpt1a* and *Hadhb* in activated CD8^+^ T cells electroporated with targeting or control sgRNA (sgOlfr2). β-actin serves as a loading control. Representative of two independent experiments.

(C) Frequency of activated CD8^+^ T cells producing IFN-γ (left) or Perforin (right) measured by intracellular staining in Ctrl- or DAHB-treated cells expressing sgOlfr2, sgCpt1a, or sgHadhb; n = 3.

(D) MFI of IFN-γ and Perforin by intracellular staining in cells from (C); n = 3.

(E) Chemical structures of L-palmitoylcarnitine and teglicar (Teg).

(F) [U-^13^C]Palmitate tracing for 5 h showing citrate m+2 after treatment with Ctrl, DAHB, or DAHB + Teg (20 μM); n = 3.

(G) Immunoblot of phosphorylated AMPK (pAMPK^T172^) and total AMPK protein in activated CD8^+^ T cells with the indicated treatments. RPS3 serves as a loading control. Representative of two independent experiments.

(H) Basal OCR from Seahorse assay in activated CD8^+^ T cells with Ctrl, DAHB, DAHB + Teg, or DAHB + Teg + butyrate (0.5 mM); n = 10 wells per condition.

(I) IFN-γ and Perforin production by flow cytometry (percent positive) in cells treated as in (H); n = 3.

(J) Immunoblot validation of AMPK targeting (sgPrkaa1) versus control (sgOlfr2). β-actin serves as a loading control. Representative of two independent experiments.

(K) IFN-γ production (percent positive) by flow cytometry in cells from (J); n = 3.

(L) Perforin production (percent positive) by flow cytometry in cells from (J); n = 3.

(M) Cell number and IFN-γ and Perforin production in activated CD8^+^ T cells grown in low-glucose media (1.3 mM) with concomitant treatment of Ctrl or DAHB for 48 h; n = 3.

Data are mean ± SD. Statistical analysis was performed by one-way ANOVA with Tukey’s multiple comparisons test (C, D, F, H, I, K, L) or unpaired two-tailed Student’s t test (M). ns, not significant; *p < 0.05; **p < 0.01; ***p < 0.001; ****p < 0.0001.

**Figure S6. DAHB promotes histone acetylation, inhibits HDACs, and alters histone methylation, related to Figure 3**.

(A) Total class I/II HDAC activity (RLU; relative luminescence units) measured in activated CD8^+^ T cells treated for 1 h with NaCl (Ctrl; 20 mM), LAHB (20 mM), DAHB (20 mM), LBHB (20 mM), DBHB (20 mM), or butyrate (0.5 mM). Data are representative of two independent experiments.

(B) Cell-free recombinant HDAC1 (n = 2) and HDAC2 (n = 3) activity in the presence of Ctrl (10 mM), DAHB (10 mM), or trichostatin A (TSA; 2 μM); AU, arbitrary units.

(C) Histone acetylation profiling by LC-MS shown as Z-score heatmap comparing Ctrl and DAHB; n = 3.

(D) Histone methylation profiling by LC-MS as in (C); n = 3.

(E) Immunoblot of H3K9ac and H3K27ac in activated CD8^+^ T cells treated with Ctrl, DAHB, or DAHB + oligomycin (O). Total H3 serves as a loading control. Representative of two independent experiments.

(F) H3K9ac ChIP-qPCR as a percentage of input at an intergenic region on chromosome 4 (negative control) and the promoters of *Prf1*, *Ifng* and *Ckb* in CD8^+^ T cells treated with Ctrl, DAHB, or DAHB + O; n = 3.

Data are mean ± SD. Statistical analysis was performed by one-way ANOVA with Tukey’s multiple comparisons test (A, F). ns, not significant; *p < 0.05; **p < 0.01; ***p < 0.001; ****p < 0.0001.

**Figure S7. DAHB mediates oxidative phosphorylation-dependent gene expression changes, related to Figure 3**.

(A) Principal component analysis (PCA) of RNA-seq from activated CD8^+^ T cells treated with Ctrl, DAHB, or DAHB + oligomycin (DAHB + O).

(B) K-means clustering (k = 5) of RNA-seq data from (A).

(C) Representative expression levels (counts ×10^3^) of indicated genes from each cluster; n = 3.

Data are mean ± SD. Statistical analysis was performed by one-way ANOVA with Tukey’s multiple comparisons test (C). ns, not significant; *p < 0.05; **p < 0.01; ***p < 0.001; ****p < 0.0001.

**Figure S8. ETC inhibitors show minimal effects on cell viability and ISRIB rescues translation during oligomycin treatment, related to Figure 3**.

(A) Flow cytometry quantification of viability (DAPI^−^) in activated CD8^+^ T cells treated with Ctrl, DAHB, DAHB + piericidin (Pier), DAHB + antimycin A (AA), or DAHB + oligomycin (O) as in Figure 3B; n = 3.

(B) Immunoblot of puromycin incorporation in activated CD8^+^ T cells treated with cycloheximide (CHX), Ctrl, DAHB, DAHB + O, or DAHB + O + ISRIB. The protein synthesis inhibitor CHX was added to an untreated well before puromycin addition as a negative control for translation. RPS3 serves as a loading control. Representative of two independent experiments.

Data are mean ± SD. Statistical analysis was performed by one-way ANOVA with Dunnett’s multiple comparisons test (A). ns, not significant; *p < 0.05; **p < 0.01; ***p < 0.001; ****p < 0.0001.

**Figure S9. Inhibiting creatine metabolism reduces DAHB-mediated effector function, related to Figure 4**.

(A) Summary of log2 fold change (log2FC) metabolite ratios in activated CD8^+^ T cells treated with DAHB vs. Ctrl averaged across independent experiments (n = 7). ATP/ADP, adenosine triphosphate/adenosine diphosphate; PCr/Cr, phosphocreatine/creatine; NADH/NAD^+^, reduced/oxidized nicotinamide adenine dinucleotide; GSH/GSSG, reduced/oxidized glutathione; PPP, pentose phosphate pathway; 6PG/G6P, 6-phosphogluconate/glucose-6-phosphate.

(B) Ratio of PCr to Cr measured by LC-MS in activated CD8^+^ T cells treated with Ctrl, LAHB, or DAHB; n = 3.

(C) PCr/Cr ratio measured by LC-MS in activated CD8^+^ T cells treated with Ctrl, DAHB, or DAHB + creatine kinase inhibitor (CKi); n = 3.

(D) Immunoblot of pAMPK^T172^ and total AMPK in activated CD8^+^ T cells treated with Ctrl, DAHB, DAHB + CKi, DAHB + cyclocreatine (Cyc), or DAHB + oligomycin (O). RPS3 serves as a loading control. Representative of two independent experiments.

(E) Perforin, IFN-γ, and TNF-α production measured by flow cytometry in activated CD8^+^ T cells treated as in (D); n = 3.

(F) Co-culture cytotoxicity assay with EL4-OVA tumor cells and OT-I CD8^+^ T cells pretreated with Ctrl, DAHB, DAHB + Cyc, or DAHB + O at a 10:1 effector-to-target (E:T) ratio; n = 3.

(G) STORM imaging of activated CD8^+^ T cells treated with Ctrl or DAHB. CKB is shown in green, Lamin B1 is shown in red, and merged images are shown in yellow; scale bar, 2 μm. Representative of two independent experiments.

(H) Proximity ligation assay for CKB and Lamin B1 interaction. Data represent the number of puncta (pink) per nucleus across 12 randomly chosen fields (∼25 cells/field) acquired at 100× magnification; scale bar, 10 μm. Representative of three independent experiments.

Data are mean ± SD. Statistical analysis was performed by one-sample t test with a hypothetical mean of 0 (A), one-way ANOVA with Tukey’s multiple comparisons test (B, C, E, F) or unpaired two-tailed Student’s t test (H). ns, not significant; *p < 0.05; **p < 0.01; ***p < 0.001; ****p < 0.0001.

**Figure S10. DAHB promotes effector function *in vivo* and can reach therapeutic levels, related to Figure 6**.

(A) OT-I CD8^+^ T cells treated with Ctrl or DAHB were mixed after labeling with CFSE (FITC) or CellTrace Far Red (APC) (left) and tumor-infiltrating lymphocyte (TIL) frequency measured after tumor harvest (right); n = 5 tumors.

(B) IFN-γ, Perforin, and TNF-α production in restimulated OT-I CD8^+^ T cells from (A); n = 5 tumors.

(C) MFI of creatine kinase B (CKB) expression in cells from (A); n = 5 tumors.

(D) Pharmacokinetics of DAHB and LAHB following i.p. injection, plotted as plasma concentration vs. time. Data are representative of two independent experiments.

(E) Plasma α-ketobutyrate (AKB) levels over time following administration of DAHB or LAHB as in (D). Data are representative of two independent experiments.

(F and G) Schematics of MC38 and B16 *in vivo* experiments from Figure 6. For MC38, mice received one dose of anti-CTLA-4 followed by daily injections of Ctrl or DAHB. For B16, mice received three doses of anti-CTLA-4 with Ctrl- or DAHB-treatment on days not receiving anti-CTLA-4.

Data are mean ± SD. Statistical analysis was performed by paired two-tailed Student’s t test (A, B, C). ns, not significant; *p < 0.05; **p < 0.01; ***p < 0.001; ****p < 0.0001.

**Figure S11. DAHB treatment promotes effector function in human CD8^+^ T cells with metabolic features overlapping those in mouse T cells, related to Figure 7**.

(A) Representative flow cytometry histograms of PD-1 and CD39 surface expression in acutely or chronically stimulated human CD8^+^ T cells treated with Ctrl (NaCl, 20 mM) or DAHB (20 mM). Representative histograms from one donor.

(B) MFI of PD-1 and CD39 from (A); quantification is from triplicate wells from the donor shown in (A). Similar results were obtained from two additional independent donors.

(C) Metabolomics volcano plots from Figure 7D with additional metabolite labeling. Colored dots indicate metabolite categories: creatine (red), redox (purple), pentose phosphate pathway (PPP; orange), glycolysis (blue), and serine/glycine (green). Statistical thresholds are as described in Figure 7D. NADH, reduced nicotinamide adenine dinucleotide; G3P, glyceraldehyde-3-phosphate; F1,6BP, fructose-1,6-bisphosphate; F6P, fructose-6-phosphate; 3PG, 3-phosphoglycerate; R5P, ribose-5-phosphate; PEP, phosphoenolpyruvic acid; Ru5P, ribulose-5-phosphate; 6PG, 6-phosphogluconate.

(D) Ratio of ATP/ADP measured by LC-MS in acutely or chronically stimulated total human T cells treated with Ctrl or DAHB; n = 3 replicates from one donor, with similar results obtained from a second donor.

(E) MFI of TNF-α and IFN-γ production in acutely or chronically stimulated human CD8^+^ T cells treated with Ctrl, DAHB or DAHB + cyclocreatine (DAHB + Cyc); n = 3 replicates from one donor, with similar results obtained from a second donor.

(F) MFI ratio of TNF-α and IFN-γ in DAHB + Cyc compared to DAHB alone from cells in (E); n = 3 replicates from one donor, with similar results obtained from a second donor.

(G) Representative flow cytometry plot of CD8α vs. mTCRβ, confirming comparable 1G4 retroviral transduction efficiency in Ctrl- and DAHB-treated human CD8^+^ T cells used in Figures 7E–7J; similar transduction efficiency was observed in a second independent donor.

Data are mean ± SD. Statistical analysis was performed by one-way ANOVA with Tukey’s multiple comparisons test (B, D, E) or unpaired two-tailed Student’s t test (F). ns, not significant; *p < 0.05; **p < 0.01; ***p < 0.001; ****p < 0.0001.

**Table S1. Oligonucleotides used in this study, related to STAR Methods.**

## EXPERIMENTAL MODEL AND STUDY PARTICIPANT DETAILS

### Mouse strains

C57BL/6J (WT), C57BL/6-Tg(*TcraTcrb*)1100Mjb/J (OT-I), NOD.Cg-*Prkdc*^scid^ *Il2rg*^tm1Wjl^/SzJ (NSG), *Ldhd*-flox, B6.Cg-Tg(*Cd4*-cre)1Cwi/BfluJ (CD4Cre), and B6.Cg-Rag2^tm1.1Cgn^/J (RAG2 KO) mice were maintained under specific pathogen-free conditions. Both male and female mice between 6–12 weeks of age were used for all experiments. C57BL/6J (strain #000664), OT-I (strain #003831), NSG (strain #005557), CD4Cre (strain #022071) and RAG2 KO (strain #008449) were purchased from The Jackson Laboratory. *Ldhd*-flox mice were a gift from the laboratory of Dr. Navdeep Chandel. All animal experiments adhered to policies and practices approved by Memorial Sloan Kettering Cancer Center’s Institutional Animal Care and Use Committee (IACUC) and were conducted in accordance with NIH guidelines for animal welfare (protocol no. 11-03-007). For *in vivo* tumor experiments, mice were euthanized when tumors reached 20 mm in any dimension or if signs of distress were observed. Mice were randomly assigned to experimental groups. No statistical methods were used to predetermine sample sizes, but sample sizes were similar to those reported in previous publications. No blinding was performed.

### Cell lines

MC38 was maintained in-house. B16-F10 (referred to as B16) was purchased from ATCC. A375 and GP2-293 were gifts from Dr. Christopher Klebanoff. E.G7-OVA (EL4-OVA) was a gift from Dr. David Scheinberg. B16, MC38, A375 and GP2-293 were cultured in DMEM High Glucose (MSKCC Media Core) supplemented with 10% FBS, 100 unit/mL penicillin, 100 μg/mL streptomycin. EL4-OVA was cultured in TCM supplemented with 0.4 mg/mL G418. All cell lines were cultured in a 37 °C incubator with 5% CO_­_. Cell lines were verified by expected morphology and growth characteristics and routinely tested to be mycoplasma-free by MycoAlert Mycoplasma Detection Kit (Lonza).

### Mouse T cell isolation and culture

CD8^+^ T cells were isolated from mouse spleen and lymph nodes using Dynabeads Untouched Mouse CD8^+^ Cells Kit (Thermo Fisher Scientific) according to the manufacturer’s instructions. CD8^+^ T cells were seeded at 200,000 cells/well in 96-well plates precoated in 50 μg/mL goat anti-hamster IgG (Jackson Immunoresearch Laboratories) for 3 h at 37 °C. T cells were activated in T cell media (TCM) RPMI 1640 (MSKCC Media Core) supplemented with 10% FBS, 4 mM L-glutamine, 20 mM HEPES, 100 unit/mL penicillin, 100 μg/mL streptomycin, 50 μM β-mercaptoethanol by the addition of 0.5 μg/mL anti-CD3 (Thermo Fisher Scientific), 1 μg/mL anti-CD28 (Thermo Fisher Scientific) and 100 U/mL human IL-2 (Peprotech). For metabolite treatment, metabolites were added either after 2 days (Figure 1B and Figures S1A–S1C) or 1 day (all other figures unless otherwise stated) of activation followed by addition of metabolites as indicated and downstream assays performed at 24 h. For cytokine measurements, cells were restimulated with eBioscience Cell Stimulation Cocktail 1:1000 (Thermo Fisher Scientific) for 3 h. To test for epigenetic effects, cells were washed with PBS after the treatment period and cultured an additional day without metabolites.

### Human T cell isolation and culture

For acute versus chronic stimulation (Figures 7A–7D, S11A–S11F), de-identified healthy donor buffy coats from the New York Blood Center (NYBC) were obtained to isolate human peripheral blood mononuclear cells (PBMCs). Total human T cells were purified from PBMCs using the Dynabeads Untouched Human T cell kit (Thermo Fisher Scientific) following manufacturer’s instructions. T cells were activated with plate-bound 5 μg/mL anti-CD3 and 2 μg/mL anti-CD28 (BioLegend) in RPMI-1640 medium supplemented with 10% FBS, 4 mM L-glutamine, 100 U/mL penicillin-streptomycin and 5 ng/mL each of recombinant human IL-7 and IL-15 (Peprotech) (RPMI-1640 complete medium).

For xenograft and cytotoxicity experiments (Figures 7E–7J), leukopaks were obtained from healthy donors (STEMCELL Technologies). CD8^+^ T cells were isolated by negative selection (STEMCELL Technologies). Isolated CD8^+^ T cells were activated with plate-bound anti-CD3 (5 μg/mL, BioLegend) and anti-CD28 antibodies (2 μg/mL, BioLegend). T cells were cultured in RPMI 1640 with 1% v/v penicillin-streptomycin, 2.5% v/v HEPES, 0.02% v/v gentamicin and 10% v/v heat-inactivated human serum AB (Fisher Scientific). Recombinant human IL-2 (Klebanoff Lab) was added to T cell cultures at 300 IU/mL. Metabolites were added as indicated.

## METHOD DETAILS

### Drug doses

All drug concentrations for cell culture unless otherwise specified in the figure legend are as follows: NaCl also referred to as Ctrl (20 mM), sodium D-lactate (20 mM), DAHB and hydroxybutyrates (20 mM, pH 7.0–7.4 adjusted with NaOH), sodium butyrate (0.5 mM), piericidin (100 nM), antimycin A (10 nM), oligomycin (10 nM), etomoxir (80 µM), teglicar (20 µM), CKi (4 µM), ISRIB (1 µM), Cyclocreatine (10 mM), AU-24118 (1 µM), BRM014 (1 µM), syrosingopine (1 µM), RSL3 (1 µM).

### Flow cytometry in mouse T cells

Cells were harvested from 96-well plates, washed, and surface stained in PBS for anti-CD8a PerCP/Cyanine5.5 (BioLegend) and Live/Dead Fixable Near-IR (Thermo Fisher Scientific) for 15 min at room temperature prior to fixation. Following surface staining, cells were fixed/permeabilized (BD Biosciences) according to the manufacturer’s protocol. Intracellular staining was performed with anti-Perforin, anti-TNF-α, and anti-IFN-γ (BioLegend) for 30 min at room temperature. Cells were washed and resuspended in PBS for flow cytometry on an LSRFortessa (BD Biosciences). Analysis was conducted using FlowJo v10.10.0.

### Cytotoxicity assay for mouse T cells

OT-I T cells were activated and treated with metabolites as described above. On the day of co-culture, EL4-OVA cells were labeled with 2.5 µM CFSE (Thermo Fisher Scientific) for 5 min at room temperature followed by washing once with PBS and plating in 96-well plate at 20,000 cells per well. OT-I CD8^+^ T cells were washed, resuspended in TCM and incubated with tumor cells overnight at the indicated ratio of effector-to-target (E:T) cells in the figure legend in triplicate. The following day, co-culture wells were washed and resuspended in PBS and DAPI (1 μg/mL). Cytotoxicity was calculated as 100 − % CFSE^+^DAPI^−^ (equivalent to % CFSE^+^DAPI^+^).

### CRISPR in CD8^+^ T cells

Naïve CD8^+^ T cells were isolated from spleen and lymph nodes by FACS sorting on CD8^+^CD62L^+^CD44^lo^ (Thermo Fisher Scientific). Electroporation of naïve CD8^+^ T cells was performed with the Amaxa P3 Primary Cell 4D-Nucleofector X kit. For sgRNA/Cas9 RNP complex formation, 1 µL of Cas9 protein (10 mg/mL, IDT) was mixed with 1 µL of sgRNA (0.3 nmol/µl, GenScript) in nuclease-free water to a final volume of 5 µl, followed by incubation for 10 min at room temperature. The freshly isolated CD8^+^ T cells (3 × 10⁶ cells) were resuspended in buffer P3 with supplement 1 according to the manufacturer’s instructions (Lonza), combined with 5 µl of preassembled sgRNA/Cas9 RNP, and transferred into a Nucleocuvette Vessel. The mixture was electroporated in the Nucleofector X unit using the pulse code DN-100. The cells were subsequently transferred to pre-warmed TCM and activated as described in the T cell culture methods section. Two days later, the cells were treated with 20 mM NaCl or DAHB overnight. The following day, the cells were restimulated with 50 ng/mL PMA (Selleckchem), 0.5 µg/mL ionomycin (STEMCELL Technologies) and Golgiplug protein transport inhibitor 1:1000 (BD Biosciences) for 3 h. An aliquot was taken for western blotting of KO efficiency, and the remainder was analyzed for cytokine production by flow cytometry.

### Western blot

CD8^+^ T cells were lysed directly in 1× SDS loading buffer and loaded onto polyacrylamide gels (Figures S5A, S5B, and S5J). Proteins were then transferred to PVDF membranes by wet transfer on ice at 400 mA for 45 min. Blots were blocked in Intercept (TBS) blocking buffer (LI-COR) and then incubated with primary antibody at 4 °C overnight. The next day, blots were washed with TBST (TBS + 0.1% Tween-20) and stained with fluorescently conjugated secondary antibodies (LI-COR) at 1:10,000 dilution in TBS blocking buffer for 1 h at room temperature and imaged in the 680- and 800-nm channels on the Odyssey CLx Imaging System (LI-COR). Antibodies: anti-HADHB (Proteintech), anti-CPT1A (Abcam), anti-AMPKα (Proteintech), and anti-β-actin (Proteintech).

For all other western blots, CD8^+^ T cells were lysed in RIPA lysis buffer (Sigma-Aldrich) supplemented with cOmplete protease inhibitor cocktail (Sigma-Aldrich) and phosphatase inhibitor cocktail (Thermo Fisher Scientific). BCA assay (Thermo Fisher Scientific) was used to determine protein concentrations followed by loading onto gradient polyacrylamide gels. Proteins were then transferred to nitrocellulose membranes and blocked in 5% milk. Immunoblotting occurred with primary antibody overnight followed by washing in TBST and staining with secondary HRP-conjugated antibody. Imaging was performed with the Digital ECL Substrate (Kindle Biosciences) and ChemiDoc MP Imaging System (Bio-Rad). Antibodies: anti-LDHA, anti-PBRM1, anti-BRG1, anti-BRM, anti-RPS3, anti-Phospho-AMPKα (Thr172), anti-AMPKα, anti-ATF4, anti-CHOP, anti-Histone H3, anti-H3K27ac (all Cell Signaling Technology), anti-H3K9ac (Thermo Fisher Scientific), anti-CKB (Santa Cruz Biotechnology).

### Intracellular acidification measurements

Activated CD8^+^ T cells were loaded with pHrodo Red AM (1:2000) by incubation at 37 °C for 30 min in flow media (FM): XF RPMI medium (Agilent), 20 mM HEPES, 10 mM sodium bicarbonate, 10 mM glucose, 2 mM L-glutamine, 50 μM BME, 500 μM L-alanine. Following loading, cells were resuspended in FM with DAPI (1 μg/mL). For syrosingopine (Syro) experiments, cells were incubated with DMSO (vehicle) or Syro (1 μM) for 10 min prior to addition of metabolites. Samples were analyzed immediately after addition of metabolites (20 mM) or after 30 min incubation (Figure S2G). The MFI ratio was calculated by normalizing to the MFI in Ctrl treatment defined as 20 mM NaCl (MFI_­_/ MFI_­_).

### Lipid peroxidation

To measure lipid peroxidation, CD8^+^ T cells treated with Ctrl or DAHB were stained with BODIPY 581/591 C11 (Thermo Fisher Scientific) in FM for 20 min in the cell culture incubator. For positive control, RSL3 (1 μM) was added 10 min prior to BODIPY. After the staining incubation, cells were resuspended in FM with DAPI for flow cytometry. Lipid peroxidation was measured by the MFI ratio of FITC/PE.

### Low-glucose media T cell culture

Glucose-free RPMI was used as base media and prepared with supplements as described for standard TCM. After addition of FBS, glucose concentration was 1.3 mM (low-glucose media) as measured by glucometer (Chemglass). CD8^+^ T cells were activated in low-glucose media concurrently with either Ctrl or DAHB at 20 mM for 48 h. Following incubation, cells were counted by hemocytometer and fixed for flow cytometry as described previously.

### Puromycin incorporation assay

Activated CD8^+^ T cells were treated with Ctrl, DAHB, DAHB + oligomycin (O), or DAHB + O + ISRIB overnight. The next day, an untreated control well was pre-treated with cycloheximide (CHX) at 50 µg/mL for 20 min, followed by the addition of puromycin (10 μg/mL) to all wells for 15 min in the cell culture incubator. Cells were quickly detached from wells and washed with PBS followed by lysis in RIPA buffer (Sigma-Aldrich) for western blot using anti-puromycin (Kerafast) antibody.

### Chromatin immunoprecipitation and qPCR

Chromatin was isolated from approximately 3 × 10^6^ CD8^+^ T cells treated with Ctrl, DAHB or DAHB+O (oligomycin) per immunoprecipitation. Cells were harvested, washed with PBS, and cross-linked with freshly prepared 1% paraformaldehyde for 5 min at room temperature with gentle rotation. Glycine was added to quench at a final concentration of 125 mM and samples were incubated for 5 min at room temperature. Cells were washed once with cold PBS and pelleted. Chromatin extraction was performed by resuspending cells in 1 mL LB1 (50 mM HEPES, 140 mM NaCl, 1 mM EDTA, 10% glycerol, 0.5% NP-40, 0.25% Triton X-100 and 1× cOmplete™, EDTA-free Protease Inhibitor Cocktail (Sigma-Aldrich)) and rotated at 4 °C for 10 min. Samples were centrifuged, resuspended in 1 mL LB2 (10 mM Tris-HCl pH 8.0, 200 mM NaCl, 1 mM EDTA, 0.5 mM EGTA and 1× protease inhibitor cocktail) and rotated at 4 °C for 10 min. Finally, samples were centrifuged, resuspended in 1 mL LB3 (10 mM Tris-HCl pH 8.0, 100 mM NaCl, 1 mM EDTA, 0.5 mM EGTA, 0.1% sodium deoxycholate, 0.5% *N*-lauroylsarcosine, 1% Triton X-100 and 1× protease inhibitor cocktail) and homogenized by passing twice through a 27-gauge needle. Chromatin extracts were sonicated in a Bioruptor Plus sonication device (Diagenode) for 80 cycles of 30 s on/30 s off on high intensity.

For immunoprecipitation, the lysates were incubated with 8 μL of anti-H3K9ac (Active Motif) bound to 50 μL protein A Dynabeads (Thermo Fisher Scientific) and incubated overnight at 4 °C with 5% of lysate kept as input DNA. Magnetic beads were subsequently washed twice each with low-salt buffer (150 mM NaCl, 0.1% SDS, 1% Triton X-100, 1 mM EDTA and 50 mM Tris-HCl), followed by high-salt buffer (500 mM NaCl, 0.1% SDS, 1% Triton X-100, 1 mM EDTA and 50 mM Tris-HCl), LiCl buffer (150 mM LiCl, 0.5% sodium deoxycholate, 0.1% SDS, 1% NP-40, 1 mM EDTA and 50 mM Tris-HCl), followed by one wash with TE buffer (1 mM EDTA and 10 mM Tris-HCl). Beads were resuspended in elution buffer (1% SDS, 50 mM Tris-HCl pH 8.0, 10 mM EDTA and 200 mM NaCl) and incubated for 30 min at 65 °C. After magnetic separation, the eluate was reverse-cross-linked overnight at 65 °C. Input DNA samples were also resuspended in an equal volume of elution buffer and reverse-cross-linked overnight. Samples were treated with RNase A (Qiagen) for 1 h at 37 °C and with Proteinase K (Qiagen) for 1 h at 55 °C and the DNA was recovered using a QIAquick PCR Purification Kit (Qiagen).

A 1:100 dilution of each sample was used for qPCR analysis. ChIP-qPCR was performed with the QuantStudio 7 Flex Real-Time PCR System (Thermo Fisher Scientific) using 40 cycles of amplification with SsoAdvanced Universal SYBR Green Supermix (Bio-Rad) according to the manufacturer’s instructions.

### HDAC activity assay

Class I/II HDAC activity was measured using the HDAC-Glo™ I/II Assay (Promega). Ctrl, LAHB, DAHB, LBHB, DBHB (each at 20 mM) or butyrate (0.5 mM) treatments were added to activated CD8^+^ T cells (25,000 cells per well in a 96-well plate) in triplicate wells (Figure S6A). After incubation for 1 h at 37 °C, the HDAC-Glo I/II reagent was added for 30 min at room temperature followed by luminescence acquisition using a Cytation 3 plate reader (Agilent).

Cell-free recombinant HDAC activity assays (HDAC1 Fluorogenic Assay Kit and HDAC2 Fluorogenic Assay Kit; BPS Bioscience) were performed according to the manufacturer’s instructions. Briefly, recombinant mouse HDAC1 or HDAC2 (supplied with kits) were incubated with Ctrl (NaCl, 10 mM), DAHB (10 mM) or TSA (2 μM) for 30 min at 37 °C (Figure S6B). Next, the HDAC developer was added for 15 min at room temperature. The plate was read on a Cytation 3 plate reader (Agilent) with excitation/emission = 365/450 nm.

### Histone modification profiling

Histones from activated CD8^+^ T cells treated with Ctrl or DAHB were isolated according to the manufacturer’s instructions (Epigentek). The isolated histones were propionylated with propionic anhydride and acetonitrile in a ratio of 1:3 (v/v). The propionylated histones were digested with trypsin (Promega) for 12 h followed by desalting with StageTips. The digests were analyzed using a Thermo Fisher Scientific EASY-nLC 1200 system coupled online to a Fusion Lumos mass spectrometer (Thermo Fisher Scientific). Buffer A (0.1% FA in water) and buffer B (0.1% FA in 80% ACN) were used as mobile phases for gradient separation. A 75 µm × 15 cm chromatography column (ReproSil-Pur C18-AQ, 3 µm, Dr. Maisch GmbH, Germany) was packed in-house for peptide separation. Peptides were separated with a gradient of 5–33% buffer B over 45 min, 33–100% B over 10 min at a flow rate of 400 nL/min. The Fusion Lumos mass spectrometer was operated in a data-independent acquisition (DIA) mode. MS1 scans were collected in the Orbitrap mass analyzer from 300–1100 m/z at 60,000 resolution. The instrument was set to select precursors in 50 × 16 m/z wide windows with 1 m/z overlap from 300–1100 m/z for HCD fragmentation. The MS/MS scans were collected in the Orbitrap at 15,000 resolution. The raw data were processed with EpiProfile v2.1 for identification and quantification of histone peptides. Heatmaps were generated by Z-score normalization of relative abundances for each histone modification across three replicates and visualized using the pheatmap package in R.

### Short-chain Acyl-CoA Analysis

For Acyl-CoA profiling, 6 × 10^6^ CD8^+^ T cells were activated and treated with Ctrl or DAHB overnight. Cells were washed with PBS and snap frozen in liquid nitrogen. Short-chain acyl-CoAs were extracted from cells using pre-cooled 90% methanol. Samples were centrifuged at 4 °C for 20 min at 20,000 × g. The supernatants containing short-chain acyl-CoAs were cleaned up with a Sep-Pak column and then dried down. Targeted LC/MS analyses were performed on a Q Exactive Orbitrap mass spectrometer (Thermo Fisher Scientific) coupled to a Vanquish UPLC system (Thermo Fisher Scientific). The Q Exactive operated in positive mode. An Imtakt Cadenza CD-C18 column (2.0 mm inner diameter × 150 mm) was used to separate metabolites. The flow rate was set at 150 μL/min. Buffers consisted of 100% methanol for mobile phase B, and 5 mM CH_­_COONH_­_in water for mobile phase A. The gradient ran from 2% to 15% B in 5 min, changed to 95% B in 3 min, and followed by a wash with 95% B and re-equilibration at 85% B. The MS data were processed using Xcalibur v4.1 (Thermo Fisher Scientific). Metabolites were identified based on exact mass within 5 ppm and standard retention times. Relative metabolite quantitation was performed based on the peak height of each metabolite. Statistical significance was assessed by one-sample t test with hypothetical value 0.

### Immunofluorescence

CD8^+^ T cells were activated in 12-well plates at a density of 3 × 10^6^ cells per well. Following treatment with Ctrl, DAHB, DAHB + O or DAHB + Cyc, cells were transferred into 2 mL Eppendorf tubes and centrifuged at 500 × g for 5 min. Supernatant was aspirated and cell pellets were fixed with pre-warmed 4% paraformaldehyde (Thermo Fisher Scientific) for 10 min. Cells were then centrifuged and washed once with PBS before permeabilization using 0.2% Triton X-100 at room temperature for 10 min. Cells were centrifuged and washed once with PBS again, followed by blocking with PBS containing 10% normal goat serum for 30 min at room temperature.

After blocking, cells were then centrifuged to remove blocking buffer and incubated with primary anti-CKB antibody at 1:150 (Santa Cruz Biotechnology) ± anti-Lamin B1 antibody at 1:150 (Abcam) for overnight incubation at 4 °C. The following day, cells were centrifuged and washed thrice with PBS before incubation with goat anti-mouse secondary antibody at 1:500 ± goat anti-rabbit secondary antibody at 1:500 (Sigma-Aldrich) ± DAPI (MilliporeSigma) for 1 h at room temperature. After secondary antibody staining, cells were washed thrice with PBS and resuspended in a 1:1 PBS:SlowFade Gold Antifade Mountant (Thermo Fisher Scientific) solution. Cells were then transferred and plated onto µ-Slide 8 Well high Glass Bottom slides (ibidi). To cluster cells onto the coverslip for imaging, slides were centrifuged for 5 min at 500 × g before imaging. For standard confocal immunofluorescence, slides were imaged using the Nikon Eclipse Ti2 inverted microscope equipped with the Yokogawa CSU-W1 SoRa spinning disk confocal unit. The Nikon SoRa microscope has a Photometrics Prime BSI sCMOS camera; 4 laser lines – 405-, 488-, 561-, and 640-nm for fluorescence imaging in the blue, green, red and far-red channels combined with their respective standard emission filter sets. Images were acquired using the CFI Plan Apochromat Lambda (100×, 1.45NA, oil) and NIS-Elements software. Z-stacks were taken with the blue and green channels at 0.2 μm step size, for 30–40 steps to fully sample from the basal to the apical region.

Raw z-stack images were deconvolved in AutoQuant X3 (Media Cybernetics) using the 3D deconvolution plugin. Under deconvolution settings, a total of 15 iterations and 15 save intervals were repeated and the output was generated as 16-bit floating point IMS files for further processing with ImageJ/Fiji to generate maximum intensity projected micrographs.

### STORM imaging

Cells were prepared as described above with the exclusion of DAPI staining. Images were acquired using an inverted ONI Nanoimager microscope (Oxford Nanoimaging Ltd., Oxford, UK) equipped with a 100× NA 1.4 oil immersion objective lens and an ORCA-Flash 4.0 camera (Hamamatsu, Shizuoka Japan) with a pixel size of 6.5×6.5 μm^2^. Fluorophores were excited using lasers with wavelengths of 488- and 561-nm. A 405-nm laser was used to promote fluorophore blinking. Total internal reflection fluorescence illumination was employed to capture cell images. Channel mapping calibration was performed at the beginning of each imaging session using a calibration slide provided by ONI. All images were acquired in ONI storm buffer. Image series were acquired for each field of view with an exposure time of 30 ms per frame for a total of 10,000 frames per channel. Localization, filtering, post-processing and image acquisition were performed in the Nanoimager software. Representative images are shown in Figure S9G from two independent experiments with similar results.

### Proximity ligation assay

For proximity ligation assay (PLA), the Duolink *In Situ* Red Starter Kit Mouse/Rabbit (Sigma-Aldrich) was used. Cells were fixed with pre-warmed 4% PFA for 10 min at room temperature in 1.5 mL Eppendorf tubes. Samples were then permeabilized using 0.2% Triton X-100 for 5 min, centrifuged at 500 × g for 5 min to remove supernatant, washed with PBS and incubated with 10% Normal Goat Serum in PBS as blocking buffer for 30 min. Cells were then centrifuged and incubated with primary antibodies mouse anti-CKB at 1:150 (Santa Cruz Biotechnology) and rabbit anti-Lamin B1 at 1:150 (Abcam) in Duolink antibody diluent overnight at 4 °C.

The following day, cells were centrifuged and washed twice with PBS followed by incubation with Duolink PLA probes PLUS and MINUS secondary antibodies for 1 h at 37 °C. Cells were then centrifuged and washed twice with Duolink wash buffer A and incubated with Duolink ligase for 30 min at 37 °C to complete the ligation step. After ligation, cells were washed twice with Duolink wash buffer A and incubated with Duolink polymerase for 100 min at 37 °C for amplification. After amplification, the cells were washed with Duolink wash buffer B and stained with 1 µg/mL DAPI (Sigma-Aldrich) in PBS for 1 h. After staining, cells were washed with PBS and resuspended in 100 µL of PBS. The cell suspensions were then transferred onto µ-Slide 8 Well high Glass Bottom plates (ibidi) containing 100 µL of SlowFade gold Antifade Mountant (Thermo Fisher Scientific) in each well. Plates were centrifuged at 500 × g for 5 min to cluster cells before imaging on the Nikon Eclipse Ti2 with Yokogawa CSU-W1 SoRa spinning disk confocal. Nuclear puncta were counted at 100× magnification with the same settings across both Ctrl- and DAHB-treated CD8^+^ T cells images using ImageJ.

### LC-MS metabolite analysis in mouse T cells

CD8^+^ T cells were activated in 12-well plates at a density of 3 × 10^6^ cells per well. Following overnight metabolite and inhibitor treatment, cells were quickly washed with 0.9% NaCl solution on ice and metabolism was quenched by the addition of 1 mL of 80:20 methanol:water, followed by storage at −80 °C overnight. The methanol-extracted metabolites were cleared by centrifugation and the supernatant was dried in a vacuum evaporator (Genevac EZ-2 Elite) for 5 h prior to analysis by LC-MS.

For nutrient tracing experiments, cells were incubated in 12-well plates as above and after treatment with metabolites, the media were removed and replaced with RPMI 1640 supplemented with 10% dialyzed FBS, 2 mM L-glutamine, 20 mM HEPES, 100 unit/mL penicillin, 100 μg/mL streptomycin and 50 µM β-mercaptoethanol. For tracing [U-^13^C]glutamine, glutamine-deficient RPMI was used and for [U-^13^C]glucose, glucose-deficient RPMI was used. Stable isotope tracers were obtained from Cambridge Isotope Laboratories or Sigma-Aldrich. The following concentrations and incubation times at 37 °C and 5% CO_­_were used for each tracer: [U-^13^C]glucose (10 mM, 30 min), [U-^13^C]glutamine (2 mM, 5 h), [U-^13^C]acetate (2 mM, 5 h), [U- ^13^C]butyrate (0.5 mM, 5 h), [U-^13^C]DAHB (5 mM, 5 h), [U-^13^C]octanoate (1 mM, 5 h), [U-^13^C]nonanoate (1 mM, 5 h), [U-^13^C]palmitate (200 µM, 5 h), [U-^13^C]oleate (200 µM, 5 h). Cells were extracted as described above and submitted for analysis by LC-MS.

LC-MS analyses were performed on a Q Exactive Orbitrap mass spectrometer (Thermo Fisher Scientific) coupled to a Vanquish UPLC system (Thermo Fisher Scientific). The Q Exactive operated in polarity-switching mode. A Sequant ZIC-HILIC column (2.1 mm inner diameter × 150 mm, Merck) was used for separation of metabolites. Flow rate was set at 150 μL/min. The column temperature was 30 °C. Buffers consisted of 100% acetonitrile for mobile phase B, and 0.1% NH_­_OH/20 mM CH_­_COONH_­_in water for mobile phase A. The gradient ran from 85% to 30% B in 20 min followed by a wash with 30% B and re-equilibration at 85% B. For untargeted metabolomics, the MS data were processed using Compound Discoverer (Thermo Fisher Scientific). The Human Metabolite Database (HMDB) and an in-house metabolite library containing standard retention time information were searched for metabolite identification. Three levels of metabolite identification were reported: 1) identified compounds: definitive identification based on the mass (within 5 ppm) and retention time of authentic chemical standards; 2) putatively annotated compounds by searching HMDB (mass tolerance 5 ppm); 3) compounds with predicted chemical composition based on mass. Relative metabolite quantitation was performed based on peak area for each metabolite. For stable isotope tracing, raw data were processed using El-MAVEN v0.12.0. Metabolites and their ^13^C isotopologues were identified based on exact mass (±5 ppm) and standard retention times. Volcano plots were generated in R using log2 fold change and Benjamini-Hochberg correction for p values derived from two-sided Welch’s t-tests (performed per metabolite). Significance was defined as FDR < 0.05 and 1.25-fold change (|log2FC| > log2(1.25)).

### [U-^13^C]DAHB synthesis

The synthesis commenced from the acetylation of Evans oxazolidinone with [U-^13^C]butyric acid (Cambridge Isotope Laboratories), which ensures the stereochemical outcome of the α-hydroxyl group to the desired configuration (D-isomer)^77^. Non-aqueous reactions were performed under an atmosphere of argon, in flame-dried glassware. All reactions were monitored by thin-layer chromatography. ^1^H NMR spectra were recorded on Bruker Avance 400. High resolution mass spectra were acquired on a UPLC-QTOF Triple TOF 6600+ mass spectrometer. Reactions were carried out according to known procedures^78^. The final product was lyophilized to produce a white powder (18.3 mg, calculated mass purity ∼40.5%). Structural identity and purity were confirmed by nuclear magnetic resonance spectroscopy and high-resolution mass spectrometry.

### RNA Isolation and RT-qPCR

RNA was isolated using the RNeasy Plus Micro Kit (Qiagen) according to the manufacturer’s instructions. Reverse transcription was performed with iScript cDNA Synthesis Kit (Bio-Rad) using oligo(dT) and random hexamers. RT-qPCR was performed with SsoAdvanced Universal SYBR Green Supermix (Bio-Rad) in triplicate using the instrument QuantStudio 7 Flex Real-Time (Thermo Fisher Scientific) with 40 cycles of amplification. Each transcript was normalized to *Rpl13a* and relative expression was calculated using the 2^−ΔΔCT^ method^79^.

### RNA-seq

RNA was prepared in triplicate and submitted for poly(A) enrichment and library preparation followed by paired-end Illumina sequencing by Azenta Life Sciences or MSKCC Integrated Genomics Core. For Azenta Life Sciences (Figure 1C): RNA sequencing reads were trimmed for adapter sequences and low-quality bases using Trimmomatic v0.36, and aligned to the Mus musculus GRCm38 reference genome using STAR v2.4. Only uniquely aligned reads were retained. Gene-level read counts were generated with featureCounts v1.5.2 using exon regions from the Ensembl annotation. If applicable, strand-specific counting was performed. Normalization and differential expression were evaluated with DESeq2 using default parameters. For MSKCC Integrated Genomics Core (Figures 5G and S7A–S7C): RNA sequencing reads were 3′ trimmed for base quality 15 and adapter sequences using TrimGalore v0.4.5 and aligned to mouse assembly mm10 with STAR v2.4 using default parameters. Data quality and transcript coverage were assessed using the Picard tool CollectRNASeqMetrics (https://broadinstitute.github.io/picard/). Read count tables were generated with HTSeq v0.9.1. Normalization and differential expression were evaluated with DESeq2 using default parameters and outliers were assessed by sample grouping in principal component analysis.

To generate volcano plots, RNA-seq results were imported into R. Genes with missing or infinite values were removed. Plots were generated using ggplot2 with Benjamini-Hochberg corrected significance p < 0.05 and fold change threshold at 2-fold (Figure 1C) or 1.25-fold (Figure 5G). Extremely small p-values were capped at 10^-300^ to avoid axis distortion. For K-means clustering, normalized gene expression data were Z-score scaled and subjected to principal component analysis. Principal components explaining ≥90% of total variance were retained and used as input (Lloyd algorithm). Genes were grouped by clusters and visualized in a heatmap using the pheatmap R package.

### Proteomics

CD8^+^ T cells were activated on 10 cm dishes at a density of 10 × 10^6^ cells/plate and treated with Ctrl or DAHB overnight in triplicate. Cells were harvested the next day and washed with ice-cold PBS followed by flash freezing in liquid nitrogen. Cell pellets were resuspended in 100 μL of 8 M urea, 50 mM HEPES (pH 8.5). Reduction was performed by adding tris-(2-carboxyethyl)phosphine (TCEP) to a final concentration of 5 mM, followed by incubation for 30 min at 25°C with shaking at 1,000 rpm on a Thermomixer. Free cysteine residues were alkylated with 2-iodoacetamide (IAA) to a final concentration 10 mM, and the samples were incubated for 30 min at 25 °C in the dark. The reaction was quenched by adding DTT to a final concentration of 5 mM and incubating for 15 min at 25 °C. For enzymatic digestion, Lys-C was added at a 1:100 enzyme-to-protein ratio, and the mixture was incubated for 1 h at 25 °C with shaking at 1,150 rpm. The urea concentration was reduced by adding 100 μL of 50 mM ammonium bicarbonate (ABC). Trypsin was then added at a 1:100 enzyme-to-protein ratio, and the samples were incubated overnight at 37 °C with shaking at 1,150 rpm.

The following day, an additional 500 ng of trypsin was added, and the samples were incubated for 2 h at 37 °C with shaking at 1,150 rpm. The samples were briefly centrifuged for 1 min, and the digests were acidified to pH <3 by adding 50% trifluoroacetic acid (TFA). Peptides were desalted using Sep-Pak C18 cartridges (Waters). Briefly, the cartridges were conditioned sequentially with (i) methanol, (ii) 70% acetonitrile (ACN)/0.1% TFA, and (iii) 5% ACN/0.1% TFA twice. After conditioning, the acidified peptide solution was loaded onto the cartridges, and the stationary phase was washed twice with 5% ACN/0.1% formic acid (FA). Peptides were eluted twice with 70% ACN/0.1% FA, then dried under vacuum using a SpeedVac centrifuge. The dried peptides were reconstituted in 12 μL of 0.1% FA, sonicated, and transferred to an autosampler vial. Peptide yield was quantified using a NanoDrop spectrophotometer.

Peptides were separated on a 25 cm column with a 75 μm inner diameter and 1.7 μm particle size, packed with C18 stationary phase (IonOpticks Aurora 3), using a gradient from 2% to 35% buffer B over 90 min, followed by an increase to 95% buffer B for 7 min (buffer A: 0.1% FA in HPLC-grade water; buffer B: 99.9% ACN, 0.1% FA) with a flow rate of 300 nL/min on a NanoElute2 system (Bruker).

MS data were acquired on a TimsTOF HT mass spectrometer (Bruker) equipped with a Captive Spray source (Bruker) using a data-independent acquisition PASEF method (dia-PASEF). The mass range was set from 100 to 1700 m/z, and the ion mobility range from 0.60 V·s/cm^2^ (collision energy 20 eV) to 1.6 V·s/cm^2^ (collision energy 59 eV), a ramp time of 100 ms, and an accumulation time of 100 ms. The dia-PASEF settings included a mass range of 400.0 to 1201.0 Da, a mobility range of 0.60–1.60, and an estimated cycle time of 1.80 s. The dia-PASEF windows were set with a mass width of 26.00 Da, a mass overlap 1.00 Da, and 32 mass steps per cycle.

Raw data files were processed using Spectronaut version 18.4 (Biognosys) and searched with the Pulsar search engine against a UniProt mouse protein database downloaded on 2022/09/26 (94,615 entries). Cysteine carbamidomethylation was specified as a fixed modification, while methionine oxidation, acetylation of the protein N-terminus and deamidation (NQ) were set as variable modifications. A maximum of two trypsin missed cleavages were permitted. Searches used a reversed sequence decoy strategy to control peptide false discovery rate (FDR) and 1% FDR was set as the threshold for identification. After filtering for non-missing intensity values across all three replicates and excluding proteins with peptide count <5, the volcano plot was generated in R. Log2-transformed protein intensities were averaged per condition, and the log2 fold change was computed for each protein (threshold 1.5-fold change). Statistical significance between conditions was determined using Welch’s t-test (threshold p <0.05).

### Seahorse bioenergetics analysis

Oxygen consumption rate (OCR) and extracellular acidification rate (ECAR) were measured using an XFe96 Extracellular Flux Analyzer (Agilent) according to the manufacturer’s instructions. Seahorse microplates were precoated with poly-D-lysine (Thermo Fisher Scientific) at 0.1 mg/mL for 30 min at room temperature, washed twice with PBS, and stored overnight at 4 °C. Assays were performed in Seahorse medium (SM), consisting of XF RPMI (Agilent) supplemented with 10 mM glucose, 2 mM glutamine, 50 μM β-mercaptoethanol, and 500 μM L-alanine. Activated T cells were plated at 1.8 × 10^5^ cells per well in 180 μL SM and incubated at 37 °C in a non-CO_­_incubator during sensor cartridge equilibration.

For mitochondrial stress assays, T cells activated overnight were plated either in SM alone for acute metabolite injection during the assay or in SM containing the indicated metabolites and inhibitors for continuous exposure throughout equilibration and measurement. OCR was measured at baseline and after sequential injection of oligomycin (1 μM), FCCP (1 μM), and rotenone plus antimycin A (1 μM each), with metabolites or Ctrl (NaCl, 20 mM) added either acutely as the first injection or present throughout the assay as indicated. For prolonged-treatment experiments, activated CD8^+^ T cells were cultured for an additional 48 h in the indicated conditions prior to assay and then replated in SM containing the same treatment.

For glycolytic stress assays, activated T cells were plated in glucose-free SM containing the indicated treatment conditions. ECAR was measured at baseline and after sequential injection of glucose (10 mM), oligomycin (1 μM), and 2-deoxyglucose (50 mM).

### Continuous oximetry

To measure oxygen consumption during cell culture, 200,000 CD8^+^ T cells were activated overnight as described previously in 96-well plates, followed by treatment with NaCl (Ctrl) or DAHB at 20 mM. The plate lid was then replaced with the Resipher (Lucid Scientific) sensing lid and connected to the Resipher hub to obtain 24 h of continuous oxygen consumption readings.

### ATAC-seq and chromatin accessibility

A total of 100,000 CD8^+^ T cells per replicate were harvested and processed using the ATAC-seq Kit (Active Motif) according to the manufacturer’s instructions. Libraries were sequenced on Illumina platforms to obtain 40–50 million reads per sample. FASTQ files were uploaded to the Galaxy platform and processed on the public server at usegalaxy.org^80^. BAM files were generated using Bowtie2, aligned to mm10 with the very sensitive local preset. BigWig files for track visualization were generated using the bamCoverage tool with normalization set to 1× genome coverage. The track locations shown are as follows: *Cd8a*, Chr6(71,364,427-71,388,171); *Prf1*, Chr10(61,288,751-61,313,684); *Ifng*, Chr10(118,432,046-118,454,894); *Tnf*, Chr17(35,198,367-35,203,007). The total signal was calculated along these windows for each replicate BigWig using rtracklayer in R.

### Pharmacokinetics of LAHB and DAHB

WT mice were injected intraperitoneally with LAHB or DAHB (1.65 g/kg) and blood was collected from two separate mice per treatment per time point into EDTA-coated tubes at the following intervals: 0, 15, 90, and 240 min. Plasma was collected by centrifugation at 4 °C (3,000 × g for 10 min). Plasma metabolites were extracted with 80% methanol containing [U-¹³C]DAHB as an internal standard. The extracts were dried and reconstituted in water before analysis by LC–MS using an Orbitrap Exploris 240 mass spectrometer coupled to a Vanquish UHPLC system (Thermo Fisher Scientific). The mass spectrometer was operated in negative ion mode. Metabolites were separated on a Sequant ZIC-HILIC column (2.1 mm × 150 mm, Merck) at a flow rate of 150 μL/min and a column temperature of 30 °C. Mobile phase A consisted of 0.1% NH₄OH with 20 mM ammonium acetate in water; mobile phase B consisted of 100% acetonitrile. The chromatographic gradient decreased from 85% to 30% B over 20 min, followed by a wash at 30% B and re-equilibration at 85% B. MS data were processed using XCalibur 4.1 (Thermo Fisher Scientific) to obtain metabolite signal intensities. Relative quantitation of α-ketobutyrate (AKB) was determined based on exact mass within 5 ppm and standard retention times. Absolute quantitation of α-hydroxybutyrate (AHB) was performed by normalizing analyte signal intensity to the mean [U-¹³C]DAHB internal standard intensity across all non-baseline samples. One-phase exponential decay curves (plateau constrained to zero) were fitted in GraphPad Prism v11.0.0 to determine metabolite half-lives.

### *In vivo* allograft tumor models

B16 melanoma or MC38 colorectal adenocarcinoma tumor cells were resuspended in PBS and implanted subcutaneously in the backs of WT or RAG2 KO mice (250,000 cells/mouse). When tumors became palpable (5–7 days later), mice were randomly separated into two cohorts to receive intraperitoneal (i.p.) injection of anti-CTLA-4 (200 μg). For B16, anti-CTLA-4 injections occurred every other day (three injections) or once for MC38. Concurrently, mice received i.p. injection of Ctrl (NaCl) or DAHB (1.65 g/kg) daily on days not receiving anti-CTLA-4. For adoptive transfer, EL4-OVA (1 × 10^6^ cells/mouse) tumors were implanted subcutaneously into the backs of RAG2 KO mice. OT-I CD8^+^ T cells were activated *ex vivo* with anti-CD3/CD28 and treated with DAHB (20 mM) or NaCl (Ctrl, 20 mM) overnight. The next day, cells were washed with PBS and 500,000 OT-I CD8^+^ T cells were injected i.p. into tumor-bearing mice, 5 days after implantation. Tumor size (mm^2^, maximal length × width) was measured every 2–3 days with calipers.

### T cell mixing for intratumoral analysis

RAG2 KO mice were implanted with EL4-OVA tumors as described previously. One week after implantation, OT-I CD8^+^ T cells were activated and treated with NaCl (Ctrl) or DAHB. The following day, Ctrl- or DAHB-treated T cells were labeled with CFSE at 2.5 μM or CellTrace Far Red at 2.5 μM (Thermo Fisher Scientific), respectively. After 10 min incubation at room temperature, cells were washed three times with PBS, counted and mixed at 1:1 ratio. Cells were then injected subcutaneously, adjacent to the EL4-OVA tumors prior to tumor harvesting the following day.

### Tumor infiltrating lymphocyte isolation

Tumors were excised and mechanically dissociated with scalpels. Single-cell suspensions were obtained by passing through a 40 μm cell strainer. Red blood cells were lysed with ACK Lysing Buffer (Thermo Fisher Scientific). Cells were then washed, resuspended in TCM and restimulated with Cell Stimulation Cocktail (eBioscience) for 4 h. Cells were subsequently fixed, permeabilized and intracellularly stained for cytokine expression by flow cytometry as described previously. For CKB staining, after fixation and permeabilization, anti-CKB (Santa Cruz Biotechnology) was added at a 1:100 dilution for 30 min at room temperature followed by a PBS wash. Next, anti-mouse PE/Cyanine7 secondary antibody (BioLegend) was added at 1:1000 for 30 min, washed, and resuspended in PBS for flow cytometry.

### Human T cell exhaustion model

After 48 h of activation with anti-CD3/CD28 in RPMI-1640 complete medium, T cells were plated at 1 × 10^6^ cells/mL with (chronic stimulation) or without (acute stimulation) plate-bound anti-CD3 (5 μg/mL) and treated with 20 mM NaCl (Ctrl) or 20 mM DAHB as previously described^76^. Cells were passaged every 2 days with fresh treatments into new plate-bound anti-CD3 plates until day 8 post activation. For the DAHB+Cyc condition, 10 mM cyclocreatine was added for the final 48 h (days 6–8). Flow cytometry and metabolomics analyses were conducted on day 8.

### Flow cytometry of human T cells

Cytokine expression of acutely and chronically stimulated T cells treated with 20 mM NaCl (Ctrl), 20 mM DAHB and/or 10 mM cyclocreatine (Cyc) was assessed by flow cytometry on day 8 post activation. T cells were treated with Cell Activation Cocktail for 1 h, followed by Brefeldin A (both BioLegend) treatment for 3 h. Cells were harvested, washed with cold PBS and incubated for 10 min with a solution containing Human TruStain FcX Fc Receptor Blocking Solution (BioLegend) and the viability dye Ghost Dye Violet 510 (Cytek). Thereafter, cells were washed with FACS buffer (2% FBS in PBS) and stained for 45 min at 4 °C with antibodies targeting surface markers using BD Horizon Brilliant Stain Buffer (BD Biosciences). Cells were washed with FACS buffer and fixed and permeabilized using the eBioscience Foxp3/Transcription Factor Staining Buffer Set (Thermo Fisher Scientific) following manufacturer’s instructions. Intracellular staining was performed overnight in 1× permeabilization buffer. Cells were washed with 1× permeabilization buffer and fixed with 4% formaldehyde solution (MilliporeSigma) in 1× PBS for 10 min. Cells were resuspended in FACS buffer for flow cytometry analysis on a five laser Cytek Aurora spectral flow cytometer. Flow cytometry data were analyzed using FlowJo v10.10.0.

### Human T cell metabolomics

T cells from the exhaustion model were centrifuged at 500 × g, 4 °C, for 2 min and washed with 1 mL ice-cold 0.9% NaCl. Cells were then centrifuged at 500 × g, 4 °C, for 2 min and supernatant was removed. Cell pellets were resuspended in 1 mL of pre-chilled 80% LC-MS grade methanol (Thermo Fisher Scientific), snap-frozen on dry ice, and stored at −80 °C in triplicate. Downstream metabolomics was performed using the same LC-MS method and statistical analysis as described for mouse T cells.

### Human T cell transduction

The plasmid encoding the HLA-A*0201-restricted NY-ESO-1_­_T cell receptor (1G4) has been described previously^64^. Human constant regions of the 1G4 TCR were replaced with codon-optimized murine constant regions as previously described^81^. Codon-optimized sequences encoding the 1G4 TCR (GenScript) were subcloned into the retroviral pMSGV1 vector. For viral packaging, 5 × 10^6^ GP2-293 (Takara) cells were seeded on 100 mm Poly-D-Lysine-coated plates and were co-transfected the following day with 9 μg 1G4-pMSGV1 and 4.5 μg of the envelope plasmid RD114 using Lipofectamine 3000 (Thermo Fisher Scientific). Supernatant was collected on day 2 for viral transduction. Non-tissue culture 24-well plates were coated with RetroNectin (Takara, 20 μg/mL diluted in PBS at 500 μl/well) overnight at 4 °C. The next day, RetroNectin was removed, plates were blocked for 30 min with 2% BSA in PBS, washed once with PBS followed by addition of fresh viral supernatant (1–2 mL/well). Plates were centrifuged at 2000 × g for 2 h at 32 °C. Thereafter, viral supernatant was removed and activated human CD8^+^ T cells (0.5–1 × 10^6^/well, 48 h after stimulation) were added onto the virus-coated plates followed by centrifugation (32 °C for 10 min at 300 × g without brakes). Viral transduction efficiency was assessed by flow cytometry by gating on murine TCRβ^+^. On day 6 of culture, NaCl (Ctrl) or DAHB was added at 20 mM. On day 8, T cells were harvested for adoptive cell therapy and cytotoxicity assays.

### Cytotoxicity of human T cells

For cytotoxicity assays, A375 cells were retrovirally transduced with nuclear-localized mCherry (pMSGV1_mCherry-NLS_P2A_blastR), selected with blasticidin, and single-cell cloned by limiting dilution. 1G4-transduced human CD8^+^ T cells were treated for 48 h with Ctrl or DAHB at 20 mM. An aliquot was restimulated for 3 h with Cell Stimulation Cocktail (1:500) containing the protein transport inhibitors monensin and brefeldin A (Thermo Fisher Scientific) and then analyzed for cytokine production by flow cytometry. The remaining 1G4-transduced human CD8^+^ T cells were co-cultured with A375/mCherry^+^ cells in 96-well plates at varying effector-to-target (E:T) ratios in triplicate wells as previously described^82^. Ctrl and DAHB were maintained at 20 mM throughout the co-culture. Induction of tumor cell apoptosis was quantified using the IncuCyte instrument (Sartorius) with CellEvent Caspase-3/7 Green Detection Reagent (Thermo Fisher Scientific, 1:500 final dilution). The percentage of tumor cell apoptosis was calculated per image as the number of mCherry⁺CellEvent Caspase-3/7 Green⁺ co-positive events (apoptotic tumor cells) divided by the total number of mCherry⁺ events (all tumor cells), equivalent to 100 − %mCherry⁺Caspase-3/7⁻. Tumor cells cultured with metabolites only and untransduced CD8^+^ T cells (Utr) were used as controls.

### *In vivo* xenograft tumor models

HLA-A*0201^+^ A375 melanoma cells expressing NY-ESO-1 were resuspended in PBS and implanted subcutaneously in the backs of NSG mice (1 × 10^6^ cells/mouse) two weeks prior to adoptive cell therapy (ACT). ACT of 1G4-transduced human CD8^+^ T cells (preconditioned with Ctrl or DAHB for 48 h prior to transfer; 1.5 × 10^6^ cells/mouse) occurred by tail vein injection. Starting one day after transfer, mice received daily i.p. injections of NaCl (Ctrl) or DAHB (2.1 g/kg). Tumor area (mm^2^, maximal length × width) was measured daily starting two days post-ACT with calipers. For the no-ACT experiment shown in Figure 7H, tumor-bearing mice did not receive T cells and were treated on the same schedule with daily i.p. injections of PBS, NaCl (Ctrl), or DAHB (2.1 g/kg), and tumor growth was monitored similarly.

### Glucose and lactate measurement

T cells were activated at a density of 200,000 cells/well in 96-well plates overnight followed by treatment with NaCl (Ctrl, 20 mM) or DAHB (20 mM) for 24 h. After treatment, cells were cultured in glucose-free RPMI (MSKCC Media Core) supplemented with dialyzed FBS and 10 mM glucose overnight. Starting media were incubated identically without cells as a baseline control. After 8 h, media glucose consumption and lactate production were measured using the YSI 2900 Analyzer (MSKCC Metabolomics Core) and normalized to cell number.

## QUANTIFICATION AND STATISTICAL ANALYSIS

Statistical analysis was performed using GraphPad Prism (v11.0.0). Mean values are plotted with error bars representing standard deviation. Representative data are shown from at least two independent experiments with similar results and n refers to independent wells per condition unless otherwise indicated. An unpaired two-tailed Student’s t test was used to assess statistical significance between two groups, one-way ANOVA was used to assess statistical significance among three or more groups with one experimental factor, and two-way ANOVA was used to assess statistical significance among three or more groups with two experimental factors. Statistical methods for RNA-seq, metabolomics, and proteomics, are described in their sections. Multiple comparisons corrections were applied as indicated in the figure legends; tumor growth curve significance refers to endpoint tumor area unless otherwise indicated.

## References

1. Frauwirth, K.A., Riley, J.L., Harris, M.H., Parry, R.V., Rathmell, J.C., Plas, D.R., Elstrom, R.L., June, C.H., and Thompson, C.B. (2002). The CD28 Signaling Pathway Regulates Glucose Metabolism. Immunity 16, 769–777. 10.1016/S1074-7613(02)00323-0.

2. Longo, J., Watson, M.J., Williams, K.S., Sheldon, R.D., and Jones, R.G. (2025). Nutrient allocation fuels T cell-mediated immunity. Cell Metab 37, 2311–2322. 10.1016/j.cmet.2025.09.008.

3. Wu, H., Zhao, X., Hochrein, S.M., Eckstein, M., Gubert, G.F., Knöpper, K., Mansilla, A.M., Öner, A., Doucet-Ladevèze, R., Schmitz, W., et al. (2023). Mitochondrial dysfunction promotes the transition of precursor to terminally exhausted T cells through HIF-1α-mediated glycolytic reprogramming. Nat Commun 14, 6858. 10.1038/s41467-023-42634-3.

4. Bailis, W., Shyer, J.A., Zhao, J., Canaveras, J.C.G., Al Khazal, F.J., Qu, R., Steach, H.R., Bielecki, P., Khan, O., Jackson, R., et al. (2019). Distinct modes of mitochondrial metabolism uncouple T cell differentiation and function. Nature 571, 403–407. 10.1038/s41586-019-1311-3.

5. Ikeda, H., Kawase, K., Nishi, T., Watanabe, T., Takenaga, K., Inozume, T., Ishino, T., Aki, S., Lin, J., Kawashima, S., et al. (2025). Immune evasion through mitochondrial transfer in the tumour microenvironment. Nature 638, 225–236. 10.1038/s41586-024-08439-0.

6. Scharping, N.E., Rivadeneira, D.B., Menk, A.V., Vignali, P.D.A., Ford, B.R., Rittenhouse, N.L., Peralta, R., Wang, Y., Wang, Y., DePeaux, K., et al. (2021). Mitochondrial stress induced by continuous stimulation under hypoxia rapidly drives T cell exhaustion. Nat Immunol 22, 205–215. 10.1038/s41590-020-00834-9.

7. Yu, Y.-R., Imrichova, H., Wang, H., Chao, T., Xiao, Z., Gao, M., Rincon-Restrepo, M., Franco, F., Genolet, R., Cheng, W.-C., et al. (2020). Disturbed mitochondrial dynamics in CD8+ TILs reinforce T cell exhaustion. Nat Immunol 21, 1540–1551. 10.1038/s41590-020-0793-3.

8. Vardhana, S.A., Hwee, M.A., Berisa, M., Wells, D.K., Yost, K.E., King, B., Smith, M., Herrera, P.S., Chang, H.Y., Satpathy, A.T., et al. (2020). Impaired mitochondrial oxidative phosphorylation limits the self-renewal of T cells exposed to persistent antigen. Nat Immunol 21, 1022–1033. 10.1038/s41590-020-0725-2.

9. Blagih, J., Coulombe, F., Vincent, E.E., Dupuy, F., Galicia-Vázquez, G., Yurchenko, E., Raissi, T.C., van der Windt, G.J.W., Viollet, B., Pearce, E.L., et al. (2015). The energy sensor AMPK regulates T cell metabolic adaptation and effector responses in vivo. Immunity 42, 41–54. 10.1016/j.immuni.2014.12.030.

10. Costa-Mattioli, M., and Walter, P. (2020). The integrated stress response: From mechanism to disease. Science 368, eaat5314. 10.1126/science.aat5314.

11. Bachem, A., Makhlouf, C., Binger, K.J., de Souza, D.P., Tull, D., Hochheiser, K., Whitney, P.G., Fernandez-Ruiz, D., Dähling, S., Kastenmüller, W., et al. (2019). Microbiota-Derived Short-Chain Fatty Acids Promote the Memory Potential of Antigen-Activated CD8+ T Cells. Immunity 51, 285–297.e5. 10.1016/j.immuni.2019.06.002.

12. Qiu, J., Villa, M., Sanin, D.E., Buck, M.D., O’Sullivan, D., Ching, R., Matsushita, M., Grzes, K.M., Winkler, F., Chang, C.-H., et al. (2019). Acetate Promotes T Cell Effector Function during Glucose Restriction. Cell Rep 27, 2063–2074.e5. 10.1016/j.celrep.2019.04.022.

13. Frisch, A.T., Wang, Y., Xie, B., Yang, A., Ford, B.R., Joshi, S., Kedziora, K.M., Peralta, R., Wilfahrt, D., Mullett, S.J., et al. (2025). Redirecting glucose flux during in vitro expansion generates epigenetically and metabolically superior T cells for cancer immunotherapy. Cell Metab 37, 870–885.e8. 10.1016/j.cmet.2024.12.007.

14. Chaneton, B., Hillmann, P., Zheng, L., Martin, A.C.L., Maddocks, O.D.K., Chokkathukalam, A., Coyle, J.E., Jankevics, A., Holding, F.P., Vousden, K.H., et al. (2012). Serine is a natural ligand and allosteric activator of pyruvate kinase M2. Nature 491, 458–462. 10.1038/nature11540.

15. Pekkala, S., Martínez, A.I., Barcelona, B., Gallego, J., Bendala, E., Yefimenko, I., Rubio, V., and Cervera, J. (2009). Structural insight on the control of urea synthesis: identification of the binding site for N-acetyl-L-glutamate, the essential allosteric activator of mitochondrial carbamoyl phosphate synthetase. Biochem J 424, 211–220. 10.1042/BJ20090888.

16. Schvartzman, J.-M., Reuter, V.P., Koche, R.P., and Thompson, C.B. (2019). 2-hydroxyglutarate inhibits MyoD-mediated differentiation by preventing H3K9 demethylation. Proc Natl Acad Sci U S A 116, 12851–12856. 10.1073/pnas.1817662116.

17. Cai, X., Ng, C.P., Jones, O., Fung, T.S., Ryu, K.W., Li, D., and Thompson, C.B. (2023). Lactate activates the mitochondrial electron transport chain independently of its metabolism. Molecular Cell 83, 3904–3920.e7. 10.1016/j.molcel.2023.09.034.

18. Morovic, P., Gonzalez Moreno, M., Trampuz, A., and Karbysheva, S. (2024). In vitro evaluation of microbial D- and L-lactate production as biomarkers of infection. Front Microbiol 15, 1406350. 10.3389/fmicb.2024.1406350.

19. Remund, B., Yilmaz, B., and Sokollik, C. (2023). D-Lactate: Implications for Gastrointestinal Diseases. Children (Basel) 10, 945. 10.3390/children10060945.

20. Schalkwijk, C.G., and Stehouwer, C.D.A. (2020). Methylglyoxal, a Highly Reactive Dicarbonyl Compound, in Diabetes, Its Vascular Complications, and Other Age-Related Diseases. Physiol Rev 100, 407–461. 10.1152/physrev.00001.2019.

21. Kowlgi, N.G., and Chhabra, L. (2015). D-Lactic Acidosis: An Underrecognized Complication of Short Bowel Syndrome. Gastroenterol Res Pract 2015, 476215. 10.1155/2015/476215.

22. Matelska, D., Shabalin, I.G., Jabłońska, J., Domagalski, M.J., Kutner, J., Ginalski, K., and Minor, W. (2018). Classification, substrate specificity and structural features of D-2-hydroxyacid dehydrogenases: 2HADH knowledgebase. BMC Evol Biol 18, 199. 10.1186/s12862-018-1309-8.

23. Qin, F., Li, J., Mao, T., Feng, S., Li, J., and Lai, M. (2023). 2 Hydroxybutyric Acid-Producing Bacteria in Gut Microbiome and Fusobacterium nucleatum Regulates 2 Hydroxybutyric Acid Level In Vivo. Metabolites 13, 451. 10.3390/metabo13030451.

24. Gao, Y., Bi, D., Xie, R., Li, M., Guo, J., Liu, H., Guo, X., Fang, J., Ding, T., Zhu, H., et al. (2021). Fusobacterium nucleatum enhances the efficacy of PD-L1 blockade in colorectal cancer. Sig Transduct Target Ther 6, 398. 10.1038/s41392-021-00795-x.

25. Wang, X., Fang, Y., Liang, W., Wong, C.C., Qin, H., Gao, Y., Liang, M., Song, L., Zhang, Y., Fan, M., et al. (2024). Fusobacterium nucleatum facilitates anti-PD-1 therapy in microsatellite stable colorectal cancer. Cancer Cell 42, 1729–1746.e8. 10.1016/j.ccell.2024.08.019.

26. Furukawa, N., Miyanaga, A., Nakajima, M., and Taguchi, H. (2018). Structural Basis of Sequential Allosteric Transitions in Tetrameric d-Lactate Dehydrogenases from Three Gram-Negative Bacteria. Biochemistry 57, 5388–5406. 10.1021/acs.biochem.8b00557.

27. Fan, Z., Tang, P., Li, C., Yang, Q., Xu, Y., Su, C., and Li, L. (2023). Fusobacterium nucleatum and its associated systemic diseases: epidemiologic studies and possible mechanisms. J Oral Microbiol 15, 2145729. 10.1080/20002297.2022.2145729.

28. Halestrap, A.P., and Price, N.T. (1999). The proton-linked monocarboxylate transporter (MCT) family: structure, function and regulation. Biochem J 343 *Pt* *2*, 281–299.

29. Kaymak, I., Luda, K.M., Duimstra, L.R., Ma, E.H., Longo, J., Dahabieh, M.S., Faubert, B., Oswald, B.M., Watson, M.J., Kitchen-Goosen, S.M., et al. (2022). Carbon source availability drives nutrient utilization in CD8+ T cells. Cell Metab 34, 1298–1311.e6. 10.1016/j.cmet.2022.07.012.

30. Man, C.H., Mercier, F.E., Liu, N., Dong, W., Stephanopoulos, G., Jiang, L., Jung, Y., Lin, C.P., Leung, A.Y.H., and Scadden, D.T. (2022). Proton export alkalinizes intracellular pH and reprograms carbon metabolism to drive normal and malignant cell growth. Blood 139, 502–522. 10.1182/blood.2021011563.

31. Benjamin, D., Robay, D., Hindupur, S.K., Pohlmann, J., Colombi, M., El-Shemerly, M.Y., Maira, S.-M., Moroni, C., Lane, H.A., and Hall, M.N. (2018). Dual Inhibition of the Lactate Transporters MCT1 and MCT4 Is Synthetic Lethal with Metformin due to NAD+ Depletion in Cancer Cells. Cell Reports 25, 3047–3058.e4. 10.1016/j.celrep.2018.11.043.

32. Bui, D., Ravasz, D., and Chinopoulos, C. (2019). The Effect of 2-Ketobutyrate on Mitochondrial Substrate-Level Phosphorylation. Neurochem Res 44, 2301–2306. 10.1007/s11064-019-02759-8.

33. Jin, S., Chen, X., Yang, J., and Ding, J. (2023). Lactate dehydrogenase D is a general dehydrogenase for D-2-hydroxyacids and is associated with D-lactic acidosis. Nat Commun 14, 6638. 10.1038/s41467-023-42456-3.

34. Shi, L., and Tu, B.P. (2015). Acetyl-CoA and the regulation of metabolism: mechanisms and consequences. Curr Opin Cell Biol 33, 125–131. 10.1016/j.ceb.2015.02.003.

35. Choi, J., Smith, D.M., Lee, Y.J., Cai, D., Hossain, M.J., O’Connor, T.J., Deme, P., Haughey, N.J., Scafidi, S., Riddle, R.C., et al. (2024). Etomoxir repurposed as a promiscuous fatty acid mimetic chemoproteomic probe. iScience 27, 110642. 10.1016/j.isci.2024.110642.

36. Raud, B., Roy, D.G., Divakaruni, A.S., Tarasenko, T.N., Franke, R., Ma, E.H., Samborska, B., Hsieh, W.Y., Wong, A.H., Stüve, P., et al. (2018). Etomoxir Actions on Regulatory and Memory T Cells Are Independent of Cpt1a-Mediated Fatty Acid Oxidation. Cell Metab 28, 504–515.e7. 10.1016/j.cmet.2018.06.002.

37. Conti, R., Mannucci, E., Pessotto, P., Tassoni, E., Carminati, P., Giannessi, F., and Arduini, A. (2011). Selective reversible inhibition of liver carnitine palmitoyl-transferase 1 by teglicar reduces gluconeogenesis and improves glucose homeostasis. Diabetes 60, 644–651. 10.2337/db10-0346.

38. Giannessi, F., Pessotto, P., Tassoni, E., Chiodi, P., Conti, R., De Angelis, F., Dell’Uomo, N., Catini, R., Deias, R., Tinti, M.O., et al. (2003). Discovery of a Long-Chain Carbamoyl Aminocarnitine Derivative, a Reversible Carnitine Palmitoyltransferase Inhibitor with Antiketotic and Antidiabetic Activity. J. Med. Chem. 46, 303–309. 10.1021/jm020979u.

39. Lepez, A., Pirnay, T., Denanglaire, S., Perez-Morga, D., Vermeersch, M., Leo, O., and Andris, F. (2020). Long-term T cell fitness and proliferation is driven by AMPK-dependent regulation of reactive oxygen species. Sci Rep 10, 21673. 10.1038/s41598-020-78715-2.

40. Braverman, E.L., McQuaid, M.A., Schuler, H., Qin, M., Hani, S., Hippen, K., Monlish, D.A., Dobbs, A.K., Ramsey, M.J., Kemp, F., et al. (2024). Overexpression of AMPKγ2 increases AMPK signaling to augment human T cell metabolism and function. Journal of Biological Chemistry 300. 10.1016/j.jbc.2023.105488.

41. Mick, E., Titov, D.V., Skinner, O.S., Sharma, R., Jourdain, A.A., and Mootha, V.K. (2020). Distinct mitochondrial defects trigger the integrated stress response depending on the metabolic state of the cell. eLife 9, e49178. 10.7554/eLife.49178.

42. Tsai, J.C., Miller-Vedam, L.E., Anand, A.A., Jaishankar, P., Nguyen, H.C., Renslo, A.R., Frost, A., and Walter, P. (2018). Structure of the nucleotide exchange factor eIF2B reveals mechanism of memory-enhancing molecule. Science 359, eaaq0939. 10.1126/science.aaq0939.

43. Kazak, L., and Cohen, P. (2020). Creatine metabolism: energy homeostasis, immunity and cancer biology. Nat Rev Endocrinol 16, 421–436. 10.1038/s41574-020-0365-5.

44. Kazak, L., Chouchani, E.T., Jedrychowski, M.P., Erickson, B.K., Shinoda, K., Cohen, P., Vetrivelan, R., Lu, G.Z., Laznik-Bogoslavski, D., Hasenfuss, S.C., et al. (2015). A Creatine-Driven Substrate Cycle Enhances Energy Expenditure and Thermogenesis in Beige Fat. Cell 163, 643–655. 10.1016/j.cell.2015.09.035.

45. Di Biase, S., Ma, X., Wang, X., Yu, J., Wang, Y.-C., Smith, D.J., Zhou, Y., Li, Z., Kim, Y.J., Clarke, N., et al. (2019). Creatine uptake regulates CD8 T cell antitumor immunity. J Exp Med 216, 2869–2882. 10.1084/jem.20182044.

46. Samborska, B., Roy, D.G., Rahbani, J.F., Hussain, M.F., Ma, E.H., Jones, R.G., and Kazak, L. (2022). Creatine transport and creatine kinase activity is required for CD8+ T cell immunity. Cell Rep 38, 110446. 10.1016/j.celrep.2022.110446.

47. Turner, D.C., Wallimann, T., and Eppenberger, H.M. (1973). A protein that binds specifically to the M-line of skeletal muscle is identified as the muscle form of creatine kinase. Proc Natl Acad Sci U S A 70, 702–705. 10.1073/pnas.70.3.702.

48. Bessman, S.P., and Geiger, P.J. (1981). Transport of Energy in Muscle: The Phosphorylcreatine Shuttle. Science 211, 448–452. 10.1126/science.6450446.

49. Billingham, L.K., Stoolman, J.S., Vasan, K., Rodriguez, A.E., Poor, T.A., Szibor, M., Jacobs, H.T., Reczek, C.R., Rashidi, A., Zhang, P., et al. (2022). Mitochondrial electron transport chain is necessary for NLRP3 inflammasome activation. Nat Immunol 23, 692–704. 10.1038/s41590-022-01185-3.

50. Darabedian, N., Ji, W., Fan, M., Lin, S., Seo, H.-S., Vinogradova, E.V., Yaron, T.M., Mills, E.L., Xiao, H., Senkane, K., et al. (2023). Depletion of creatine phosphagen energetics with a covalent creatine kinase inhibitor. Nat Chem Biol 19, 815–824. 10.1038/s41589-023-01273-x.

51. Popović, B., Nicolet, B.P., Guislain, A., Engels, S., Jurgens, A.P., Paravinja, N., Freen-van Heeren, J.J., van Alphen, F.P.J., van den Biggelaar, M., Salerno, F., et al. (2023). Time-dependent regulation of cytokine production by RNA binding proteins defines T cell effector function. Cell Reports 42, 112419. 10.1016/j.celrep.2023.112419.

52. Lattanzio, M.V., Šoštarić, N., Kanagasabesan, N., Popović, B., Bradarić, A., Wardak, L., Guislain, A., Savakis, P., Tutucci, E., and Wolkers, M.C. (2025). Single-molecule imaging of transcription dynamics, RNA localization and fate in human T cells. EMBO J 44, 6732–6749. 10.1038/s44318-025-00592-0.

53. Turgay, Y., Eibauer, M., Goldman, A.E., Shimi, T., Khayat, M., Ben-Harush, K., Dubrovsky-Gaupp, A., Sapra, K.T., Goldman, R.D., and Medalia, O. (2017). The molecular architecture of lamins in somatic cells. Nature 543, 261–264. 10.1038/nature21382.

54. Ren, G., Ku, W.L., Ge, G., Hoffman, J.A., Kang, J.Y., Tang, Q., Cui, K., He, Y., Guan, Y., Gao, B., et al. (2024). Acute depletion of BRG1 reveals its primary function as an activator of transcription. Nat Commun 15, 4561. 10.1038/s41467-024-48911-z.

55. McDonald, B., Chick, B.Y., Ahmed, N.S., Burns, M., Ma, S., Casillas, E., Chen, D., Mann, T.H., O’Connor, C., Hah, N., et al. (2023). Canonical BAF complex activity shapes the enhancer landscape that licenses CD8+ T cell effector and memory fates. Immunity 56, 1303–1319.e5. 10.1016/j.immuni.2023.05.005.

56. He, T., Cheng, C., Qiao, Y., Cho, H., Young, E., Mannan, R., Mahapatra, S., Miner, S.J., Zheng, Y., Kim, N., et al. (2024). Development of an orally bioavailable mSWI/SNF ATPase degrader and acquired mechanisms of resistance in prostate cancer. Proceedings of the National Academy of Sciences 121, e2322563121. 10.1073/pnas.2322563121.

57. Papillon, J.P.N., Nakajima, K., Adair, C.D., Hempel, J., Jouk, A.O., Karki, R.G., Mathieu, S., Möbitz, H., Ntaganda, R., Smith, T., et al. (2018). Discovery of Orally Active Inhibitors of Brahma Homolog (BRM)/SMARCA2 ATPase Activity for the Treatment of Brahma Related Gene 1 (BRG1)/SMARCA4-Mutant Cancers. J. Med. Chem. 61, 10155–10172. 10.1021/acs.jmedchem.8b01318.

58. McPhedran, S.J., Carleton, G.A., and Lum, J.J. (2024). Metabolic engineering for optimized CAR-T cell therapy. Nat Metab 6, 396–408. 10.1038/s42255-024-00976-2.

59. Bacigalupa, Z.A., Landis, M.D., and Rathmell, J.C. (2024). Nutrient inputs and social metabolic control of T cell fate. Cell Metabolism 36, 10–20. 10.1016/j.cmet.2023.12.009.

60. Corrado, M., and Pearce, E.L. (2022). Targeting memory T cell metabolism to improve immunity. J Clin Invest 132. 10.1172/JCI148546.

61. Feng, Q., Liu, Z., Yu, X., Huang, T., Chen, J., Wang, J., Wilhelm, J., Li, S., Song, J., Li, W., et al. (2022). Lactate increases stemness of CD8 + T cells to augment anti-tumor immunity. Nat Commun 13, 4981. 10.1038/s41467-022-32521-8.

62. Belk, J.A., Yao, W., Ly, N., Freitas, K.A., Chen, Y.-T., Shi, Q., Valencia, A.M., Shifrut, E., Kale, N., Yost, K.E., et al. (2022). Genome-wide CRISPR screens of T cell exhaustion identify chromatin remodeling factors that limit T cell persistence. Cancer Cell 40, 768–786.e7. 10.1016/j.ccell.2022.06.001.

63. Chen, Z., Olszewski, K.L., Ryseck, R.-P., Xu, X., Boyer, J.A., Peace, C.G., Shen, Y., Bartman, C.R., Lynch, L., and Rabinowitz, J.D. (2025). Oxidative pentose phosphate pathway is required for T cell activation and antitumor immunity. Proc Natl Acad Sci U S A 122, e2516288122. 10.1073/pnas.2516288122.

64. Robbins, P.F., Li, Y.F., El-Gamil, M., Zhao, Y., Wargo, J.A., Zheng, Z., Xu, H., Morgan, R.A., Feldman, S.A., Johnson, L.A., et al. (2008). Single and dual amino acid substitutions in TCR CDRs can enhance antigen-specific T cell functions. J Immunol 180, 6116–6131. 10.4049/jimmunol.180.9.6116.

65. Furukawa, N., Miyanaga, A., Togawa, M., Nakajima, M., and Taguchi, H. (2014). Diverse allosteric and catalytic functions of tetrameric d-lactate dehydrogenases from three Gram-negative bacteria. AMB Express 4, 76. 10.1186/s13568-014-0076-1.

66. Simula, L., Fumagalli, M., Vimeux, L., Rajnpreht, I., Icard, P., Birsen, G., An, D., Pendino, F., Rouault, A., Bercovici, N., et al. (2024). Mitochondrial metabolism sustains CD8+ T cell migration for an efficient infiltration into solid tumors. Nat Commun 15, 2203. 10.1038/s41467-024-46377-7.

67. Zhang, Y., Kurupati, R., Liu, L., Zhou, X.Y., Zhang, G., Hudaihed, A., Filisio, F., Giles-Davis, W., Xu, X., Karakousis, G.C., et al. (2017). Enhancing CD8+ T Cell Fatty Acid Catabolism within a Metabolically Challenging Tumor Microenvironment Increases the Efficacy of Melanoma Immunotherapy. Cancer Cell 32, 377–391.e9. 10.1016/j.ccell.2017.08.004.

68. Chowdhury, P.S., Chamoto, K., Kumar, A., and Honjo, T. (2018). PPAR-Induced Fatty Acid Oxidation in T Cells Increases the Number of Tumor-Reactive CD8+ T Cells and Facilitates Anti-PD-1 Therapy. Cancer Immunol Res 6, 1375–1387. 10.1158/2326-6066.CIR-18-0095.

69. Xu, S., Chaudhary, O., Rodríguez-Morales, P., Sun, X., Chen, D., Zappasodi, R., Xu, Z., Pinto, A.F.M., Williams, A., Schulze, I., et al. (2021). Uptake of oxidized lipids by the scavenger receptor CD36 promotes lipid peroxidation and dysfunction in CD8+ T cells in tumors. Immunity 54, 1561–1577.e7. 10.1016/j.immuni.2021.05.003.

70. Ma, X., Xiao, L., Liu, L., Ye, L., Su, P., Bi, E., Wang, Q., Yang, M., Qian, J., and Yi, Q. (2021). CD36-mediated ferroptosis dampens intratumoral CD8+ T cell effector function and impairs their antitumor ability. Cell Metab 33, 1001–1012.e5. 10.1016/j.cmet.2021.02.015.

71. Yi, Y., Luo, C., Wang, J., Wang, Y., Jin, L., Sun, X., Hao, J., Deng, G., Jin, R., and Ge, Q. (2026). Basal level of ATF4 promotes T cell readiness for activation-induced proliferation. iScience 29, 114277. 10.1016/j.isci.2025.114277.

72. Ramgopal, A., Braverman, E.L., Sun, L.-K., Monlish, D., Wittmann, C., Kemp, F., Qin, M., Ramsey, M.J., Cattley, R., Hawse, W., et al. (2024). AMPK drives both glycolytic and oxidative metabolism in murine and human T cells during graft-versus-host disease. Blood Adv 8, 4149–4162. 10.1182/bloodadvances.2023010740.

73. Wallimann, T., Tokarska-Schlattner, M., and Schlattner, U. (2011). The creatine kinase system and pleiotropic effects of creatine. Amino Acids 40, 1271–1296. 10.1007/s00726-011-0877-3.

74. Battistello, E., Hixon, K.A., Comstock, D.E., Collings, C.K., Chen, X., Rodriguez Hernaez, J., Lee, S., Cervantes, K.S., Hinkley, M.M., Ntatsoulis, K., et al. (2023). Stepwise activities of mSWI/SNF family chromatin remodeling complexes direct T cell activation and exhaustion. Mol Cell 83, 1216–1236.e12. 10.1016/j.molcel.2023.02.026.

75. Guo, A., Huang, H., Zhu, Z., Chen, M.J., Shi, H., Yuan, S., Sharma, P., Connelly, J.P., Liedmann, S., Dhungana, Y., et al. (2022). cBAF complex components and MYC cooperate early in CD8+ T cell fate. Nature 607, 135–141. 10.1038/s41586-022-04849-0.

76. Kojima, H., Wayne, C.R., Patterson, L.F.S., Sanford, H., Chen, T.-J., Lin, Y.-H., Schoenfeld, J.D., McGary, L.H.F., Chen, Y.-T., Kropp, K.N., et al. (2026). A post-translational regulatory map of chronic antigen-driven human T cell dysfunction. Preprint at bioRxiv, 10.64898/2026.03.04.709614.

77. Evans, D.A., Morrissey, M.M., and Dorow, R.L. (1985). Asymmetric oxygenation of chiral imide enolates. A general approach to the synthesis of enantiomerically pure .alpha.-hydroxy carboxylic acid synthons. J. Am. Chem. Soc. 107, 4346–4348. 10.1021/ja00300a054.

78. Steele, A.D., Knouse, K.W., Keohane, C.E., and Wuest, W.M. (2015). Total Synthesis and Biological Investigation of (−)-Promysalin. J. Am. Chem. Soc. 137, 7314–7317. 10.1021/jacs.5b04767.

79. Livak, K.J., and Schmittgen, T.D. (2001). Analysis of relative gene expression data using real-time quantitative PCR and the 2(-Delta Delta C(T)) Method. Methods 25, 402–408. 10.1006/meth.2001.1262.

80. The Galaxy Community (2024). The Galaxy platform for accessible, reproducible, and collaborative data analyses: 2024 update. Nucleic Acids Research 52, W83–W94. 10.1093/nar/gkae410.

81. Cohen, C.J., Zhao, Y., Zheng, Z., Rosenberg, S.A., and Morgan, R.A. (2006). Enhanced antitumor activity of murine-human hybrid T-cell receptor (TCR) in human lymphocytes is associated with improved pairing and TCR/CD3 stability. Cancer Res 66, 8878–8886. 10.1158/0008-5472.CAN-06-1450.

82. Carnevale, J., Shifrut, E., Kale, N., Nyberg, W.A., Blaeschke, F., Chen, Y.Y., Li, Z., Bapat, S.P., Diolaiti, M.E., O’Leary, P., et al. (2022). RASA2 ablation in T cells boosts antigen sensitivity and long-term function. Nature 609, 174–182. 10.1038/s41586-022-05126-w.

