## Supplemental for "Mitochondrial ATP production promotes T cell differentiation and function by regulating chromatin accessibility"

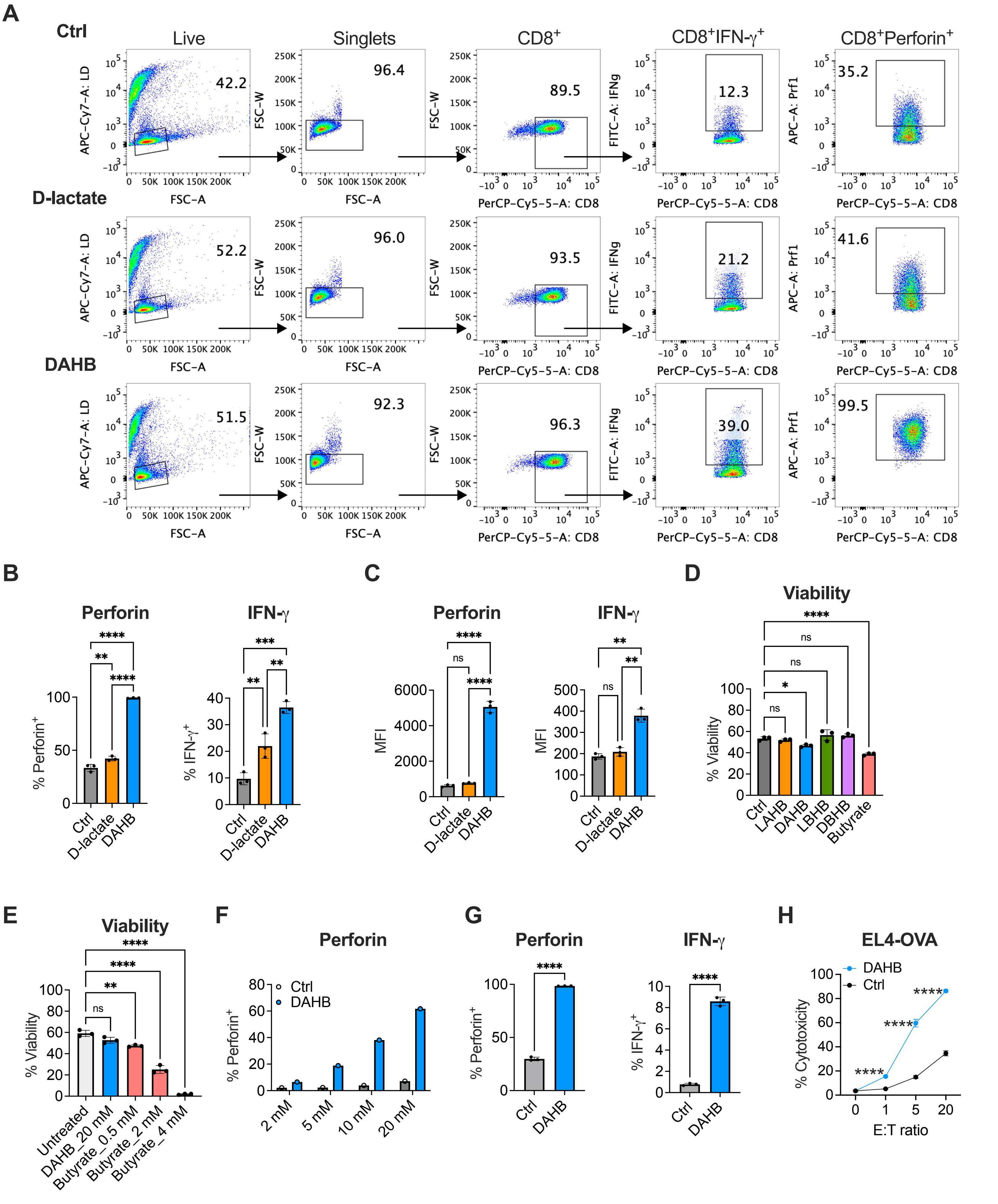

A
Ctrl
D-lactate
DAHB
Live
Singlets
CD8+
CD8+IFN-γ+
CD8+Perforin+
42.2
96.4
89.5
12.3
35.2
52.2
96.0
93.5
21.2
41.6
51.5
92.3
96.3
39.0
99.5
APC-Cy7-A: LD
FSC-A
FSC-W
PerCP-Cy5-5-A: CD8
FITC-A: IFNg
APC-A: Prf1
B
Perforin
IFN-γ
% Perforin+
% IFN-γ+
Ctrl
D-lactate
DAHB
C
MFI
D
Viability
% Viability
Ctrl
LAHB
DAHB
LBHB
DBHB
Butyrate
ns
E
Untreated
DAHB_20 mM
Butyrate_0.5 mM
Butyrate_2 mM
Butyrate_4 mM
F
2 mM
5 mM
10 mM
20 mM
G
H
EL4-OVA
% Cytotoxicity
E:T ratio


Figure S1

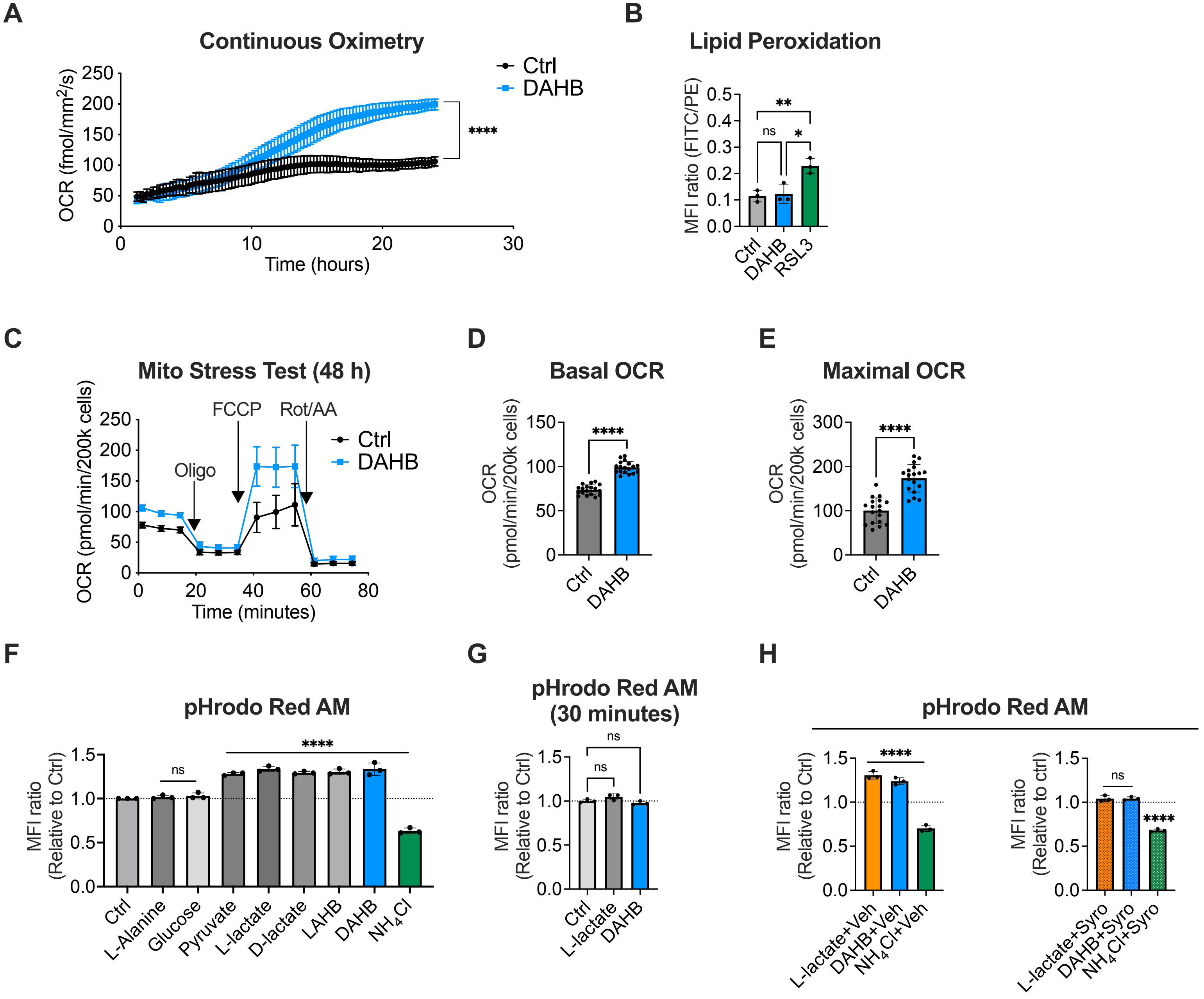
A
Continuous Oximetry
OCR (fmol/mm²/s)
Time (hours)
Ctrl
DAHB
****
B
Lipid Peroxidation
MFI ratio (FITC/PE)
**
ns
*
Ctrl
DAHB
RSL3
C
Mito Stress Test (48 h)
OCR (pmol/min/200k cells)
Oligo
FCCP
Rot/AA
Time (minutes)
D
Basal OCR
OCR (pmol/min/200k cells)
E
Maximal OCR
F
pHrodo Red AM
MFI ratio (Relative to Ctrl)
Ctrl
L-Alanine
Glucose
Pyruvate
L-lactate
D-lactate
LAHB
DAHB
NH4Cl
G
pHrodo Red AM (30 minutes)
H
pHrodo Red AM
L-lactate+Veh
DAHB+Veh
NH4Cl+Veh
MFI ratio (Relative to ctrl)
L-lactate+Syro
DAHB+Syro
NH4Cl+Syro

Figure S2

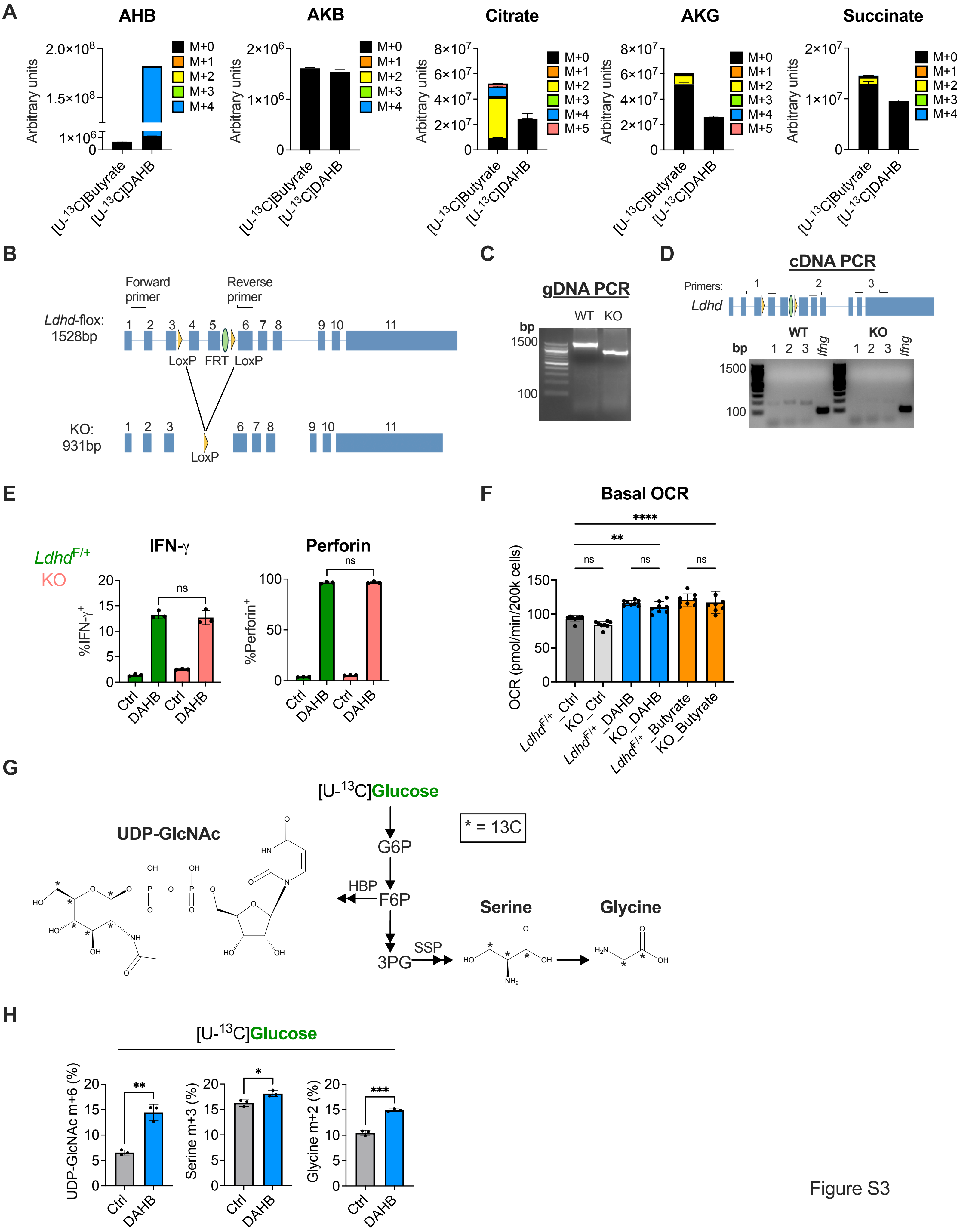
A
AHB
AKB
Citrate
AKG
Succinate
Arbitrary units
M+0
M+1
M+2
M+3
M+4
M+5
[U-13C]Butyrate
[U-13C]DAHB
B
Forward primer
Reverse primer
Ldhd-flox: 1528bp
LoxP FRT LoxP
KO: 931bp
LoxP
C
gDNA PCR
WT KO
bp
1500
100
D
cDNA PCR
Primers:
Ldhd
WT
KO
Ifng
E
LdhdF/+
KO
IFN-γ
Perforin
%IFN-γ+
%Perforin+
Ctrl
DAHB
ns
F
Basal OCR
OCR (pmol/min/200k cells)
LdhdF/+_Ctrl
KO_Ctrl
LdhdF/+_DAHB
KO_DAHB
LdhdF/+_Butyrate
KO_Butyrate
G
[U-13C]Glucose
UDP-GlcNAc
* = 13C
G6P
F6P
HBP
3PG
SSP
Serine
Glycine
H
UDP-GlcNAc m+6 (%)
Serine m+3 (%)
Glycine m+2 (%)

Figure S3

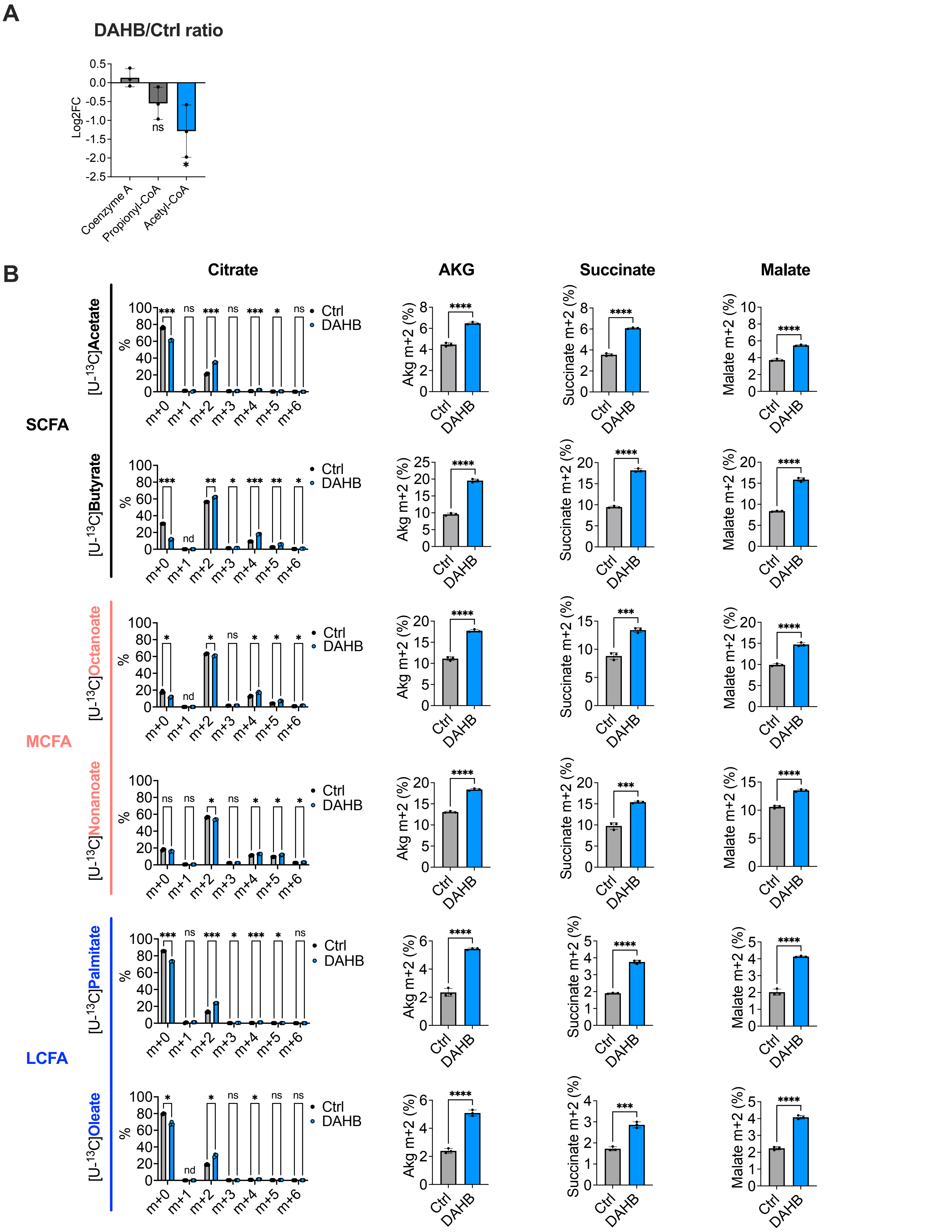
A
DAHB/Ctrl ratio
Log2FC
0.5
0.0
-0.5
-1.0
-1.5
-2.0
-2.5
ns
*
Coenzyme A
Propionyl-CoA
Acetyl-CoA
B
Citrate
AKG
Succinate
Malate
SCFA
MCFA
LCFA
[U-13C]Acetate
[U-13C]Butyrate
[U-13C]Octanoate
[U-13C]Nonanoate
[U-13C]Palmitate
[U-13C]Oleate
%
m+0
m+1
m+2
m+3
m+4
m+5
m+6
Ctrl
DAHB
nd
ns
*
**
***
****
Akg m+2 (%)
Succinate m+2 (%)
Malate m+2 (%)

Figure S4

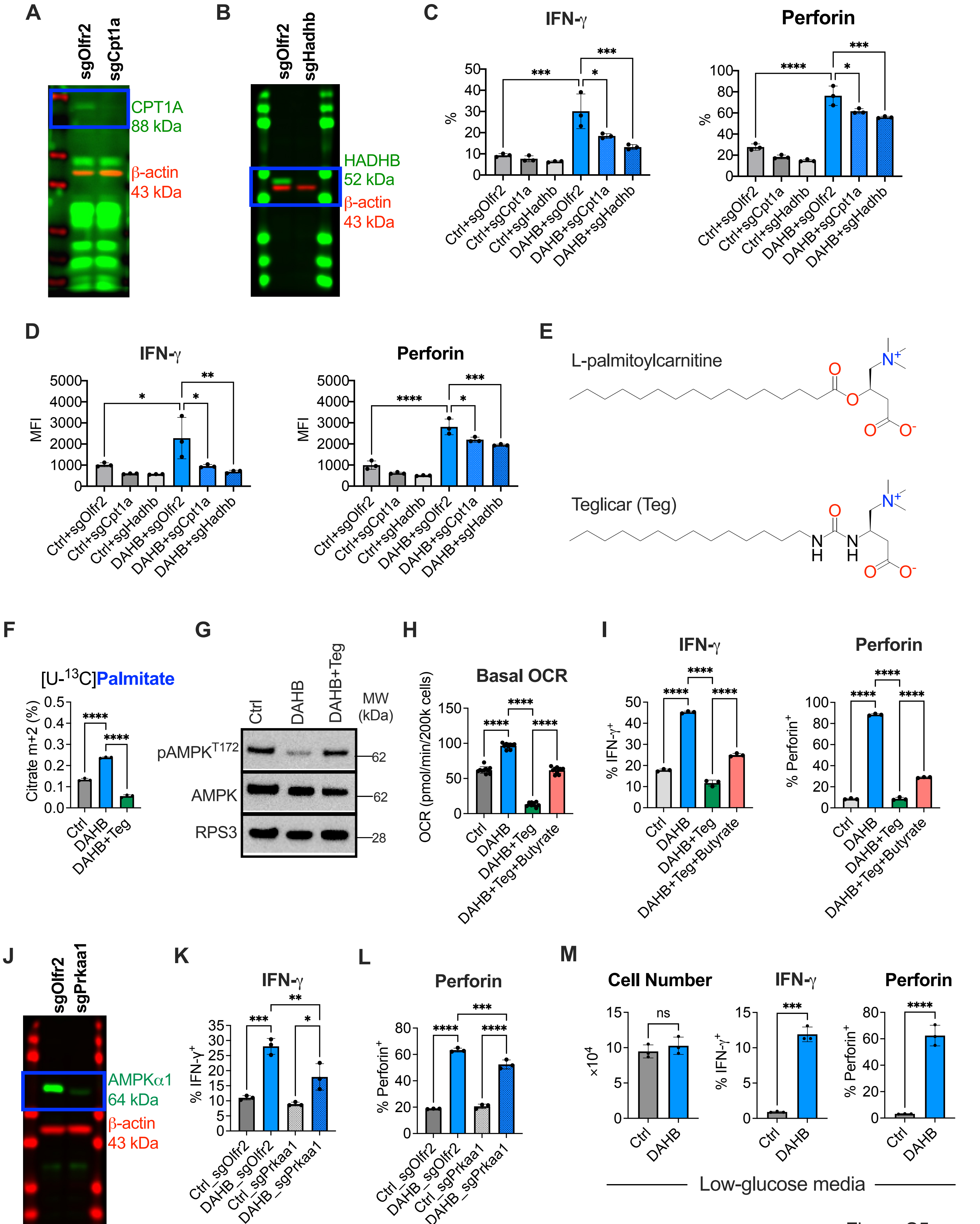

A
sgOlfr2
sgCpt1a
CPT1A
88 kDa
β-actin
43 kDa
B
sgOlfr2
sgHadhb
HADHB
52 kDa
β-actin
43 kDa
C
IFN-γ
Perforin
%
Ctrl+sgOlfr2
Ctrl+sgCpt1a
Ctrl+sgHadhb
DAHB+sgOlfr2
DAHB+sgCpt1a
DAHB+sgHadhb
D
IFN-γ
Perforin
MFI
E
L-palmitoylcarnitine
Teglicar (Teg)
F
[U-13C]Palmitate
Citrate m+2 (%)
Ctrl
DAHB
DAHB+Teg
G
Ctrl
DAHB
DAHB+Teg
MW (kDa)
pAMPKT172
AMPK
RPS3
62
62
28
H
Basal OCR
OCR (pmol/min/200k cells)
DAHB+Teg+Butyrate
I
IFN-γ
Perforin
% IFN-γ+
% Perforin+
J
sgOlfr2
sgPrkaa1
AMPKα1
64 kDa
β-actin
43 kDa
K
IFN-γ
% IFN-γ+
Ctrl_sgOlfr2
DAHB_sgOlfr2
Ctrl_sgPrkaa1
DAHB_sgPrkaa1
L
Perforin
% Perforin+
M
Cell Number
IFN-γ
Perforin
×10^4
% IFN-γ+
% Perforin+
ns
Low-glucose media


Figure S5

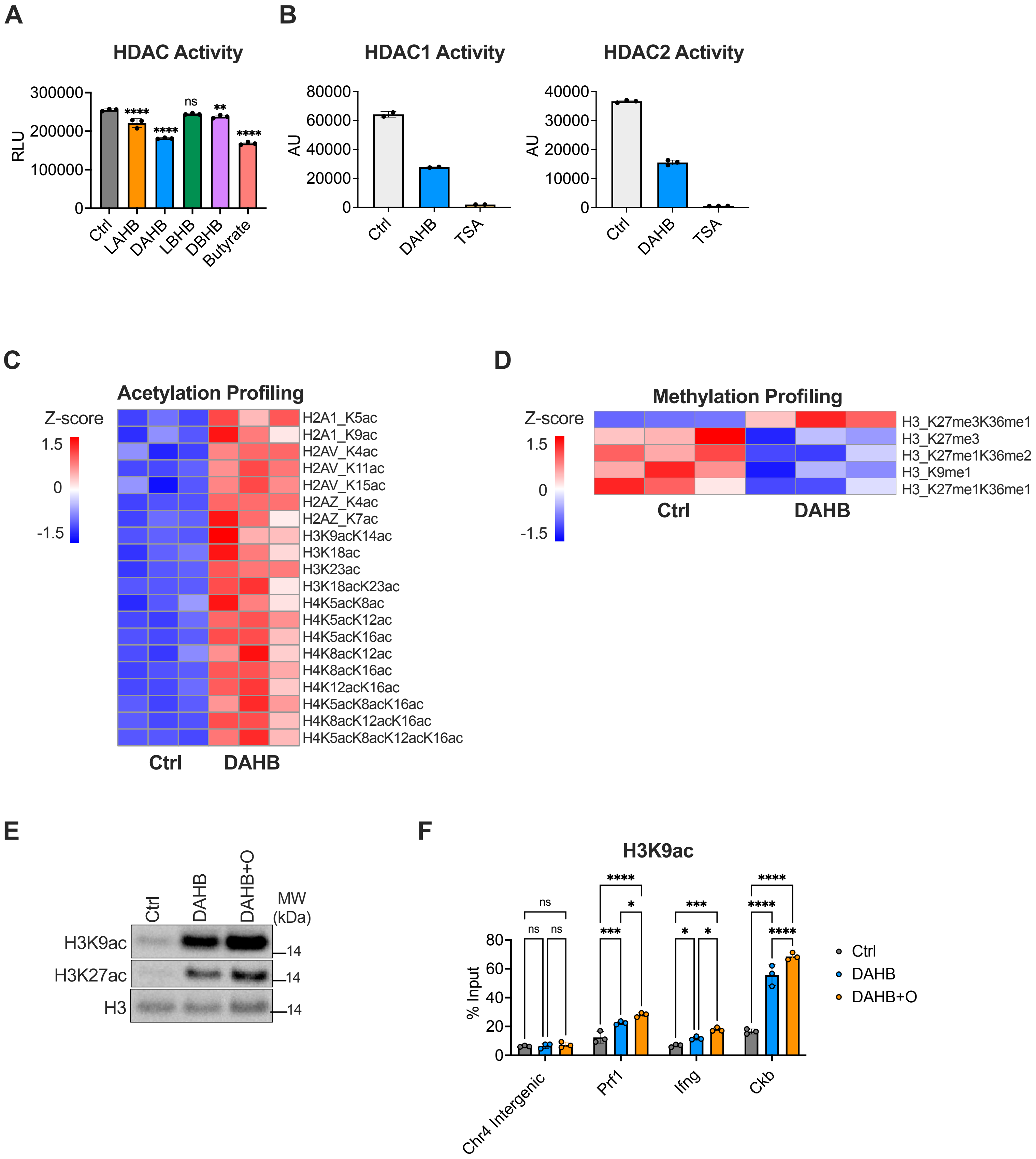
A
HDAC Activity
RLU
300000
200000
100000
0
Ctrl
LAHB
DAHB
LBHB
DBHB
Butyrate
****
****
ns
**
****
B
HDAC1 Activity
AU
80000
60000
40000
20000
0
Ctrl
DAHB
TSA
HDAC2 Activity
AU
40000
30000
20000
10000
0
C
Acetylation Profiling
Z-score
1.5
0
-1.5
H2A1_K5ac
H2A1_K9ac
H2AV_K4ac
H2AV_K11ac
H2AV_K15ac
H2AZ_K4ac
H2AZ_K7ac
H3K9acK14ac
H3K18ac
H3K23ac
H3K18acK23ac
H4K5acK8ac
H4K5acK12ac
H4K5acK16ac
H4K8acK12ac
H4K8acK16ac
H4K12acK16ac
H4K5acK8acK16ac
H4K8acK12acK16ac
H4K5acK8acK12acK16ac
Ctrl
DAHB
D
Methylation Profiling
H3_K27me3K36me1
H3_K27me3
H3_K27me1K36me2
H3_K9me1
H3_K27me1K36me1
E
Ctrl
DAHB
DAHB+O
MW (kDa)
H3K9ac
H3K27ac
H3
14
F
H3K9ac
% Input
80
60
40
20
0
Chr4 Intergenic
Prf1
Ifng
Ckb
ns
***
*
****
Ctrl
DAHB
DAHB+O

Figure S6

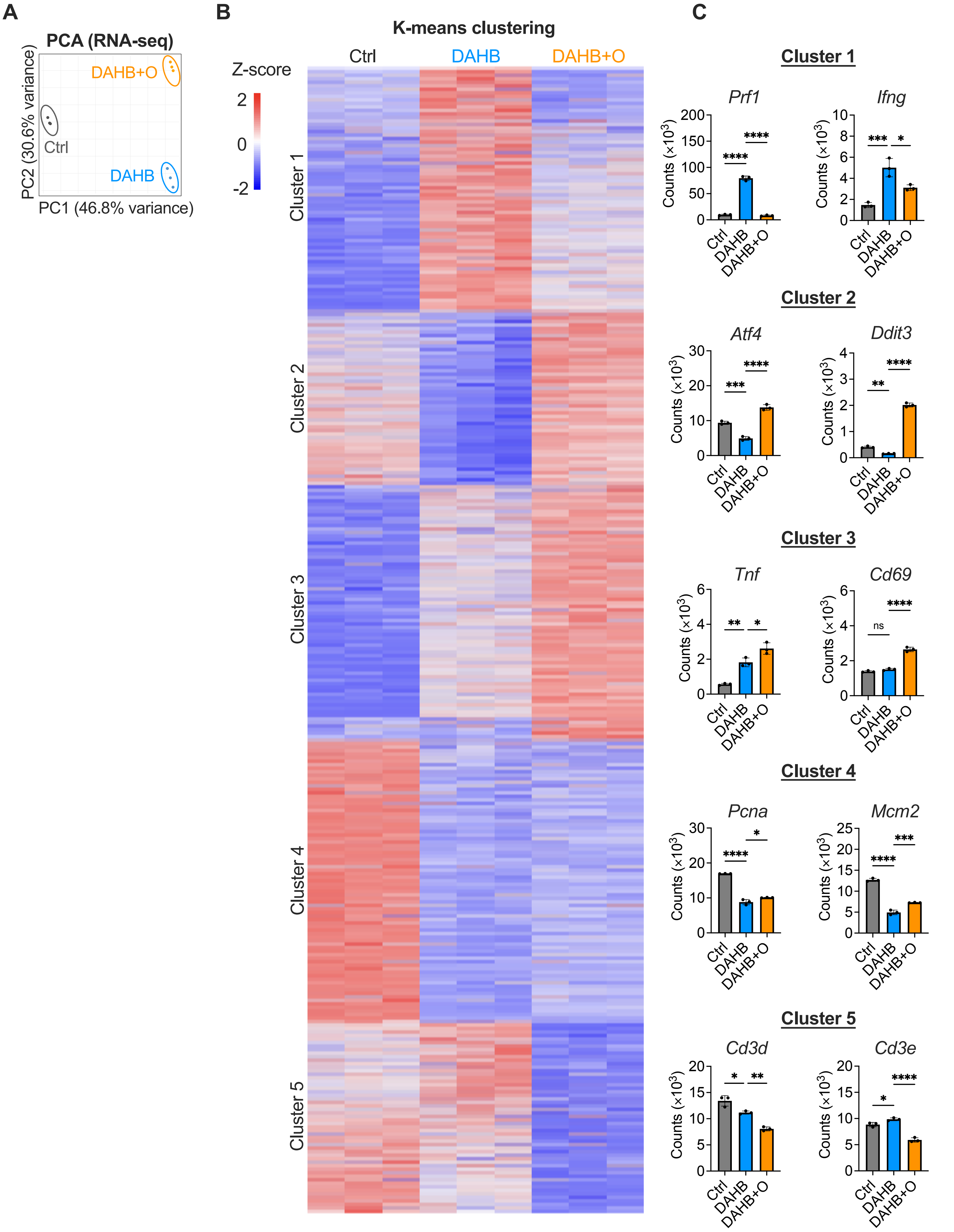
A
PCA (RNA-seq)
DAHB+O
Ctrl
DAHB
PC2 (30.6% variance)
PC1 (46.8% variance)
B
K-means clustering
Z-score
2
0
-2
Ctrl
DAHB
DAHB+O
Cluster 1
Cluster 2
Cluster 3
Cluster 4
Cluster 5
C
Cluster 1
Prf1
Ifng
Cluster 2
Atf4
Ddit3
Cluster 3
Tnf
Cd69
Cluster 4
Pcna
Mcm2
Cluster 5
Cd3d
Cd3e
Counts (×10^3)
Ctrl
DAHB
DAHB+O
ns

Figure S7

**A**

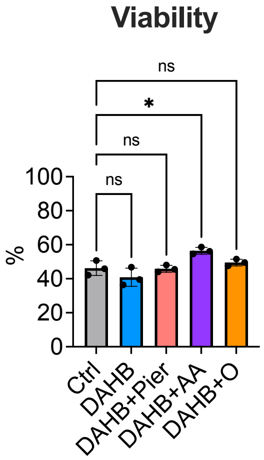
Viability
ns
*
ns
ns
100
80
60
40
20
0
%
Ctrl
DAHB
DAHB+Pier
DAHB+AA
DAHB+O

**B**

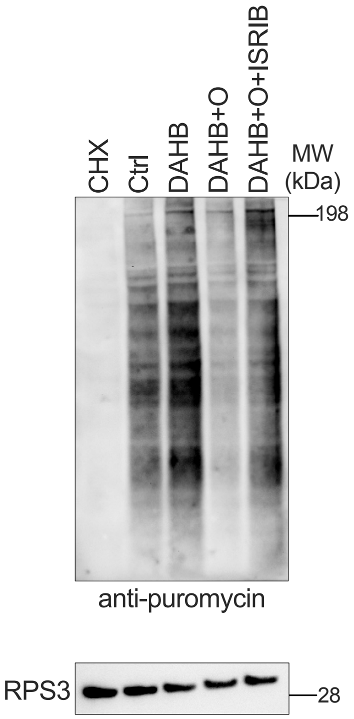
CHX
Ctrl
DAHB
DAHB+O
DAHB+O+ISRIB
MW (kDa)
198
anti-puromycin
RPS3
28

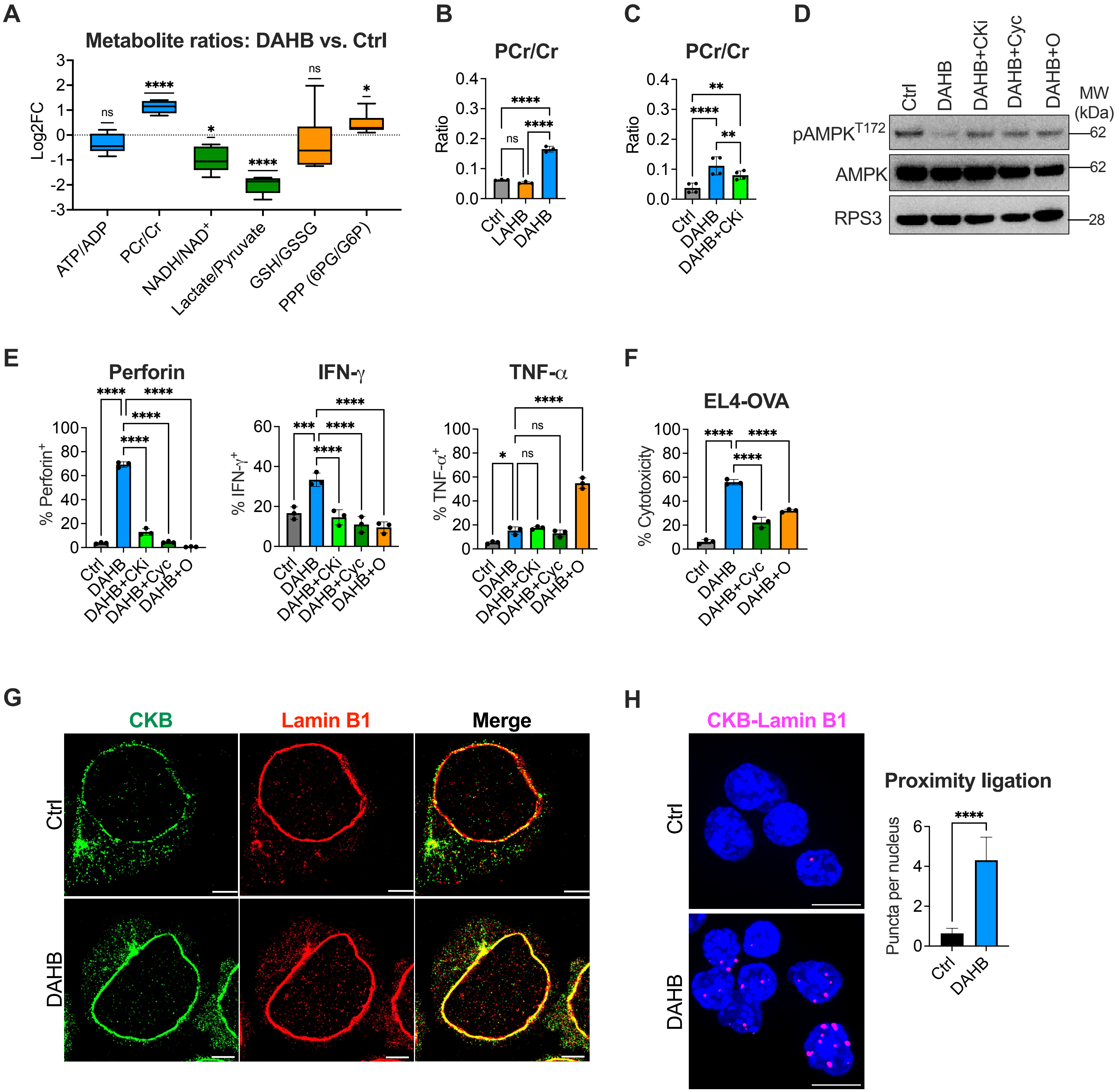
A
Metabolite ratios: DAHB vs. Ctrl
Log2FC
ATP/ADP
PCr/Cr
NADH/NAD+
Lactate/Pyruvate
GSH/GSSG
PPP (6PG/G6P)
B
PCr/Cr
Ratio
Ctrl
LAHB
DAHB
C
PCr/Cr
Ratio
Ctrl
DAHB
DAHB+CKi
D
Ctrl
DAHB
DAHB+CKi
DAHB+Cyc
DAHB+O
MW (kDa)
pAMPKT172
AMPK
RPS3
E
Perforin
% Perforin+
IFN-γ
% IFN-γ+
TNF-α
% TNF-α+
Ctrl
DAHB
DAHB+CKi
DAHB+Cyc
DAHB+O
F
EL4-OVA
% Cytotoxicity
G
CKB
Lamin B1
Merge
Ctrl
DAHB
H
CKB-Lamin B1
Proximity ligation
Puncta per nucleus
Ctrl
DAHB

Figure S9

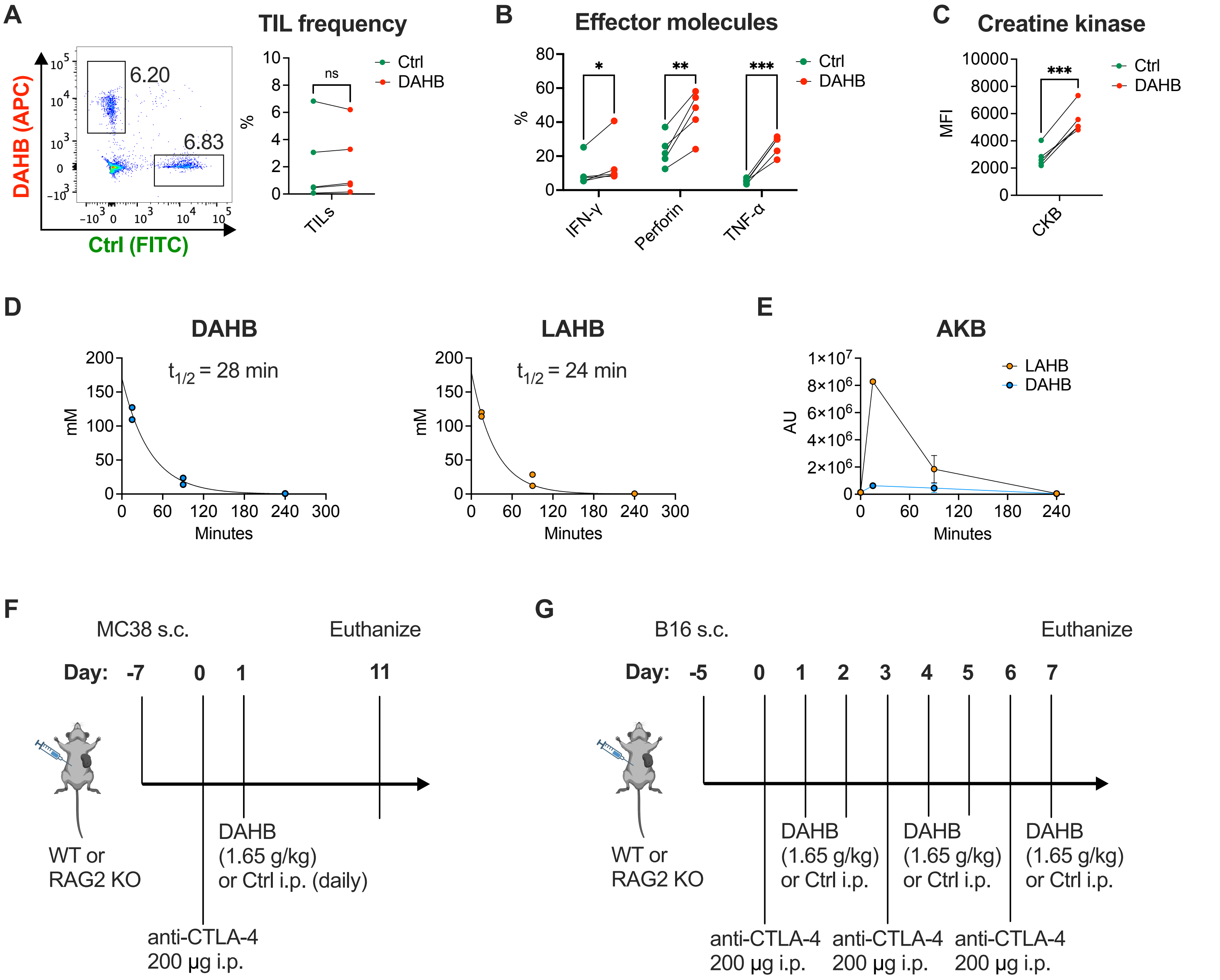

A
TIL frequency
DAHB (APC)
Ctrl (FITC)
6.20
6.83
ns
%
TILs
Ctrl
DAHB
B
Effector molecules
*
**
***
IFN-γ
Perforin
TNF-α
C
Creatine kinase
MFI
CKB
D
DAHB
$t_{1/2}$ = 28 min
mM
Minutes
LAHB
$t_{1/2}$ = 24 min
E
AKB
AU
LAHB
DAHB
F
MC38 s.c.
Euthanize
Day: -7 0 1 11
WT or
RAG2 KO
DAHB
(1.65 g/kg)
or Ctrl i.p. (daily)
anti-CTLA-4
200 µg i.p.
G
B16 s.c.
Euthanize
Day: -5 0 1 2 3 4 5 6 7
WT or
RAG2 KO
DAHB
(1.65 g/kg)
or Ctrl i.p.
anti-CTLA-4
200 µg i.p.


Figure S10

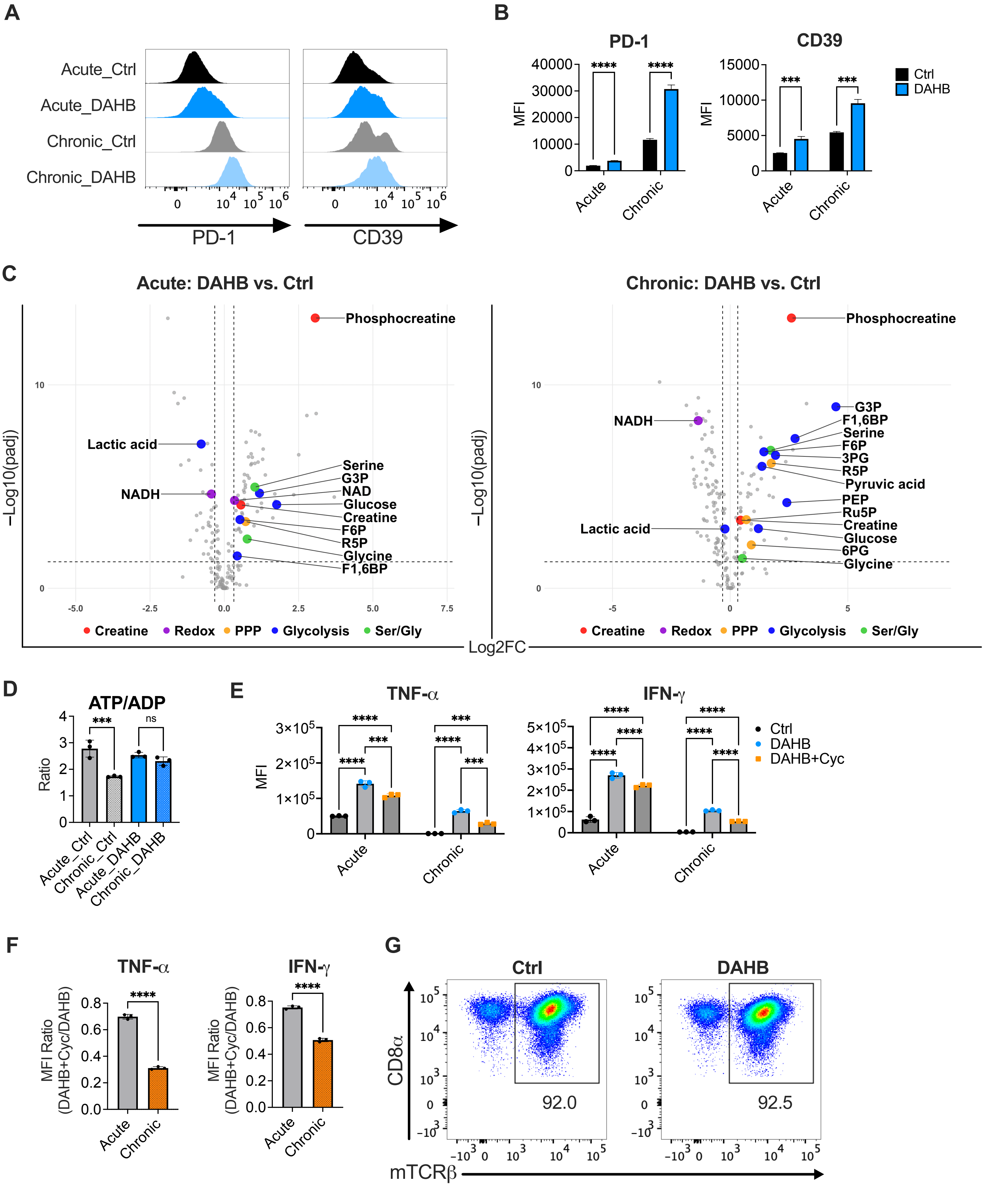

A
Acute_Ctrl
Acute_DAHB
Chronic_Ctrl
Chronic_DAHB
PD-1
CD39
B
PD-1
CD39
MFI
Ctrl
DAHB
Acute
Chronic
C
Acute: DAHB vs. Ctrl
Chronic: DAHB vs. Ctrl
Phosphocreatine
Lactic acid
NADH
Serine
G3P
NAD
Glucose
Creatine
F6P
R5P
Glycine
F1,6BP
3PG
Pyruvic acid
PEP
Ru5P
6PG
−Log10(padj)
Creatine
Redox
PPP
Glycolysis
Ser/Gly
Log2FC
D
ATP/ADP
Ratio
Acute_Ctrl
Chronic_Ctrl
Acute_DAHB
Chronic_DAHB
E
TNF-α
IFN-γ
MFI
Ctrl
DAHB
DAHB+Cyc
F
TNF-α
IFN-γ
MFI Ratio (DAHB+Cyc/DAHB)
G
Ctrl
DAHB
CD8α
mTCRβ
92.0
92.5


Figure S11
